# Deciphering deep-sea chemosynthetic symbiosis by single-nucleus RNA-sequencing

**DOI:** 10.1101/2023.05.29.542712

**Authors:** Hao Wang, Kai He, Huan Zhang, Quanyong Zhang, Lei Cao, Jing Li, Zhaoshan Zhong, Hao Chen, Li Zhou, Chao Lian, Minxiao Wang, Kai Chen, Pei-Yuan Qian, Chaolun Li

## Abstract

Bathymodioline mussels dominate deep-sea methane seep and hydrothermal vent habitats and obtain nutrients and energy primarily through chemosynthetic endosymbiotic bacteria in the bacteriocytes of their gill. However, the molecular mechanisms that orchestrate mussel host-symbiont interactions remain unclear. Here, we constructed a comprehensive cell atlas of the gill in the mussel *Gigantidas platifrons* from the South China Sea methane seeps (1100m depth) using single-nucleus RNA sequencing (snRNA-seq) and whole-mount *in situ* hybridisation. We identified 13 types of cells, including three previously unknown ones, and uncovered unknown tissue heterogeneity. Every cell type has a designated function in supporting the gill’s structure and function, creating an optimal environment for chemosynthesis, and effectively acquiring nutrients from the endosymbiotic bacteria. Analysis of snRNA-seq of *in situ* transplanted mussels clearly showed the shifts in cell state in response to environmental oscillations. Our findings provide insight into principles of host-symbiont interaction and the bivalves’ environmental adaption mechanisms.

## Introduction

Mutualistic interactions between multicellular animals and their microbiota play a fundamental role in animals’ adaptation, ecology, and evolution ^1,2^. By associating with symbionts, host animals benefit from the metabolic capabilities of their symbionts and gain fitness advantages that allow them to thrive in habitats they could not live in on their own^1,3^. Prime examples of such symbioses are bathymodioline mussels and gammaproteobacteria endosymbionts^4^. The bathymodioline mussels occur worldwide at deep-sea chemosynthetic ecosystems, such as cold-seeps, hydrothermal vents, and whale falls^4^. The host mussel acquired sulfur-oxidising (SOX) and/or methane-oxidising (MOX) symbiont through horizontal transmission at their early life stage^3^. Their symbionts, which are hosted in a specialised gill epithelial cell, namely, the bacteriocytes^5,6^, utilise the chemical energy from the reduced chemical compounds such as CH_4_ and H_2_S released from cold seeps or hydrothermal vents to fix carbon and turn into carbon source for the host mussel^7,8^. The ecological success of the bathymodioline symbioses is apparent: the bathymodiolin mussels are often among the most dominant species in the deep-sea chemosynthetic ecosystems^9,10^. Thus, it is critical to know how the bathymodioline mussels interact with the symbionts and maintain the stability and efficiency of the symbiosis.

The gill structure of bathymodiolin mussels has undergone remarkable adaptations at the molecular, cellular, and tissue levels to support their deep-sea chemosynthetic lifestyle^11,12^. Compared to shallow water mussels^13^, the gill filaments of bathymodiolin mussels have enlarged surfaces allowing them to hold more symbionts per filament (Supplementary Fig. S1). This adaption requires not only novel cellular and molecular mechanisms to maintain and support the enlarged gill filament structure, but also a strong ciliary ventilation system to pump vent/seep fluid to fuel the symbiotic bacteria. Previous studies based on whole-genome sequencing and bulk RNA-seq projects have shown that genes in the categories of nutrient transporters, lysosomal proteins, and immune receptors are either expanded in the host mussel’s genome or up-regulated in the gill^12, 14–17^, suggesting that these genes involved in host-symbiont interaction. Though providing deep molecular insights, these studies mainly used homogenised tissues that average genes’ expression levels amongst different cell types and eliminate cell and gene spatial distribution information^18–21^. In addition, the broad expression and function of potentially ‘symbiosis-related’ proteins also greatly limited data interpretation. Therefore, a systemic atlas of gill cell types and the descriptions of cell-type-specific gene expressional profiles are warranted to a better understanding of the host-symbiont interaction and environmental adaptation mechanisms of the bathymodioline symbiosis.

In recent years, single cell/nucleus RNA-sequencing (sc/sn RNA-seq) technologies have become one of the preferred methods for investigating the composition of complex tissue at the transcriptional level^22^. snRNA-seq has several advantages, such as compatibility with frozen samples, elimination of dissociation-induced transcriptional stress responses, and reduced dissociation bias^23^. These advantages are significant for deep-sea and other ecological studies because cell and molecular biology facilities are not available in the field. To examine the cellular and molecular mechanisms of the environmental adaptations and host-symbiont interactions in bathymodioline mussels, we conducted a single nucleus RNA-sequencing-based transcriptomic study. We analysed the gill symbiosis in the dominate deep-sea mussels inhabiting the F-site cold-seep *Gigantidas platifrons*^24^, which hosts a single a MOX endosymbiont population that consist of only one 16S rRNA phylotype. Our work provides a proof-of-principle for environmental adaptation mechanisms study in non-traditional but ecologically important organisms with single-nucleus RNA-sequencing technologies.

## Results and discussion

### 1. *G. platifrons* deep-sea *in situ* transplant experiment and single-cell transcriptomic sequencing

We conducted a *G. platifrons* deep-sea *in situ* transplant experiment at the “F-Site” Cold-seep (∼1117 m depth) and retrieved three groups of samples (Fig. 1A), as follows: the ‘Fanmao’ (meaning prosperous) group, which comprised the mussels collected from methane-rich (∼40 μM) site of the cold-seep, where the animals thrived; the ‘starvation’ group, which comprised the mussels collected from the methane-rich site and then moved to a methane-limited (∼0.054 μM) starvation site for 14 days before retrieval; and the ‘reconstitution’ group, which comprised the methane-rich site mussels moved to the ‘Starvation’ site for 11 days and then moved back to the methane-rich site for another 3 days.

**Figure 1:**
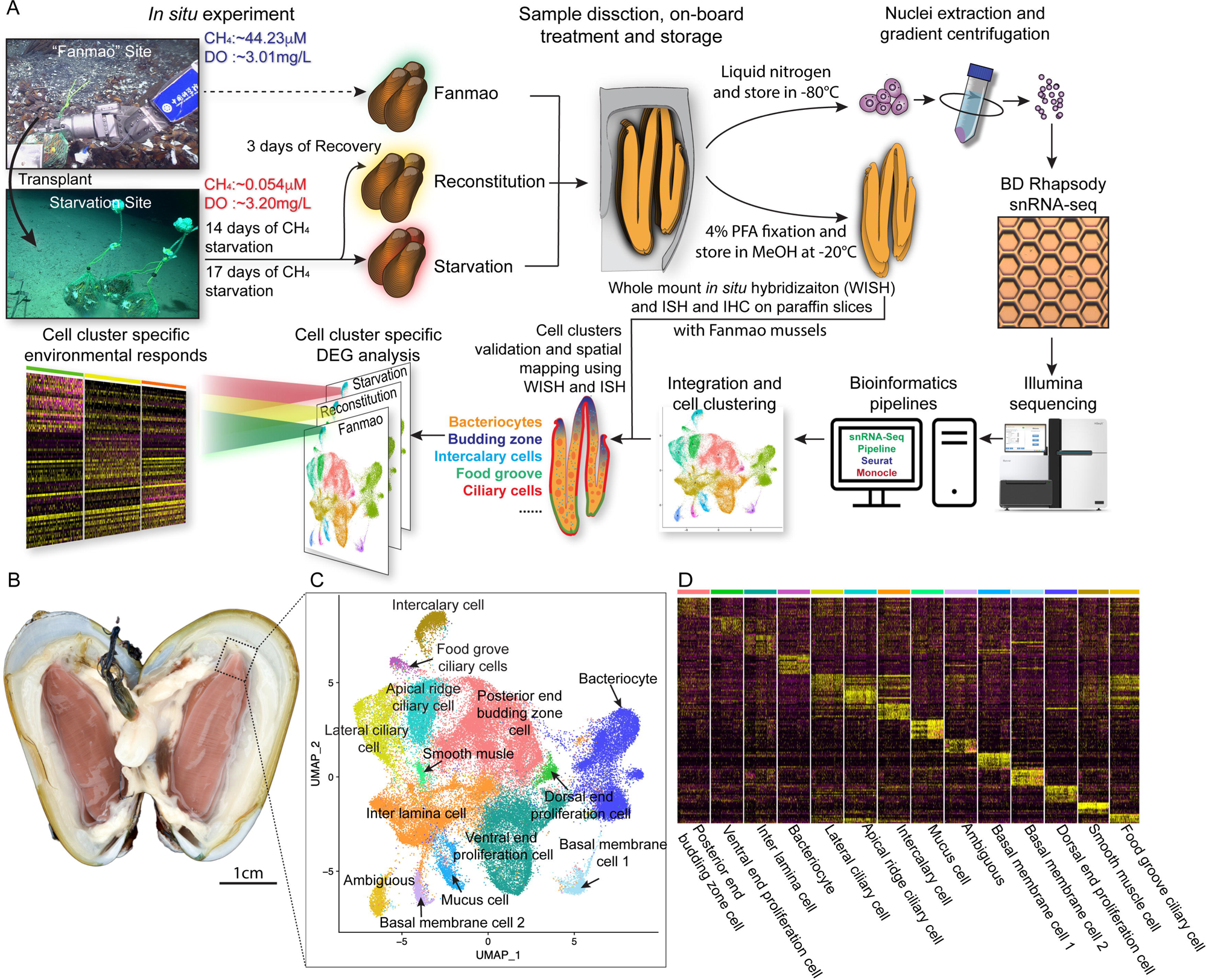
Identification of 14 cell types in the gill of deep-sea symbiotic mussel *Gigantidas platifrons*. A: Overall experimental scheme describing the deep-sea *in situ* transplant experiment, the sample preparation procedures and the single-cell analysis and validation pipeline. Three *G. platifrons* samples were included in the present study: ‘Fanmao,’ Starvation and Reconstitution. The cell nucleus was extracted from each sample, which included a pool of gill posterior tip of three mussels. The snRNA-seq libraries were constructed according to the BD Rhapsody single-nuclei 3′ protocol. Cell population-specific markers were validated by WISH and ISH. B: The image shows the posterior end of the gill of *G. platifrons*. C: UMAP representation of *G. platifrons* gill single cells. Cell clusters are coloured and distinctively labelled. D: Heat map profile of markers in each cluster. The colour gradient represents the expression level of every single cell.

We next performed single nucleus RNA-sequencing (snRNA-seq) of the gill posterior tip of the three groups of samples using the microwell-based BD Rhapsody platform^™^ Single-Cell Analysis System (Fig. 1A). After quality control, we obtained 9717 (Fanmao), 21614 (Starvation), and 28928 (Reconstitution) high-quality single nuclei transcriptomic data.

### 2. Cell atlas of *G. platifrons gill*

To unravel the intricate cellular composition, we utilised a reciprocal PCA (RPCA) strategy to project cells in the three samples onto the same space based on conserved expressed genes among them^25^. This strategy maximises the number of cells per cluster regardless of the organism’s state and therefore maximises genes per cluster^26^.

Through the implementation of Seurat, we revealed 14 cell clusters, each was associated with a set of marker genes (Figs. 1B-D) ^27,28^. Given the limited availability of canonical marker genes for *G. platifrons* and molluscs in general, we undertook a meticulous approach to characterize each cell cluster. This involved: 1) examining the cluster marker genes’ functions, 2) identifying the expression pattern of cluster marker gene using whole-mount *in situ* hybridisation (WISH) and/or double-fluorescent *in situ* hybridisation (FISH) analyses, and 3) conducting scanning electron microscopy (SEM) and transmission electron microscopy (TEM) analyses. We successfully identified and characterized 13 of the 14 cell clusters, which could be categorized into four major groups, including 1) the supportive cells, 2) the ciliary cells, 3) the proliferation cells, and 4) the bacteriocytes. One cell cluster remained ambiguous. To assess the robustness of each cell cluster, we employed a bootstrap sampling and clustering algorithm examining the similarity among clusters and obtained strong support for all clusters through a combined analysis of the three samples (Supplementary Fig. S2A). Furthermore, when examining each sample individually, we found that the majority of clusters demonstrated robust support, with the exception of the three ciliary cell clusters, which showed overlaps of assignment probabilities among them (Supplementary Fig. S2 B-D). These three cell types are derived from a same precursor, and exhibited relatively lower numbers of nuclei, resulting in a reduced availability of genes per cell type, which is particularly true for the food grove ciliary cell (Supplementary Table S1). By integrating the spatial and functional annotations of these cell clusters, we gained insight into their collective efforts in supporting the symbiotic relationship and maintaining the high-efficiency chemosynthetic system within the gill tissue.

#### Supportive cells

We have identified four supportive cell types, namely, inter lamina cells basal membrane cells (BMC)1, BMC2, and mucus cells (Figs 2A, B and C).

**Figure 2:**
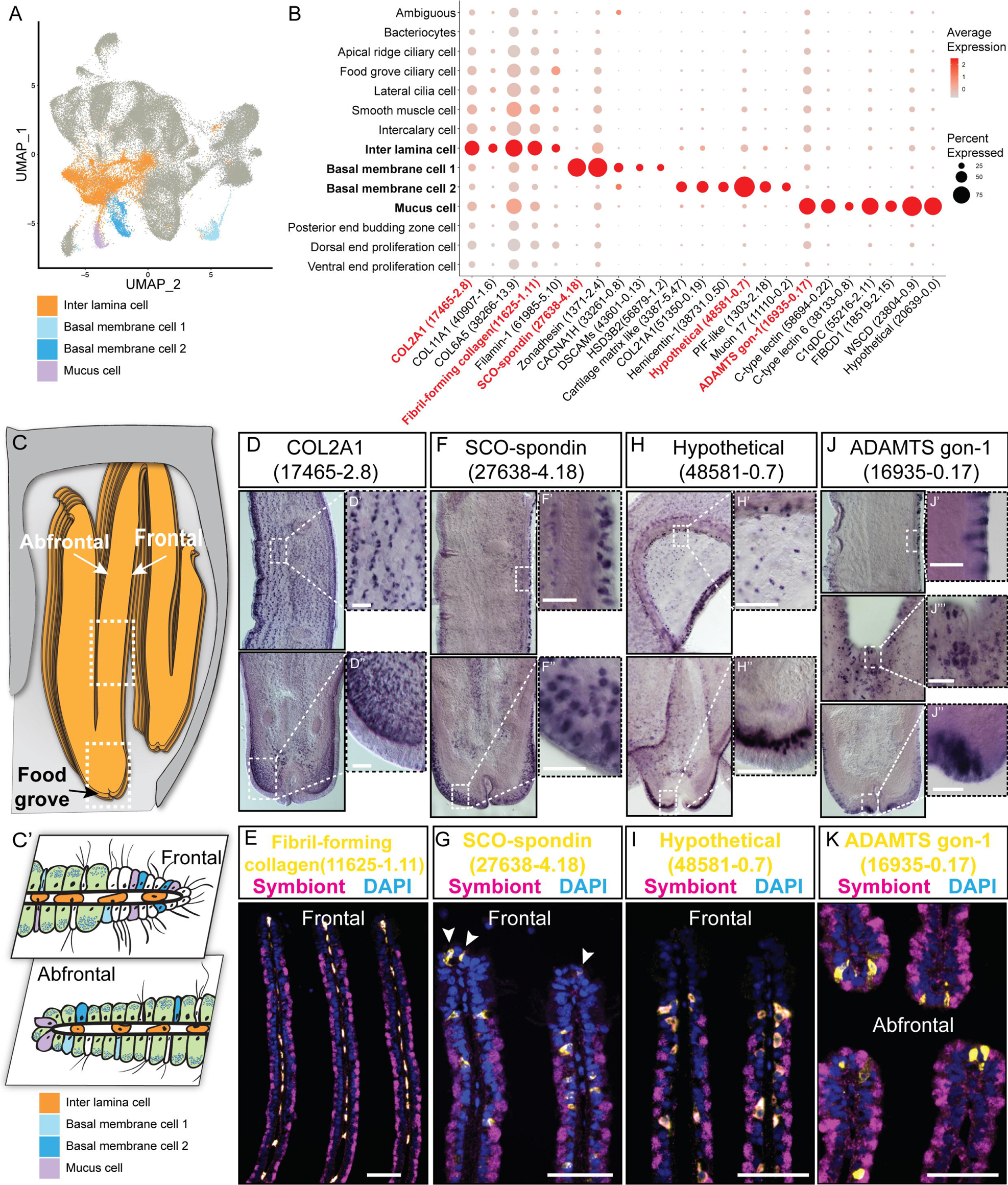
Supportive cell populations of *G. platifrons* gill. A: UMAP representation of the four supportive cell populations. B: Expression profiles of the cell markers that are specific or enriched in the supportive cell populations. The sizes of the circles represent the percentages of cells in those clusters that expressed a specific gene. Genes shown in red were validated by WISH or double FISH. C and C’: Schematics demonstrating the overall structural (panel C) and supportive cell distribution (C’). D, F, H, and J: WISH characterisation of the selected representative cell population markers. E, G, I, and K: Double FISH characterisation of the selected representative cell population markers. The white arrowheads in G indicate the BMC1 cells locates at the outer rim of gill slice. Scale bar: 50µm.

The inter lamina cells, marked by high expression of fibrillar-forming collagens, were the cells located between the two layers of basal membranes (Fig 2D and E). These cells were previously identified as amoeboid cells^13^, and their function has not been explored. The inter lamina cells were densely distributed around the food groove and at the rim of the gill filament. While on the middle part of the gill filament, the inter lamina cells distributed in parallel rows. It might help to connect the two sheets of basal membranes and maintain the spatial integrity of the gill filament. In addition, we observed the enrichment of ribosomal proteins involved in protein synthesis (Supplementary Table S2), indicating a high metabolic rate^29^ of inter lamina cells. This implies that the inter lamina cells may process the nutrients acquired from the symbiont.

We have also identified two types of BMC populations which have not been recognised before. The BMC1 (Fig. 2F and G) expressed genes encoding extracellular matrix and adhesive proteins (Fig. 2B and Supplementary Table S2), such as SCO-spondin (Bpl_scaf_27638-4.18), Zonadhesin (Bpl_scaf_1371-2.4), Cartilage matrix proteins (Bpl_scaf_16371-0.14 and Bpl_scaf_3387-5.47), and Collagen alpha-1(XXI) chain protein (Bpl_scaf_51350-0.19). BMC2 (Fig. 2H and I) was distinctive from BMC1 in the context of expression (Fig 2A) and marker genes (Fig 2B). Genes encoding hemicentins (Bpl_scaf_38731-0.50 and Bpl_scaf_7293-0.14), which are known to stabilise the cells’ contact with the basal membrane^30^, were highly expressed in BMC2. Thus, these two cell types are likely to help building the basal lamina, and stabilising the epithelial-derived cells, such as bacteriocytes and intercalary cells, on the surface of the basal membrane.

Mucus cells are specialised secretory cells with intracellular mucus vacuoles^13^. In shallow-water filter-feeding bivalves, mucus cells secret mucus, which cooperates with the ciliary ventilation system to capture, process, and transport food particles to their mouths, known as filter feeding mechanism^31–34^. Deep-sea mussels may not necessarily retain this function since they do not normally acquire food resources such as phytoplankton and planktonic bacteria but obtain most of nutrient through their chemosynthetic symbiont. Thus, it was hypothesised that mucus cells are involved in other biological functions such as immune responses to pathogens^18^. Herein, genes encoding proteins with microbe-binding functionalities were enriched in mucus cells (Fig. 2B and Supplementary Table S2), such as C-type lectins (Bpl_scaf_58694-0.22 and Bpl_scaf_38133-0.8), C1q domain-containing protein (Bpl_scaf_55216-2.11), and Fibrinogen C domain-containing protein (Bpl_scaf_18519-2.15)^35,36^. The expression of similar immune genes was upregulated in a shallow water mussel *Mytilus galloprovincialis* when challenged by a pathogenic bacteria^37^. Our WISH and double-FISH analyses showed that mucus cells were embedded within the outer rim cilia and scattered on the gill lamella alongside the bacteriocytes (Figs. 2J and 2K). These data collectively suggest that mucus cells may help mussel maintain the immune homeostasis of gill. Interestingly, WISH and double-FISH analyses showed that mucus cells were also distributed alongside the gill lamella’s food groove and the inner edge, where the density of bacteriocytes is low (Fig. 2J). Because the food groove is the main entry of food practical to the labial palps and mouth^38^, this distribution pattern implies that mucus cells may be also involved in capturing planktonic bacteria and sending them to the mouth.

#### Ciliary and smooth muscle cells

A remarkable feature of bathymodioline mussel’s gill is its ciliary ventilation system, which constantly agitates the water and provides the symbiont with the necessary gas^39^. We identified four types of ciliary cells (Figs. 3A and 3B): apical ridge ciliary cells (ARCCs), food groove ciliary cells (FGCCs), lateral ciliary cells (LCCs), and intercalary cells (ICs), as well as a newly identified type of smooth muscle cells. All ciliary cells were marked by canonical cilium genes, such as genes encoding flagellar proteins, ciliary motor kinesin proteins, ciliary dynein proteins, and ciliary microtubules and were clearly distinguishable by specifically expressed genes (Supplementary Table S3 and Supplementary Fig. S3). ARCCs were characterized by expression of Tubulin alpha-1A chain Bpl_3489-0.37 (Fig. 3C) and Bpl_scaf_20631-1.16, which encodes homeobox Dlx6a-like protein (Fig. 3D) which is a marker of the apical ectodermal ridge^40^. FGCCs, which could be ladled by expression of marker gene Bpl_scaf_5544-0.0 (Fig. 3E), expressed genes encoding primary cilia development regulator Tubby-related protein 3^41^ (Bpl_scaf_24834-2.3) and primary ciliary cell structural protein TOG array regulator of axonemal microtubules protein 1^42,43^ (Bpl_scaf_55620-1.3), suggesting the FGCCs are the sensory ciliary cells that gather information from the surrounding environment. Both ARCCs and FGCCs were located around the ventral tip of the gill filament (Fig. 3F). LCCs were distributed as two parallel rows along the gill lamella’s outer rim and ciliary disks’ outer rim (Figs. 3G and 3H) and had highly expressed genes involved in cilium structure, cilium movement and ATP synthases (Fig. 3B and Supplementary Table S3), indicating that LCCs may have a strong ability to beat their cilia. Interestingly, we identified a group of previously unreported smooth muscle cells (SMCs) co-localised with the LCCs as showed by WISH (Fig. 3I and Supplementary Fig. S4). SMCs strongly expressed several Low-density lipoprotein receptors (LDL receptor) and LDLR-associated proteins^44^ (Fig. 3B). In addition, these cells also expressed the angiotensin-converting enzyme-like protein^45^ (Bpl_scaf-29477-5.10) and the ‘molecular spring’ titin-like protein^46^ (Bpl_scaf_56354-7.8) (Supplementary Table S3). The expression of these genes could be commonly found in human vascular smooth muscle cells^47–49^. Collectively, we suspect that SMCs are involved in lateral cilium movement and the gill slice contraction.

**Figure 3:**
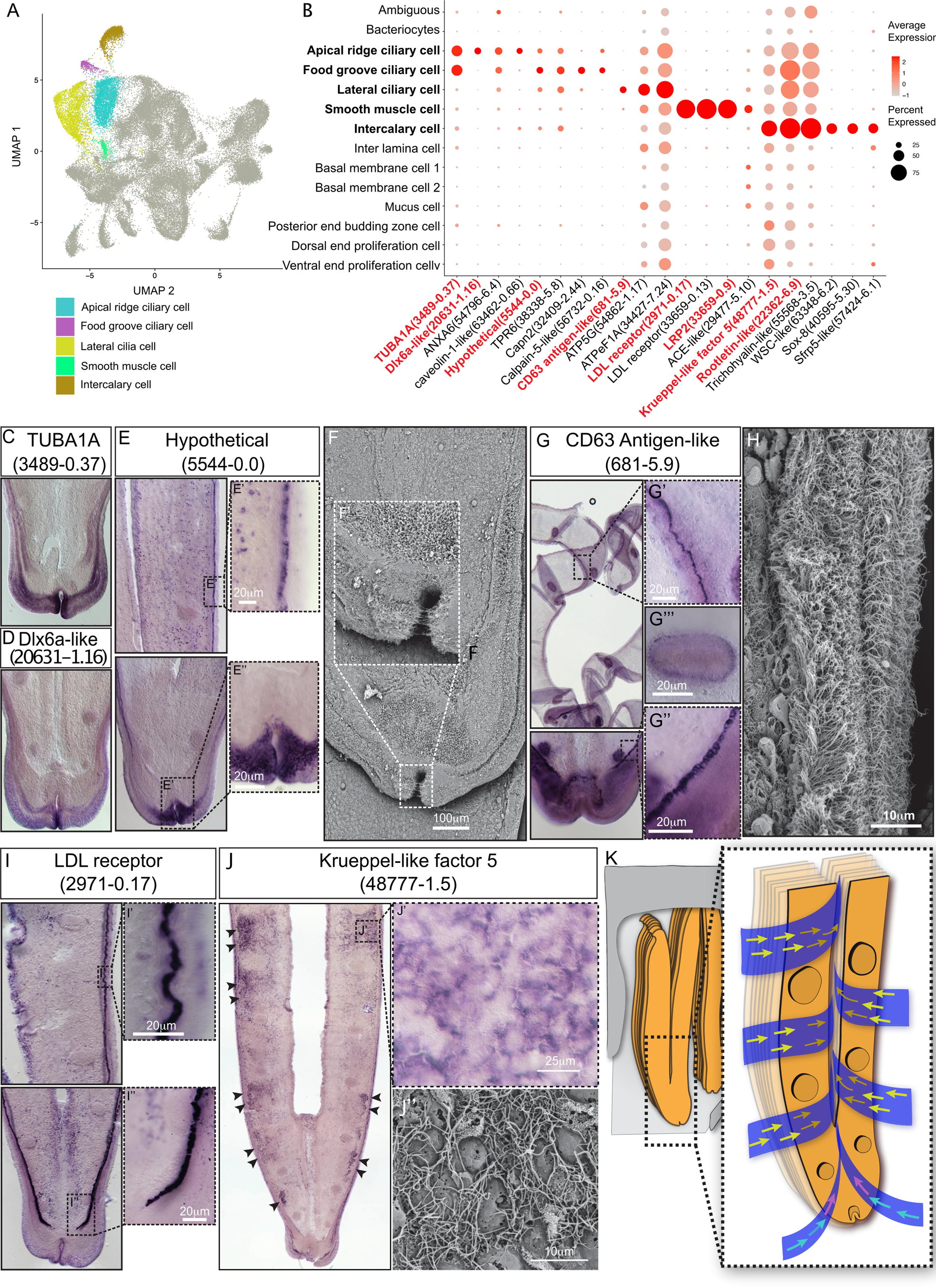
Ciliary cell populations of *G. platifrons* gill. A: UMAP representation of the four ciliary cell populations and potential smooth muscle cell population. B: Expression profiles of the cell markers that are specific or enriched in the ciliary cell populations. The sizes of the circles represent the percentages of cells in those clusters that expressed a specific gene. The genes shown in red were validated by WISH or double FISH. C-E, G and I-J: WISH characterisation of the selected representative cell population markers. F and H: SEM analysis of the ciliary cells of *G. platifrons* gill. K: Schematic of the water flow agitated by different ciliary cell types. The coluor of arrowheads corresponds to water flow potentially influenced by specific types of cilia, as indicated by their colour code in Figure 3A.

ICs are the specialised ciliary cells surrounding the bacteriocytes (Figs. 3B and 3J). WISH analysis showed that the expressions of IC markers had apparent spatial variations (Fig. 3J). The expression of ICs marker rootletin^50–52^ (Bpl_scarf_22362-6.9) was considerably higher in the ICs close to the frontal edge of the gill filament, and the expression gradually decreased along with the direction of inter lamina water flow (Fig. 3K and Supplementary Fig. S5), implying that ICs ventilate the water flow and the mucus through the gill filaments. Furthermore, compared with the other three types of ciliary cells, the ICs expressed several genes encoding transcription factors involved in determining cell fate (Fig. 3B and Supplementary Table S3), such as transcription factors Sox 8 (Bpl_scarf_40595-5.30) and Wnt pathway cell polarity regulator secreted frizzled-related protein 5 (Bpl_scaf_57424-6.1)^53^, suggesting that the ICs might also play regulatory roles^54–56^.

#### Proliferation cells

It has long been known that the bathymodioline mussel gill has three types of proliferation cells that are conserved throughout all filibranchia bivalves: the budding zone at the posterior end of the gill where new gill filaments are continuously formed, and the dorsal and ventral ends of each gill filaments^57,58^. These “cambial-zone”-like cell populations could continuously proliferate throughout the whole life span of the mussel^58^. Our snRNA-seq data recognised three types of proliferation cells, which is consistent with previous findings (Fig. 4A, Fig. 4B and Supplementary Table S4). The gill posterior end budding zone cells (PEBZCs) are located on the first few freshly proliferated filaments of the posterior tip of gill (Figs. 4C and 4D). The PBEZCs marker (Bpl_scaf_61993-0.4) gradually disappeared at around the 11th-12th row of the gill filament (Fig. 4D), suggesting the maturation of the gill filaments, which is similar to the developmental pattern reported in another deep-sea mussel *Bathymodiolus azoricus* ^59^. The dorsal end proliferation cells (DEPCs), which expressed the hallmarks genes of muscular tissue (Fig. 4B), as well as cell proliferation and differentiation regulators, were the proliferation cells in connective tissue at the dorsal end of the gill slice (Fig. 4A, 4E). The ventral end proliferation cells (VEPCs) were two symmetrical triangle-shaped cell clusters of small symbiont-free cells (Fig. 4F). In VEPCs, genes encoding ribosomal proteins, chromatin proteins, RNA and DNA binding proteins, and cell proliferation markers were all up-regulated (Fig. 4B and Supplementary Table S4), indicating that VEPCs are meristem-like cells^59,60^.

**Figure 4:**
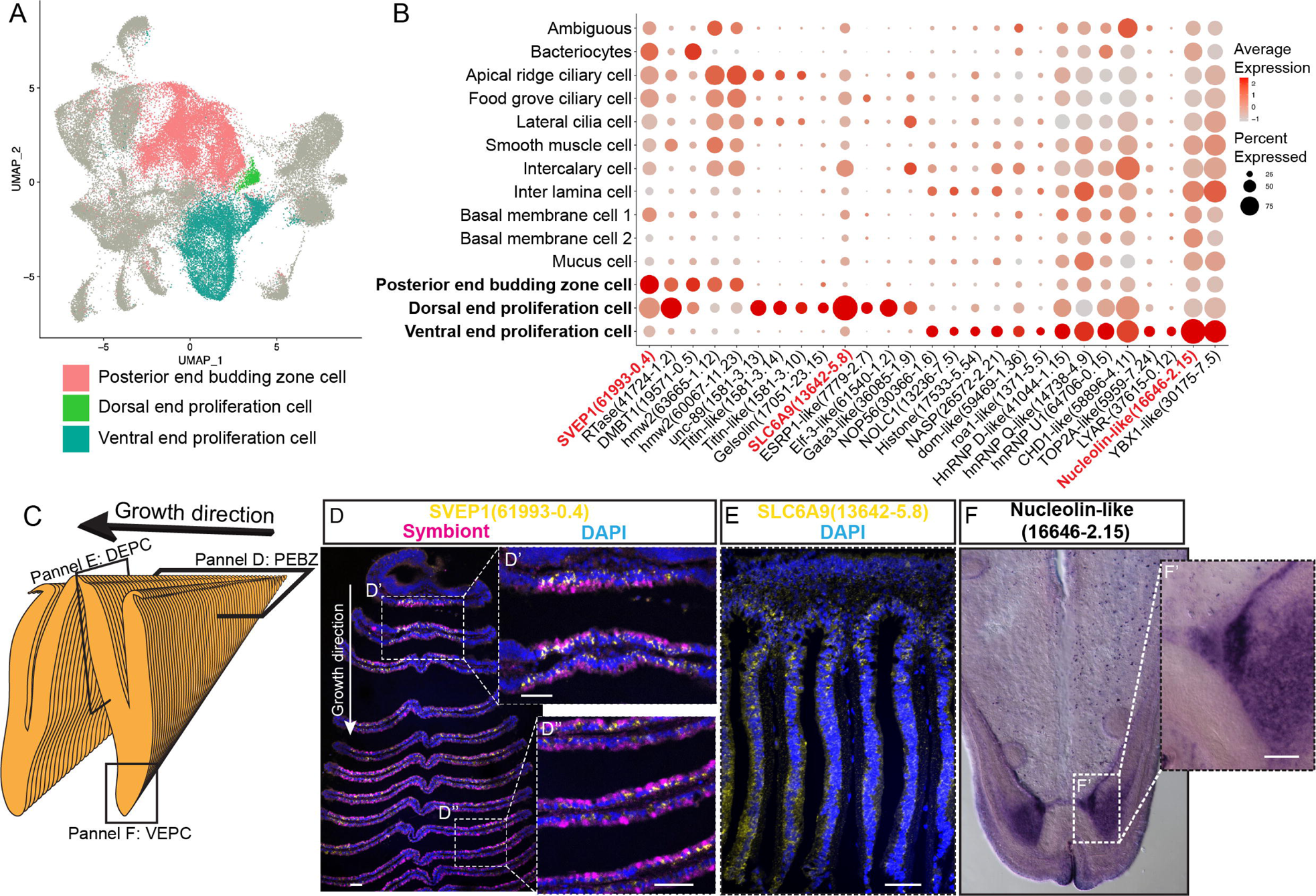
Proliferation cell populations of *G. platifrons* gill. A: UMAP representation of the three proliferation cell populations. B: Expression profiles of the cell markers that are specific or enriched in the supportive cell populations. The sizes of the circles represent the percentages of cells in those clusters that expressed a specific gene. Genes shown in red were validated by WISH or double FISH. C and D: Photographic and schematic analyses of the spatial position of the three Proliferation cell populations. E, F and J: FISH and WISH characterisation of the selected population markers. The marker genes confirmed by ISH or WISH in the current study are indicated in red. Scale bar: 50 µm.

It has been hypothesised that new proliferation cells will be colonised by symbionts, serving as the vital mechanism of bacteriocyte recruitment^59^. We then determined which proliferation cell type gives rise to the mature cells, especially the bacteriocytes, which has little available information regarding their precursor and transmission mode in bathymodioline mussels ^59,61^. We performed Slingshot analysis, which uses a cluster-based minimum spanning tree (MST) and a smoothed principal curve to determine the developmental path of cell clusters. The result shows that the PEBZCs might be the origin of all gill epithelial cells, including the other two proliferation cells (VEPC and DEPC) and bacteriocytes (Supplementary Fig. S6). The sole exception was BMC2, which may be derived from VEPC rather than PEBZC. This result is consistent with previous studies which suggested new gill filaments of the filibranch mussels are formed in the gill’s posterior budding zones^59^. The colonisation by the symbiont might play a crucial role in determining the fate of the bacteriocytes. Noticeably, in *G. platifrons*, only the pillar-shaped first row of gill filament, comprised of small meristem-like cells, was symbiont-free (Figs. 4D and D’), whereas all the other gill filaments were colonised by symbionts. This pattern of symbiosis establishment is different from that of *B. azoricus*, in which PEBZCs are symbiont free and are gradually colonised by symbionts released from the bacteriocytes on the adjacent mature gill filaments after maturation^59^. On mature gill filaments, the DEPCs and VEPCs are seemingly the sources of new cells that sustain the growth of the gill filament from both dorsal and ventral directions, respectively^57^. Interestingly, comparable active ventral and dorsal end proliferation zones have also been identified in the symbiotic mussel *B. azoricus*, whereas they are absent in the shallow-water mussel *Mytilus edulis*^62^. This contrast further suggests the potential involvement of DEPCs and VEPCs in the establishment of symbiosis.

### 3. Bacteriocytes and host-symbiont interaction

We conducted the whole-mount FISH using a bacterial 16S rRNA probe for symbiont to determine the spatial distribution of bacteriocytes, and their positions relative to the other cell types on the gill filament (Figs. 5A, B). Bacteriocytes covered the majority of the surface of the gill filament, except the ventral tip, the ciliary disk, and the frontal edge (lateral ciliary). The bacteriocytes were surrounded by intercalary cells with microvilli and cilia on the surface (Fig. 5C).

**Figure 5:**
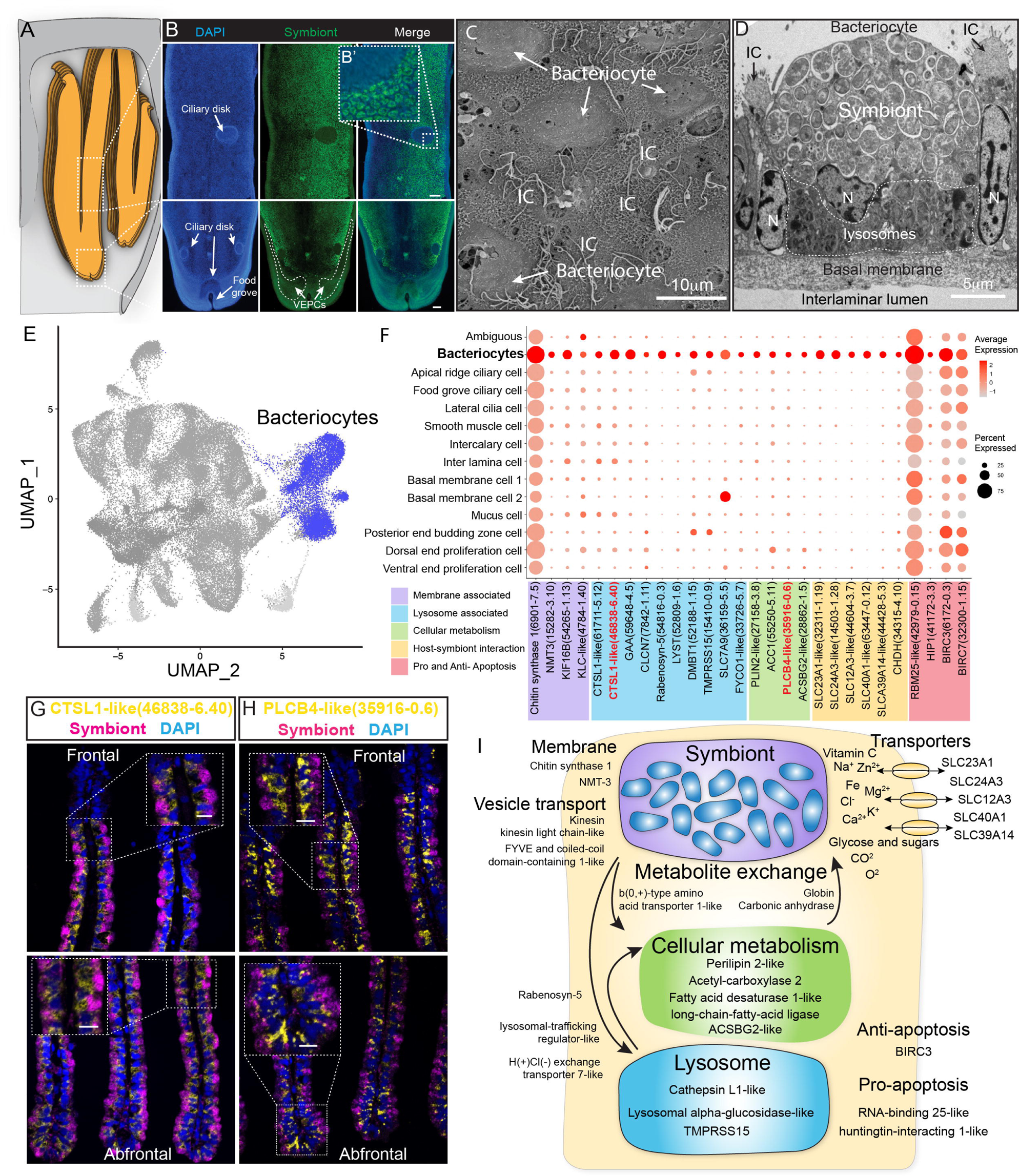
Characterisation of the bacteriocytes of *G. platifrons*. A: Schematic of the overall structure of *G. platifrons* gill filaments. B: Whole-mount FISH analyses of the overall distribution of bacteriocytes on the *G. platifrons* gill filament. C: SEM analysis of the bacteriocytes. D: TEM analysis of a bacteriocyte. E: UMAP representation of *G. platifrons* bacteriocytes. F: Expression profiles of the cell markers that are specific or enriched in the bacteriocytes. The sizes of the circles represent the percentages of cells in those clusters that expressed a specific gene. Double FISH validated the genes shown in red. I: Schematic of the host-symbiont interaction based on the single-cell transcriptome of *G. platifrons* bacteriocytes. The marker genes confirmed by ISH in the current study are indicated in red. Scale bar in panel G and H: 25 µm.

The gene expression profile of bacteriocytes aligned well with the ultrastructural analysis, which suggested that the bacteriocytes have structural, metabolic, and regulatory adaptions to cope with the symbiont (Figs. 5D-5I). It has been hypothesised that the bacteriocytes extract nutrients from the symbiont through two pathways: lysing the symbiont through the lysosome (the ‘farming’ pathway) or directly utilising the nutrient produced by the symbiont (the ‘milking’ pathway)^63,64^. Our TEM observations clearly detected intracellular vacuole and lysosome system of the bacteriocyte that harbour, transport, and digest the symbionts (Fig. 5D), which is consistent with previous studies^18,65^. The snRNA data showed bacteriocytes expressed cellular membrane synthesis enzyme (phosphoethanolamine N-methyltransferase, Bpl_scaf_15282-3.10) and a series of lysosomal proteins such as lysosomal proteases (cathepsins; Bpl_scaf_61711-5.12, Bpl_scaf_46838-6.40 and Bpl_scaf_59648-4.5; lysosomal alpha-glucosidase-like), protease regulators 56 (TMPRSS15 proteins; Bpl_scaf_52188-1.15 and Bpl_scaf_15410-0.9), and lysosomal traffic regulator proteins (rabenosyn-5, Bpl_scaf_54816-0.3 and Bpl_scaf_52809-1.6, lysosomal-trafficking regulator-like isoform X1). Among the proteases, cathepsins were thought to be evolutionary conserved molecular tools that host utilized to control the residence of their symbiont microbes^66^. They were also highly expressed in symbiotic tissue of other deep-sea chemosynthetic animals, such as vesicomyid clams and vestimentiferan tubeworms^67–69^. Bacteriocytes also expressed genes encoding cellular vesicle transports (kinesins, Bpl_scaf_14819-0.11, Bpl_scaf_54265-1.13, and Bpl_scaf_4784-1.40)^53^, potential amino acid transporter^57^ (Bpl_scaf_36159-5.5), and genes involved in intracellular vesicle transport, such as the FYVE and coiled-coil domain-containing protein 1^58^ (Bpl_scaf_33726-5.7). In addition to form and mobilise early endosomes, the protein products of these genes could transport symbiont-secreted nutrient vesicles to the host (Fig. 5F and Supplementary Table S5), supporting the “milking” pathway.

The symbiont of *G. platifrons* belongs to type I methanotrophy, of which the core metabolic function is linked with the development of intracytoplasmic membranes leading to a high lipid/biomass content^70,71^. Recent lipid biomarker analyses showed that the gill of *G. platifrons* contains a high amount of bacterial lipids, which are directly utilised by the host to synthesise most of its lipid contents^67^. Downstream of the bacteriocyte’s metabolic cascade, genes encoding proteins that may be involved in fatty acid/lipid metabolism, such as perilipin2 (Bpl_scaf_27158-3.8), which is the critical protein to form intracellular lipid droplets^72^, and a variety of fatty acid metabolism enzymes (acetyl carboxylase 2, Bpl_scaf_55250-5.11^73^; fatty acid desaturase 1-like isoform X1, Bpl_scaf_35916-0.6^74,75^; long-chain fatty acid-ligase ACSBG2-like isoform X1, Bpl_scaf_28862-1.5 ^76,77^), were up-regulated, suggesting that the fatty acid could be a major form of nutrients passing from the symbiont to the host mussel.

Additionally, bacteriocytes expressed several solute carriers, including sodium/ascorbate co-transporter (Fig. 5I, solute carrier family 23 members 1-like, Bpl_scaf_32311-1.19), sodium/potassium/calcium exchanger (sodium potassium calcium exchanger 3-like, Bpl_scaf_14503-1.28), sodium/chloride ion cotransporter (solute carrier family 12 member 3-like isoform X1, Bpl_scaf_44604-3.7), ferrous iron transporter (solute carrier family 40 member 1-like, Bpl_scaf_63447-0.12) and zinc transporter (zinc transporter ZIP14-like, Bpl_scaf_44428-5.3). The solute carriers are a large family of ATP-dependent transporters that shuttle a variety of small molecules across the cellular membrane^78^. In several model symbiotic systems, solute carriers play a vital role in host-symbiont interaction by either providing the symbiont with substrates^79,80^ or transporting symbiont-produced nutrients to the host ^81–83^. Previous studies demonstrated that solute carrier genes were expanded in deep-sea chemosynthetic animals’ genome^84^ and highly expressed in symbiotic organs^85^ including bathymodioline mussel’s gill^12^. In bathymodioline mussels, previous bulk RNA-seq studies detected up-regulated expression of a large variety of solute carriers in the gill, which is consistent with the present study, suggesting that solute carriers may play crucial roles in shuttling nutrients in and out of bacteriocytes and in maintaining the suitable intracellular micro-environment (such as the SLC23A1 and SLC39A14) for the symbiont^86,87^.

### 4. Cell-type-specific response to environmental stresses

To examine cell-type-specific acclimatisation to environmental changes the expression profiles of differentially expressed genes (DEGs) were compared between the three states (Fanmao, starved and reconstituted animals) of samples collected in our *in situ* transplantation experiment (full lists of DEGs are shown in Supplementary Table S6-S18). Notably, for each state three mussels were processed but pooled for nuclei extraction before snRNA sequencing (see Method for detail). We tested our hypothesis here that at the starvation site, a relatively low concentration of methane shall upset symbiont metabolism and thus substantially affect symbiont-hosting bacteriocytes by assessing the transcriptional changes per cell type. We calculated the centroid coordinates for each cell type in each state on the 2-dimensional UMAP plot (Fig. 6A). Then, for each cell type, we determined the Euclidean distance between the centroid coordinates of each pair of states (Supplementary Table S19). The impact of starvation was variable across cell types, as reflected by the cross-state distances (Fig. 6B, green bar). Starvation resulted in the most significant transcriptional changes in bacteriocytes reflected by a large Fanmao-vs-starvation distance (2.3), followed by VEPC and inter lamina cells (2.1 and 1.4, respectively). On the other hand, starvation had little impact on the transcriptions of ciliary cells and most supportive cells such as the food grove ciliary cell, BMC2, and mucus cells (distances <0.5). Fig 6B also shows that after reconstitution (moving the mussels back to the methane-rich site Fanmao for 3 days), the expressional profile of bacteriocytes rapidly changed back, reflected by a large starvation-vs-reconstitution distance and much smaller Fanmao-vs- reconstitution distance (2.3 vs. 0.8; Suppl. Tab. S19). This result coincided with and was supported by our pseudo-time analysis for bacteriocytes (Fig. 6C), showing that bacteriocytes in the reconstitution are in intermediate and transitional states between Fanmao and starvation.

**Figure 6:**
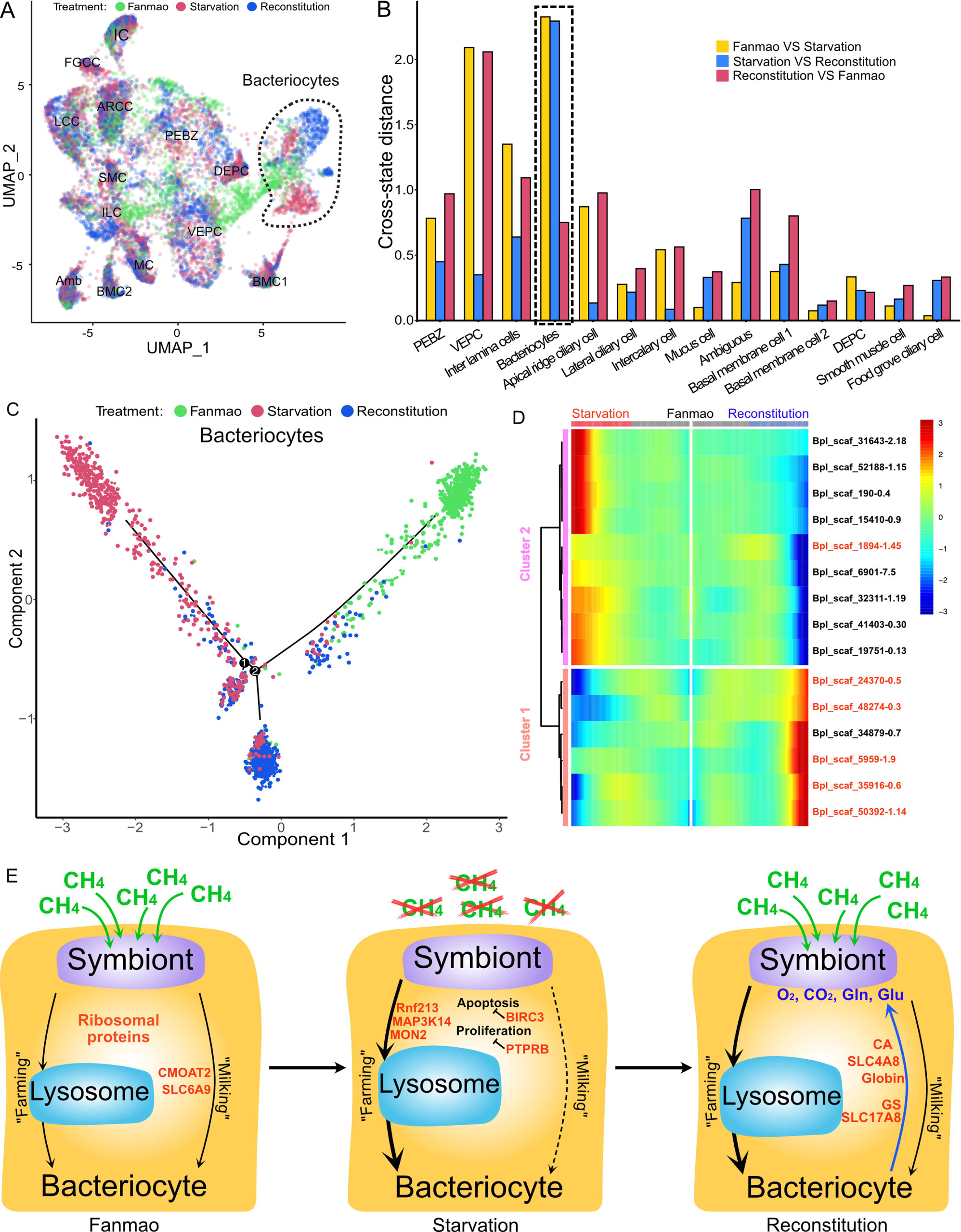
Analysis of cell population-specific DEGs. A: UMAP representation of the impact of deep-sea *in situ* transplant treatments on the gene expression pattern of each cell population. The cells from different treatments were labelled with different colours. The dashed line encircled bacteriocyte populations have a considerably altered expression profile. B: Histogram of cross-state distances between the centroids of the Fanmao, Starvation and Reconstitution groups per cell type on UMAP. The black dashed lines indicate the bacteriocyte populations whose expression profile was remarkably altered. C: Visualization of bacteriocytes onto the pseudo time map using monocle. The black lines indicate the main path of the pseudotime ordering of the cells. D: Bifurcation of selected gene expression along two branches in response to environmental perturbation. Genes are clustered hierarchically into two groups, illustrating up- (cluster 1) and down- (cluster2) regulated genes in the starvation state compared with Fanmao. Genes in red colour were discussed in Section 4. The heat map showing the gene expression profiles of all bacteriocytes’ DEGs is shown in supplementary Fig. S7; E: Proposed model for the molecular mechanisms of host-symbiont interactions in response to environmental changes.

For the bacteriocytes population, we conducted cell trajectory pseudo-timing and detailed DEG analysis to interpret the mechanism of host-symbiont interactions. It is important to acknowledge that our sampling strategy might have limitations, as more closely spaced time points could enhance the confidence of trajectory reconstruction. The branched heatmap showed both up- and down-regulated genes during starvation and reconstitution compared with the Fanmao state (Fig. 6D and supplementary Fig. S7). The Kyoto Encyclopedia of Genes and Genomes (KEGG) pathway enrichment analysis of the bacteriocytes’ DEGs provides an overall view of the pathways enriched in each environmental state (Supplementary Fig. S8 A-C). The genes encoding ribosomal proteins are highly expressed in the bacteriocytes under the methane-rich ‘Fanmao’ state (Supplementary Fig. S8A), suggesting an active protein synthesis and cellular metabolism ^88^. Moreover, organic ion transporters (A BCB P-glycol, Bpl_scaf_18613-16.10; canalicular multispecific organic anion transporter 2, Bpl_scaf_52110-4.43; sodium- and chloride-dependent glycine transporter 1-like, Bpl_scaf_13642-5.8), which may be involved in transporting symbiont-produced nutrients, are also highly expressed (Supplementary Fig. S9).

In starved *G. platifrons*, negative regulators of cell proliferation, such as autocrine proliferation repressors and receptor-type tyrosine phosphatase beta, were up-regulated in bacteriocytes, suggesting repression of cell growth. We also observed the enrichment of genes in the apoptosis pathway (Supplementary Fig. S8B), which is attributed to the upregulation of caspases (Bpl_scaf_25165-3.8 and Bpl_scaf_24225-0.12) and cathepsins (Bpl_scaf_64706-0.16, Bpl_scaf_61711-5.12) which can trigger caspases-dependent cell death, suggesting cellular stress condition. Correspondingly, a gene encoding baculoviral IAP repeat-containing protein (Bpl_scaf_6172-0.3), which can bind caspase and inhibit apoptosis, was up-regulated in bacteriocytes (Fig. 6E). In *Drosophila*, the baculoviral IAP proteins are important in animal’s response to cellular stress and promote cell survival^89,90^. Similarly, the gene encoding MAP3K14 (Bpl_scaf_60908-4.14) and potential E3 ubiquitin ligase RNF213 (Bpl_scaf_25983-1.26, Bpl_scaf_1894-1.45, Supplementary Fig. S10), which were apoptosis suppressor or regulator, were up-regulated^91^. These results suggest that starved bacteriocytes were carrying out cell-type-specific adjustments to cope with stresses. The E3 ubiquitin ligases could also work as intracellular immune sensors of bacterial lipopolysaccharides^92^. Thus, the encoded protein may be able to active downstream immunological toolkit to digest the symbiont population for nutrients.

As mentioned above, bacteriocytes obtain nutrients from endosymbionts. KEGG pathway analyses suggested that phagosome, lysosome-related pathways were upregulated in the starvation state, indicating that bacteriocytes were actively digesting the endosymbionts in the starved *G. platifrons*. Interestingly, protein synthesis activity was more activated in inter lamina cells and VEPCs as supported by the up-regulation of genes encoding ribosomal proteins in both cell populations (Supplementary Tables S6 and S17). This was contrary to the situation in bacteriocytes and may be a consequence of the high activity of “farming” in bacteriocytes.

We anticipated that after moving the starved *G. platifrons* back to Fanmao site, bacteriocytes and their endosymbionts would “reconstitute” leading to the partly restored “farming” and “milking” pathways. This was confirmed by higher expression of fatty acid metabolic genes in the mussels in the reconstitution state than those in the starvation state, such as long-chain-fatty-acid-ligase ACSBG2-like (Bpl_scaf_28862-1.5), elongation of very long-chain fatty acids 7-like (Bpl_scaf_5959-1.9), and fatty acid desaturase 1-like (Bpl_scaf_35916-0.6). These findings were also consistent with the result of KEGG analyses. The mitochondrial trifunctional enzyme (Bpl_scaf_42376-0.5) was also highly expressed, suggesting a high level of energy-producing activity. The KEGG analyses showed that glutamatergic synthase was enriched in the reconstitution state in comparison with those in both the Fanmao and starvation states. In the insect aphid–Buchnera endosymbiosis model, the host-produced glutamate could be transported and directly utilised by the symbiont^93^. Similar glutamate-based host–symbiont metabolic interaction mechanisms were also proposed in the deep-sea mussel *Bathymodiolus thermophilus* and *B. azoricus*^64,94^. As the pseudotime analyses showed above (Fig. 6C), reconstitution was the intermediate state between Fanmao and starvation. Metabolic and gene regulatory functions of bacteriocytes were still different from that in Fanmao. The KEGG analyses suggested regulatory (Rap1 signalling, retrograde endocannabinoid signalling, chemokine signalling, glucagon signalling, thyroid hormone synthesis, mRNA surveillance pathway, etc.) and metabolic pathways were enriched in the reconstitution state (fatty acid metabolism, salivary secretion, aldosterone synthesis and section, insulin secretion and pancreatic section) (Supplementary Fig. S8C). For example, we detected up-regulation of the genes encoding carbonic anhydrases (Bpl_scaf_33596-7.3 and Bpl_scaf_48274-0.3), electroneutral sodium bicarbonate exchanger 1 (Bpl_scaf_61230-5.11), and globin-like proteins (Bpl_scaf_50392-1.14 and Bpl_scaf_24370-0.5), which could provide the symbiont with carbon dioxide and oxygen necessary for the symbiont’s chemosynthetic metabolism^84,95^.

### 5. Summary and outlook

Using the deep-sea mussel *G. platifrons* as a model organism, we demonstrated the power of integrating snRNA-seq and WISH data in unravelling the mechanism behind animal-microbe symbiosis. The robustness of this strategy showed stable and highly distinguishable expression patterns of each cell type regardless of the different environmental states. We successfully profiled the specific roles of different types of cells, including the previously unknown cell types, in maintaining the structure and function of the gill. The supportive cells that located in between (inter lamina cells) and on the basal membrane’s surface (BMC1 and BMC2) helped maintain the anatomical structure of the basal membrane. Ciliary and smooth muscle cells involved in gill slice contraction and cilium beat allowing the gathering of material and information from the surrounding environment. Proliferation cells (PBEZCs, DEPCs and VEPCs) gave rise to new cells, including bacteriocytes which obtained nutrients from endosymbionts using intracellular vacuoles and lysosomes.

The snRNA analyses also revealed that different cell types collaborated to support bacteriocytes’ functionality. The PBEZCs gave rise to new bacteriocytes that allowed symbiont colonisation. Bacteriocytes attached to the basal membrane and were stabilised by supportive cells. Mucus cells co-localise with bacteriocytes and help to maintain immune homeostasis. The beating ciliary cells controlled the water flow, providing bacteriocytes with necessary inorganic substances from the environment. This new information on cell-cell interaction certainly advanced our overall understanding of how endosymbiotic microbes and host cells communicate and collaborate, which cannot be easily achieved through other methods.

Moreover, the analysis of snRNA data from bacteriocytes has provided insight into the molecular mechanisms employed by the host to maintain and regulate the symbiosis. Notably, the bacteriocytes-enriched transcripts involved in harbouring, digesting symbionts, and transporting nutrients produced by the symbionts were identified and characterised. Our *in situ* transplant experiments by moving mussels between methane-rich and methane-limited sites also provided clues of cell-type specific responses to environmental change. Under a methane-limited environment, the staved mussels more actively consumed endosymbionts through the “farming” pathway. After being moved back to the methane-rich environment, mussels produced more glutamates which sustained the regrowth of symbionts. These preliminary findings showed that the deep-sea mussels were able to control their endosymbionts using a set of genes in response to environmental change. Due to the limitation of remotely operated vehicle (ROV) dives and sampling capacity, we had to have pooled samples of each state for nuclei extraction and sequencing. Thus, the cells per cluster could be considered as technical than biological replicates. Although this sampling process strategy has been broadly used in snRNA/scRNA sequencing^96^, we recognise the possible violation of assumptions in *p*-value calculation for DEGs between the three states.

Overall, the single-cell spatial and functional atlas developed in the present work will decipher some common principles of symbiosis and environmental adaption mechanisms of animals. The workflow developed in the present study could provide insightful references for researchers focusing on the mechanistic study of the biological adaptation of biologically and ecologically important non-model animals.

## Acknowledgements

This study was supported by the Science and Technology Innovation Project of Laoshan Laboratory (Project Number No. LSKJ202203104), the National Natural Science Foundation of China (Grant No. 42030407), Southern Marine Science and Engineering Guangdong Laboratory (Guangzhou)(HJ202101, SMSEGL20SC01), Major Project of Basic and Applied Basic Research of Guangdong Province (2019B030302004), the Research Grants Council of Hong Kong (C6026-19G-A and 16101822) and the Guangdong Natural Science Funds for Distinguished Young Scholar (2022B1515020033). We appreciate all the assistance provided by the crew of *R/V* ‘Kexue’ and the operation team of ROV ‘Faxian’.

## Author contributions

H.W., K.C., P.Y.Q. and C.L.L designed the study. H.Z., L. Cao, Z.S.Z., L. Chao and M.X.W. conducted the *in situ* transplant experiments and collected the samples; Q.Y.Z. and H.W. constructed the snRNA-seq libraries. K.H. performed the bioinformatics analysis. H.W., H.Z., L. Cao, J.L., H.C. and L.Z. performed the experiments. This paper was written by H.W., K.H., K.C., P.Y.Q. and C.L.L All authors revised the manuscript. All authors read, approved, and contributed to the final manuscript.

## Materials and methods

### Deep-sea *in situ* transplant experiment and sample collection

*In situ* transplant experiment was conducted at the ‘F-site’ cold seep during the R/V ‘Kexue’ 2020 South China Sea cold seep cruise. The overall design of the *in situ* transplant experiment and environmental states are shown in Figure 1. First, mussels in the methane-rich ‘Fanmao’ site (meaning ‘prosperous’ site, 22°06′55.425″N,119°17′08.287″E, depth 1117m) were scoped into three nylon bags with approximately 10 mussels in each bag. Then, two bags of mussels were transplanted to the low methane Starvation site (Figure 1, 22°07′00″N, 119°17′07.02″E, depth 1147.42m). After 11 days of transplantation, one bag of mussels in the Starvation site was moved back to the ‘Fanmao’ site. On the 14th day of the transplant, three bags of mussels: one bag from the Fanmao site (designated as the ‘Fanmao’ sample), one bag from the starvation site (designated as the ‘Starvation’ sample) and one bag of mussels which were first transplanted to the Starvation site for 11 days and moved back to the Fanmao site for 3 days (designated as the ‘Reconstitution’ sample) were all retrieved by ROV Fanxian. The mussels were kept in a hydraulic pressure-sealed biobox during the ascending of the ROV. The biobox is made of heat-insulated material, which will prohibit heat exchange with warm surface water. The samples were immediately processed once onboard R/V Kexue. For snRNA-seq, the posterior end tip of the mussel’s gill was dissected (Figure 1C), snap-frozen with liquid nitrogen and then stored at −80 °C until use. For WISH, ISH and FISH, the gills of the mussels were first fixed with 4% paraformaldehyde (PFA, prepared with autoclaved 0.22 μM membrane filtered *in situ* seawater) at 4 °C overnight. Then, the gill was washed with ice-cold 1× PBS three times, dehydrated and stored in 100% methanol at −20 °C.

### Single-nucleus RNA-sequencing of *G. platifrons*

The gill nucleus was extracted using the Nuclei PURE prep nuclei isolation kit (Sigma-Aldrich). For each sample, posterior end tips from 3 individual mussels were randomly selected and pooled together. The posterior tips were homogenised in 10 mL of ice-cold lysis solution (Nuclei PURE lysis buffer with 0.1% Triton X100 and 1 mM DTT) on ice. The cell nuclei were then separated by sucrose gradient centrifugation according to the manufacturer’s protocol. The cell nuclei pellets were washed by re-suspending in ice-cold DPBS–BSA solution (1× DPBS, 0.04% nuclease-free BSA, 0.01% RNase inhibitor; Takara). The nucleus was spun down by centrifugation at 500×g for 5 min at 4 °C. This step was repeated to remove the contaminants from the cell plasma. Finally, the nucleus was re-suspended in the DPBS–BSA solution. The concentration of the cell nucleus was counted by Cell Countess II. Single-nucleus RNA-seq libraries were then constructed with the BD Rhapsody™ single-cell analysis system using the BD Rhapsody WTA Amplification Kit according to the manufacturer’s protocol. The library was subjected to 150bp paired-end sequencing using the Illumina HiSeq 4000 platform. The clean reads of three datasets were submitted to the National Centre for Biotechnology Information Sequence Read Archive database (Bioproject: PRJNA779258).

### Bioinformatics

The three raw datasets (Fanmao, Starvation, and Reconstitution) were processed individually following the BD single-cell genomics analysis setup user guide (Doc ID: 47383 Rev. 8.0) and BD single-cell genomics bioinformatics handbook (Doc ID: 54169 Rev. 7.0). This process involved the preparation of gene names, alignment of data to the genome, and generation of an expression matrix for each dataset. A *G. platifrons* genome reference v1.0 (available at the Dryad Digital Repository; http://dx.doi.org/10.5061/dryad.h9942) was utilized. The genome was indexed and constructed using STAR 2.5.2b^97^. Subsequently, the sequencing data was mapped to the indexed genome using the BD Rhapsody single-nucleus pipeline v1.9, employing default parameters.

Each data matrix was converted to a SingleCellExperiment data format, and empty barcodes were removed using the emptyDrops function of DropletUtils v3.14^98^. Then, we converted and processed the data in Seurat v3^28^. We removed cells that had <100 or > 2500 genes and < 100 or > 6000 unique molecular identifiers (UMIs). We also removed genes that had <10 UMIs in each data matrix. Then, we log-normalised the data and used DoubletFinder v2.0^99^ to remove potential doublet, assuming a 7.5% doublet formation rate. The numbers of retained nuclei were 9717, 21614 and 28928 for Fanmao, Starvation and Reconstitution data, respectively (Supplementary Table S20). We used the top 3000 highly variable genes for principal component analysis (PCA) and a reciprocal PCA approach to integrating the three datasets^100^. The first 40 principal components (PCs) were used for uniform manifold approximation and projection (UMAP) dimensional reduction and following clustering (Supplementary Fig. S11). We employed an empirical parameter - a resolution of 0.2 - in the FindClusters function, utilizing the original Louvain algorithm. To identify unique marker genes associated with each cluster, we utilized the FindAllMarkers function from the Seurat package. This analysis employed the Wilcoxon rank sum test, focusing on genes that demonstrated a minimum 0.3-fold difference between two groups of cells. The annotation of cell types relied on the reference of previously published marker genes. In instances where clusters exhibited marker genes that couldn’t be associated with known cell types, we pursued the validation process through whole-mount *in situ* hybridization (WISH; elaborated below), discerning cell types based on gene expression patterns and morphological characteristics. When WISH displayed uniform expression of marker genes from different clusters within the same cell type, we consolidated those clusters into a single cell type. We assigned cells to 14 reliable cell types (Supplementary Fig. S11). Supplementary data S1 presents the counting matrix and Supplementary data S1 presents the average expression of each gene per cell type. To evaluate the stability of each cell type, we implemented a bootstrap sampling and clustering strategy comprising 100 iterations using cells combined from all three samples and individual samples^101^. The determination of marker genes per cell type followed the previously described methodology. The identified marker genes were subsequently utilized in KEGG enrichment analysis, employing ClusterProfiler v4.2^102^.

### Cell trajectory analysis

We conducted slingshot trajectory analyses using Slingshot^103^ v2.2.1 for a subset of clusters (intercalary cells, dorsal-end proliferation cells, ventral-end proliferation cells, mucus cells, basal membrane cells 1, basal membrane cells 2 and bacteriocytes) to explore the developmental trajectory of cells, assuming that all these cells were developed from the same precursor (’posterior-end budding zone’). We performed PCA for the subsampled data set and conducted dimensional reduction using phateR v1.0.7 for slingshot analyses. We used the Potential of Heat-diffusion for Affinity-based Trajectory Embedding (PHATE) because it could better reveal developmental branches than other tools^104^.

We also examined the effect of deep-sea transplant experiments on shaping gene expression patterns by comparing the expression levels amongst the three different states of a given cluster. We conducted Monocle analyses using Monocle2 and Monocle 3 in R environment^105,106^. This comparison was done for the bacteriocytes. The biased number of cells per state could affect the results of the dimensional reduction and calculation of marker genes, and the sequenced nucleus per state was unbalanced; therefore, we first downsampled the cells per cluster per state to to a maximum of 1000 nuclei per cell type. Thereafter, we performed PCA and dimensional reduction using UMAP and PHATE, calculated marker genes and conducted slingshot trajectory and KEGG enrichment analyses as mentioned above for each cell type.

### Phylogenetic estimation

Phylogenetic estimation was conducted for E3 ubiquitin ligase RNF213 genes. We downloaded RNF213 genes from GenBank for representative vertebrate and invertebrate species across the tree of life. The amino acid sequences were aligned with the *G. platifrons* sequences annotated as E3 ubiquitin ligase (Bpl_scaf_25983-1.26 and Bpl_scaf_1894-1.45) using MAFFT v7.450. The alignment was used to estimate a maximum-likelihood best gene tree and to calculate bootstrap values on each node using RAxML v8.2.

### *G. platifrons* gill fixation and storage

The gill tissues of *G. platifrons* collected from Fanmao site were dissected within minutes after the ROV, and samples were retrieved on board R/V Kexue. The gill tissues were briefly washed with ice-cold filtered and autoclaved *in situ* seawater (FAISW) and then fixed in 4% PFA prepared in FAISW at 4 °C overnight. The gill tissues were washed three times with ice-cold PBST, dehydrated in 100% methanol and stored at −20 °C until use.

### Synthesise probes for mRNA *in situ* hybridisations

For WISH and double FISH analyses, the DNA fragments (∼1000bp) of the targeted genes were first PCR amplified with gene-specific primers (GSPs) pairs using *G. platifrons* gill cDNA as template (the sequences of targeted genes and gene-specific primers were provided in Supplementary Data S3). The amplified fragments were ligated into the pMD18-simpleT vector (Takara) and transformed into *E. coli*. Individual colonies were picked up, and their plasmids were sequenced to confirm the inserts. The templates for in vitro mRNA transcription were amplified using T7 forward GSP (sense probe control) or Sp6 reversed GSPs (antisense probe) combined with either forward or reversed gene-specific primer. Labelled probes and control probes were generated using digoxigenin (DIG)-12-UTP (Roche) or fluorescein-12-UTP (Roche) according to the protocol described by Thisse^107^ with Sp6 and T7 RNA polymerase, respectively.

### Paraffin embedding and double fluorescent *in situ* hybridisation

The methanal-dehydrated gill slices were incubated in 100% ethanol, a 1:1 mixture of 100% ethanol and xylene and Xylene twice for 1 h each at RT. The samples were embedded by incubating in Paraplast Plus (Sigma-Aldrich) for 2 h at 65 °C and then cooled down to RT. Sections with 5 μM thickness were cut using a microtome (Leica).

For double FISH, sections were dewaxed by incubating in xylene twice, a 1:1 mixture of 100% ethanol and xylene, 100% ethanol twice, 95% ethanol, 85% ethanol and 75% ethanol for 15 min each at RT. The sections were washed with PBST three times for 10 min each and then permeabilised by 2 μg/mL proteinase K (NEB) in PBST for 15 min at RT. Post-digestion fixation was conducted by incubating the sections in 4% PFA in PBST for 30 min at RT. The sections were washed three times with PBST for 15 min each. Pre-hybridisation was conducted by incubating the sections in HM for 1 h at 55 °C. Then, the *in situ* hybridisation was performed by incubating the sections in ∼0.5 ng/μL fluorescein (Roche)-labelled probed prepared in fresh HM overnight at 55 °C. The sections were washed three times with 2×SSC for 15 min each at 55 °C, cooled down to room temperature and washed three times with PBST.

A second-round hybridisation was conducted on the DIG-labelled oligonucleotides to label the symbiont. The slices were hybridised for 1 h at 46 °C with 100 ng DIG-labelled *G. platifrons* symbiont-specific probe in FISH buffer (0.9 M NaCl, 0.02 M Tris-HCl, 0.01% sodium dodecyl sulphate [SDS] and 30% formamide). The slices were then washed with FISH washing buffer (0.1 M NaCl, 0.02 M Tris HCl, 0.01% SDS and 5 mM EDTA) three times at 5 min each at 48 °C. The slices were washed with PBST three times and then blocked with blocking buffer (2.5% sheep serum and 2% BSA in sheep serum) for 1 h at room temperature.

The slices were then incubated with 1:1000 diluted anti-fluorescein–peroxidase (POD) (Roche) overnight at 4 °C, then washed six times with PBST for 15 min each and three times with TNT buffer (100 mM Tris-HCl, pH 7.5; 100 mM NaCl; 0.1% Tween 20) for 15 min each. Afterwards, the fluorescent signal of the *G. platifrons* gene expression pattern was developed by the TSA fluorescein kit (Akoya Biosciences) according to the manufacturer’s protocol. The slices were washed three times, and the remaining POD activity was quenched by incubation in 1% hydrogen peroxide solution for 1 h at room temperature. Then, the slices were washed three times with PBS, blocked with blocking buffer for 30 min at room temperature, incubated with 1:2500 diluted anti-DIG–POD (Roche) for 2 h at RT and washed with PBST for six times and TNT for three times. The symbiont FISH signal was developed using the TSA Cy3 kit (Akoya Biosciences). Finally, the slices were washed with PBST, stained with DAPI, and mounted with ProLong Diamond Antifade Mountant (Thermo Fisher).

### Whole-mount *in situ* hybridisation

For WISH, the connective region at the end of the W-shaped gill filament was cut off, and each gill slice was carefully peeled off with fine-tip tweezers. We dissected gill tissues from 5 individual mussels and pooled all the gill slices together. The gill slices were then rehydrated in 75%, 50% and 25% methanol-PBST (1×PBS with 0.1% Tween 20) for 15 min each, followed by 3×5 min PBST washes. The gill slices were then permeabilised with 2 μg/mL proteinase K in PBST for 30 min at 37 °C. Post-digestion fixation was conducted by fixing the gill slices with 4% PFA in PBST for 30 min at room temperature (RT). After 3×5 min PBST wash to remove the residual fixative, the gill slices were pre-hybridised with a hybridisation mix (HM; containing 50% formamide, 5×saline–sodium citrate (SSC), 0.1% Tween 20, 10 μg/mL heparin, 500 μg/mL yeast tRNA) for 1 h at 65 °C. For each hybridisation, 5-10 gill slices were added to 400 μL of fresh HM containing ∼0.5 ng/μL DIG-labelled probe. Hybridisation was conducted in a 55 °C shaking water bath overnight. Post-hybridisation washes were performed according to the following steps: the gill slices were first washed 3×15 min with hybridisation washing buffer (50% formamide, 5×SSC, 0.1% Tween 20), followed by 3×15 min 2×SSC with 0.1% Tween 20 and 3×15 min 0.2×SSC with 0.1% Tween 20. The washings were also conducted in a shaking water batch, and all the washing buffers were pre-heated to 55 °C. The samples were washed three times with PBST and then blocked in blocking buffer (2.5% sheep serum, 2% BSA in PBST) for 1 h at RT. Each sample was incubated with 1:10,000 diluted anti-DIG-AP antibody (Roche) at 4 °C overnight. The samples were incubated with the antibody, then washed for 6×15 min with PBST, followed by washing in 3×15 min alkaline Tris buffer (100 mM Tris-HCl, pH 9.5; 100 mM NaCl; 50 mM MgCl_2_). The samples were incubated in nitro blue tetrazolium/5-Bromo-4-chloro-3-indolyl phosphate staining solution (Sangon). After the desired expression pattern was revealed, the staining reaction was stopped by 3×15 min PBST–EDTA wash (PBST, 1 mM ETDA). The gill slices were cleared by incubating in 100% glycerol overnight at 4 °C and then mounted on glass slides. The results of control hybridisations (with sense probes) were provided in Supplementary Fig. S13-S16. WISH analyses were repeated with another batch of *G. platifrons* gill slices samples collected during the *R/V* ‘Kexue’ 2017 South China Sea cold seep cruise to confirm the consistency in expression patterns.

### Microscopy imaging

All the WISH samples and whole-mount 16S FISH images were observed and imaged with a Nikon Eclipse Ni microscope with a DS-Ri2 camera. The double FISH slides were imaged with a Zeiss LSM710 confocal microscope.

### Electron microscopy analysis

The gill slices of the *G. platifrons* were dissected and fixed in electron microscopy fixative (2.5% glutaraldehyde and 2% PFA) at 4 °C. For SEM analysis, the samples were dehydrated in a graded ethanol series and then dried at the critical point. The samples were then coated with gold (sputter/carbon Thread, EM ACE200) and observed under a scanning electron microscope (VEGA3, Tescan). For TEM analysis, the samples were rinsed with double distilled water, post-fixed with 1% osmium tetroxide and then washed with double distilled water. The samples were then rinsed, dehydrated and embedded in Ep812 resin. Ultrathin sections were obtained with an ultramicrotome (70 nm thickness, Reichert-Jung Ultracut E). The sections were then double-stained with lead citrate and uranyl acetate. The cells were observed under a transmission electron microscope (JEM1200, Jeol) operated under 100 kV.

## References

1 Kremer, N. et al. Initial symbiont contact orchestrates host-organ-wide transcriptional changes that prime tissue colonization. Cell Host & Microbe. 14, 183–194 (2013). 10.1016/j.chom.2013.07.006

2 Bang, C. et al. Metaorganisms in extreme environments: do microbes play a role in organismal adaptation? Zoology (Jena) (2018). 10.1016/j.zool.2018.02.004

3 Franke, M., Geier, B., Hammel, J. U., Dubilier, N. & Leisch, N. Coming together-symbiont acquisition and early development in deep-sea bathymodioline mussels. Proc. R. Soc. 288, 20211044 (2021). 10.1098/rspb.2021.1044

4 Dubilier, N., Bergin, C. & Lott, C. Symbiotic diversity in marine animals: the art of harnessing chemosynthesis. Nat. Rev. Microbiol. 6, 725 (2008). 10.1038/nrmicro1992

5 Sogin, E. M., Kleiner, M., Borowski, C., Gruber-Vodicka, H. R. & Dubilier, N. Life in the dark: phylogenetic and physiological diversity of chemosynthetic symbioses. Annu. Rev.Microbiol. 75, 695–718 (2021). 10.1146/annurev-micro-051021-123130

6 Xu, T., Feng, D., Tao, J. & Qiu, J.-W. A new species of deep-sea mussel (Bivalvia: Mytilidae: *Gigantidas*) from the South China Sea: Morphology, phylogenetic position, and gill-associated microbes. Deep Sea Res. Part I Oceanogr. Res. Pap. 146, 79–90 (2019). 10.1016/j.dsr.2019.03.001

7 DeChaine, E. G. & Cavanaugh, C. M. Symbioses of methanotrophs and deep-sea mussels (Mytilidae: Bathymodiolinae). Prog. Mol. Subcell. Biol. 41, 227–249 (2006). 10.1007/3-540-28221-1_11

8 Fujiwara, Y. et al. Phylogenetic characterization of endosymbionts in three hydrothermal vent mussels influence on host distributions. Mar. Ecol. Prog. Ser. 208, 147–155 (2000). doi:10.3354/meps208147

9 Kiel, S. The vent and seep biota : aspects from microbes to ecosystems. (Springer, 2010). 10.1007/978-90-481-9572-5

10 Vrijenhoek, R. C. Genetic diversity and connectivity of deep-sea hydrothermal vent metapopulations. Mol. Ecol. 19, 4391–4411 (2010). 10.1111/j.1365-294X.2010.04789.x

11 Halary, S., Riou, V., Gaill, F., Boudier, T. & Duperron, S. 3D FISH for the quantification of methane- and sulphur-oxidizing endosymbionts in bacteriocytes of the hydrothermal vent mussel *Bathymodiolus azoricus*. ISME J 2, 284–292 (2008). 10.1038/ismej.2008.3

12 Zheng, P. et al. Insights into deep-sea adaptations and host-symbiont interactions: A comparative transcriptome study on Bathymodiolus mussels and their coastal relatives. Mol. Ecol. 26, 5133–5148 (2017). 10.1111/mec.14160

13 Fiala-Médioni, A., Métivier, C., Herry, A. & Le Pennec, M. Ultrastructure of the gill of the hydrothermal-vent mytilid Bathymodiolus sp.. Mar. Biol. 92, 65–72 (1986). 10.1007/BF00392747

14 Wong, Y. H. et al. High-throughput transcriptome sequencing of the cold seep mussel Bathymodiolus platifrons. Sci. Rep. 5, 16597 (2015). 10.1038/srep16597

15 Sun, J. et al. Adaptation to deep-sea chemosynthetic environments as revealed by mussel genomes. Nat. Ecol. Evol. 1, 121 (2017). 10.1038/s41559-017-0121

16 Bettencourt, R. et al. An Insightful Model to Study Innate Immunity and Stress Response in Deep-Sea Vent Animals: Profiling the Mussel *Bathymodiolus azoricus*. Organismal and Molecular Malacology, 8, (2017). 10.5772/68034.

17 Barros, I. et al. Post-capture immune gene expression studies in the deep-sea hydrothermal vent mussel *Bathymodiolus azoricus* acclimatized to atmospheric pressure. Fish Shellfish Immunol 42, 159–170 (2015). 10.1016/j.fsi.2014.10.018

18 Wang, H. et al. Molecular analyses of the gill symbiosis of the bathymodiolin mussel *Gigantidas platifrons*. iScience 24, 101894 (2021). 10.1016/j.isci.2020.101894

19 Chen, K. H., Boettiger, A. N., Moffitt, J. R., Wang, S. & Zhuang, X. RNA imaging. Spatially resolved, highly multiplexed RNA profiling in single cells. Science 348, 6090 (2015). 10.1126/science.aaa6090

20 Hwang, B., Lee, J. H. & Bang, D. Single-cell RNA sequencing technologies and bioinformatics pipelines. Exp. Mol. Med. 50, 1–14 (2018). 10.1038/s12276-018-0071-8

21 Saliba, A. E., Westermann, A. J., Gorski, S. A. & Vogel, J. Single-cell RNA-seq: advances and future challenges. Nucleic Acids Res 42, 8845–8860 (2014). 10.1093/nar/gku555

22 Chen, X., Teichmann, S. A. & Meyer, K. B. From tissues to cell types and back: single-cell gene expression analysis of tissue architecture. Annu. Rev. Biomed. Data Sci. 1, 29–51 (2018). 10.1146/annurev-biodatasci-080917-013452

23 Wu, H., Kirita, Y., Donnelly, E. L. & Humphreys, B. D. Advantages of single-nucleus over single-cell RNA sequencing of adult kidney: rare cell types and novel cell states revealed in fibrosis. J. Am. Soc. Nephrol. 30, 23–32 (2019). 10.1681/ASN.2018090912

24 Feng, D. et al. Using Bathymodiolus tissue stable carbon, nitrogen and sulfur isotopes to infer biogeochemical process at a cold seep in the South China Sea. Deep Sea Res. Part I Oceanogr. Res. 104, 52–59 (2015). 10.1016/j.dsr.2015.06.011

25 Kharchenko, P. V., Silberstein, L. & Scadden, D. T. Bayesian approach to single-cell differential expression analysis. Nat. Methods. 11, 740–742 (2014). 10.1038/nmeth.2967

26 Elyanow, R., Dumitrascu, B., Engelhardt, B. E. & Raphael, B. J. netNMF-sc: leveraging gene-gene interactions for imputation and dimensionality reduction in single-cell expression analysis. Genome. Res. 30, 195–204 (2020). 10.1101/gr.251603.119

27 Hao, Y. et al. Integrated analysis of multimodal single-cell data. Cell 184, 3573–3587 (2021). 10.1016/j.cell.2021.04.048

28 Stuart, T. et al. Comprehensive integration of single-cell data. Cell 177, 1888–1902 (2019). 10.1016/j.cell.2019.05.031

29 Petibon, C., Malik Ghulam, M., Catala, M. & Abou Elela, S. Regulation of ribosomal protein genes: An ordered anarchy. WIREs RNA 12, e1632 (2021). 10.1002/wrna.1632

30 Welcker, D. et al. Hemicentin-1 is an essential extracellular matrix component of the dermal–epidermal and myotendinous junctions. Sci. Rep. 11, 17926 (2021). 10.1038/s41598-021-96824-4

31 Dufour, S. C. & Beninger, P. G. A functional interpretation of cilia and mucocyte distributions on the abfrontal surface of bivalve gills. Mar. Biol. 138, 295–309 (2001). 10.1007/s002270000466

32 Dufour, S. C. Gill anatomy and the evolution of symbiosis in the bivalve family Thyasiridae. Biol. Bull. 208, 200–212 (2005). 10.2307/3593152

33 Gómez-Mendikute, A., Elizondo, M., Venier, P. & Cajaraville, M. P. Characterization of mussel gill cells in vivo and in vitro. Cell Tissue Res. 321, 131–140 (2005). 10.1007/s00441-005-1093-9

34 Beninger, P. G. & Dufour, S. Mucocyte distribution and relationship to particle transport on the pseudolamellibranch gill of *Crassostrea virginica* (Bivalvia:Ostreidae). Mar. Ecol. Progr. Ser. 137, 133–138 (1996). 10.3354/meps137133

35 Gerdol, M. et al. The C1q domain containing proteins of the Mediterranean mussel Mytilus galloprovincialis: A widespread and diverse family of immune-related molecules. Dev. Comp. Immunol. 35, 635–643 (2011). 10.1016/j.dci.2011.01.018

36 Wang, W., Song, X., Wang, L. & Song, L. Pathogen-derived carbohydrate recognition in molluscs immune defense. Int. J. Mol. Sci. 19, 721 (2018). 10.3390/ijms19030721

37 Saco, A., Rey-Campos, M., Novoa, B. & Figueras, A. Transcriptomic response of mussel gills after a *Vibrio splendidus* infection demonstrates their role in the immune response. Front. Immunol. 11, 615580 (2020). 10.3389/fimmu.2020.615580

38 Richoux, N. B. & Thompson, R. J. Regulation of particle transport within the ventral groove of the mussel (*Mytilus edulis*) gill in response to environmental conditions. J. Exp. Mar. Biol. Ecol. 260, 199–215 (2001). 10.1016/s0022-0981(01)00254-4

39 Riisgård, H. U., Egede, P. P. & Barreiro Saavedra, I. Feeding behaviour of the mussel, *Mytilus edulis*: New observations, with a minireview of current knowledge. J. Mar. Biol. 2011, 312459 (2011). 10.1155/2011/312459

40 Heude, É., Shaikho, S. & Ekker, M. The dlx5a/dlx6a genes play essential roles in the early development of zebrafish median fin and pectoral structures. PLOS ONE 9, e98505 (2014). 10.1371/journal.pone.0098505

41 Han, S. et al. TULP3 is required for localization of membrane-associated proteins ARL13B and INPP5E to primary cilia. Biochem. Biophys. Res. Commun. 509, 227–234 (2019). 10.1016/j.bbrc.2018.12.109

42 Louka, P. et al. Proteins that control the geometry of microtubules at the ends of cilia. J. Cell Biol. 217, 4298–4313 (2018). 10.1083/jcb.201804141

43 Das, A., Dickinson, D. J., Wood, C. C., Goldstein, B. & Slep, K. C. Crescerin uses a TOG domain array to regulate microtubules in the primary cilium. Mol. Biol. Cell. 26, 4248–4264 (2015). 10.1091/mbc.E15-08-0603

44 Llorente-Cortes, V., Martinez-Gonzalez, J. & Badimon, L. LDL receptor-related protein mediates uptake of aggregated LDL in human vascular smooth muscle cells. Arterioscler. Thromb. Vasc. Biol. 20, 1572–1579 (2000). 10.1161/01.atv.20.6.1572

45 Chen, X. et al. Angiotensin-converting enzyme in smooth muscle cells promotes atherosclerosis-brief report. Arterioscler. Thromb. Vasc. Biol. 36, 1085–1089 (2016). 10.1161/ATVBAHA.115.307038

46 Linke, W. A. & Grützner, A. Pulling single molecules of titin by AFM—recent advances and physiological implications. Pflug. Arch. Eur. 456, 101–115 (2008). 10.1007/s00424-007-0389-x

47 St Paul, A., Corbett, C. B., Okune, R. & Autieri, M. V. Angiotensin II, Hypercholesterolemia, and vascular smooth muscle cells: A perfect trio for vascular pathology. Int. J. Mol. Sci. 21 4525 (2020). 10.3390/ijms21124525

48 Ytrehus, K., Ludvigsen, S., Mancusi, C., Gerdts, E. & de Simone, G. Heart angiotensin-converting enzyme and angiotensin-converting enzyme 2 gene expression associated with male sex and salt-sensitive hypertension in the Dahl Rat. Front. physiol. 12 (2021). 10.3389/fphys.2021.663819

49 Keller, T. C., 3rd et al. Role of titin in nonmuscle and smooth muscle cells. Adv. Exp. Med. Biol. 481, 265–277 (2000). 10.1007/978-1-4615-4267-4_16

50 Chen, J. V. et al. Rootletin organizes the ciliary rootlet to achieve neuron sensory function in *Drosophila*. J. Cell. Biol. 211, 435–453 (2015). 10.1083/jcb.201502032

51 Styczynska-Soczka, K. & Jarman, A. P. The *Drosophila* homologue of rootletin is required for mechanosensory function and ciliary rootlet formation in chordotonal sensory neurons. Cilia 4, 9 (2015). 10.1186/s13630-015-0018-9

52 Mohan, S., Timbers, T. A., Kennedy, J., Blacque, O. E. & Leroux, M. R. Striated rootlet and nonfilamentous forms of rootletin maintain ciliary function. Curr. Biol. 23, 2016–2022 (2013). 10.1016/j.cub.2013.08.033

53 Jones, S. E. & Jomary, C. Secreted Frizzled-related proteins: searching for relationships and patterns. Bioessays 24, 811–820 (2002). 10.1002/bies.10136

54 Gillis, W. Q., Bowerman, B. A. & Schneider, S. Q. The evolution of protostome GATA factors: molecular phylogenetics, synteny, and intron/exon structure reveal orthologous relationships. BMC Evol. Biol. 8, 112–112 (2008). 10.1186/1471-2148-8-112

55 Zhang, Z., Liu, L., Twumasi-Boateng, K., Block, D. H. S. & Shapira, M. FOS-1 functions as a transcriptional activator downstream of the *C. elegans* JNK homolog KGB-1. Cell. Signal. 30, 1–8 (2017). 10.1016/j.cellsig.2016.11.010

56 Phochanukul, N. & Russell, S. No backbone but lots of Sox: Invertebrate Sox genes. Int. J. Biochem. Cell Biol. 42, 453–464 (2010). 10.1016/j.biocel.2009.06.013

57 Cannuel, R., Beninger, P. G., McCombie, H. & Boudry, P. Gill Development and its functional and evolutionary implications in the blue mussel *Mytilus edulis* (Bivalvia: Mytilidae). Biol. Bull. 217, 173–188 (2009). 10.1086/BBLv217n2p173

58 Leibson, N. L. & Movchan, O. T. Cambial zones in gills of Bivalvia. Mar. Biol. 31, 175–180 (1975). 10.1007/BF00391629

59 Wentrup, C., Wendeberg, A., Schimak, M., Borowski, C. & Dubilier, N. Forever competent: deep-sea bivalves are colonized by their chemosynthetic symbionts throughout their lifetime. Environ. Microbiol. 16, 3699–3713 (2014). 10.1111/1462-2920.12597

60 Mohieldin, A. M. et al. Proteomic identification reveals the role of ciliary extracellular-like vesicle in cardiovascular function. Adv. Sci. 7, 1903140 (2020). 10.1002/advs.201903140

61 Neumann, D. & Kappes, H. On the growth of bivalve gills initiated from a lobule-producing budding zone. Biol. Bull. 205, 73–82 (2003). 10.2307/1543447

62 Piquet, B. et al. Regionalized cell proliferation in the symbiont-bearing gill of the hydrothermal vent mussel Bathymodiolus azoricus. Symbiosis 82, 225–233 (2020). 10.1007/s13199-020-00720-w

63 Streams, M. E., Fisher, C. R. & Fiala-Médioni, A. Methanotrophic symbiont location and fate of carbon incorporated from methane in a hydrocarbon seep mussel. Mar. Biol. 129, 465–476 (1997). 10.1007/s002270050187

64 Ponnudurai, R. et al. Metabolic and physiological interdependencies in the *Bathymodiolus azoricus* symbiosis. ISME J 11, 463–477 (2017). 10.1038/ismej.2016.124

65 Barry, J. P. et al. Methane-based symbiosis in a mussel, *Bathymodiolus platifrons*, from cold seeps in Sagami Bay, Japan. Invertebr. Biol. 121, 47–54 (2002). doi:10.1111/j.1744-7410.2002.tb00128.x

66 Renoz, F., Noel, C., Errachid, A., Foray, V. & Hance, T. Infection dynamic of symbiotic bacteria in the pea *Aphid acyrthosiphon pisum* gut and host immune response at the early steps in the infection process. Plos One 10, 0122099 (2015). ARTNe012209910.1371/journal.pone.

67 Guan, H. et al. Lipid Biomarker Patterns Reflect Nutritional Strategies of Seep-Dwelling Bivalves From the South China Sea. Front. Mar. Sci. 9, 3398 (2022). 10.3389/fmars.2022.831286

68 Li, Y. et al. Genomic adaptations to chemosymbiosis in the deep-sea seep-dwelling tubeworm *Lamellibrachia luymesi*. BMC Biol. 17, 91 (2019). 10.1186/s12915-019-0713-x

69 Sun, Y. et al. Genomic signatures supporting the symbiosis and formation of chitinous tube in the deep-sea tubeworm *Paraescarpia echinospica*. Mol. Biol. Evol. 38, 4116–4134 (2021). 10.1093/molbev/msab203

70 Takishita, K. et al. Genomic evidence that methanotrophic endosymbionts likely provide deep-sea bathymodiolus mussels with a sterol intermediate in cholesterol biosynthesis. Genome Biol. Evol. 9, 1148–1160 (2017). 10.1093/gbe/evx082

71 Demidenko, A., Akberdin, I. R., Allemann, M., Allen, E. E. & Kalyuzhnaya, M. G. Fatty acid biosynthesis pathways in *Methylomicrobium buryatense* 5G(B1). Front. Microbiol. 7, 2167 (2016). 10.3389/fmicb.2016.02167

72 Brasaemle, D. L. et al. Adipose differentiation-related protein is an ubiquitously expressed lipid storage droplet-associated protein. J. Lipid Res. 38, 2249–2263 (1997). 10.1016/S0022-2275(20)34939-7

73 Cheng, D. et al. Expression, purification, and characterization of human and rat acetyl coenzyme A carboxylase (ACC) isozymes. Protein Expr. Purif. 51, 11–21 (2007). 10.1016/j.pep.2006.06.005

74 Monroig, Ó. & Kabeya, N. Desaturases and elongases involved in polyunsaturated fatty acid biosynthesis in aquatic invertebrates: a comprehensive review. Fish. Sci. 84, 911–928 (2018). 10.1007/s12562-018-1254-x

75 Kabeya, N. et al. Unique fatty acid desaturase capacities uncovered in Hediste diversicolor illustrate the roles of aquatic invertebrates in trophic upgrading. Philos. Trans. R. Soc. 375, 20190654 (2020). 10.1098/rstb.2019.0654

76 Soupene, E. & Kuypers, F. A. Mammalian long-chain acyl-CoA synthetases. Exp. Biol. Med. (Maywood). 233, 507–521 (2008). 10.3181/0710-MR-287

77 Alves-Bezerra, M. et al. Long-chain acyl-CoA synthetase 2 knockdown leads to decreased fatty acid oxidation in fat body and reduced reproductive capacity in the insect *Rhodnius prolixus*. Biochim. Biophys. Acta. 1861, 650–662 (2016). 10.1016/j.bbalip.2016.04.007

78 Hoglund, P. J., Nordstrom, K. J. V., Schioth, H. B. & Fredriksson, R. The solute carrier families have a remarkably long evolutionary history with the majority of the human families present before divergence of bilaterian species. Mol. Biol. Evol. 28, 1531–1541 (2011). 10.1093/molbev/msq350

79 Mohamed, A. R. et al. Dual RNA-sequencing analyses of a coral and its native symbiont during the establishment of symbiosis. Mol. Ecol. 29, 3921–3937 (2020). 10.1111/mec.15612

80 Bertucci, A., Foret, S., Ball, E. E. & Miller, D. J. Transcriptomic differences between day and night in *Acropora millepora* provide new insights into metabolite exchange and light-enhanced calcification in corals. Mol. Ecol. 24, 4489–4504 (2015). 10.1111/mec.13328

81 Hamada, M. et al. Metabolic co-dependence drives the evolutionarily ancient Hydra-Chlorella symbiosis. Elife 7, e35122 (2018). 10.7554/eLife.35122

82 Duncan, R. P., Feng, H., Nguyen, D. M. & Wilson, A. C. Gene family expansions in aphids maintained by endosymbiotic and nonsymbiotic traits. Genome Biol. Evol. 8, 753–764 (2016). 10.1093/gbe/evw020

83 Feng, H. L. et al. Trading amino acids at the aphid-Buchnera symbiotic interface. P. Natl. Acad. Sci. USA. 116, 16003–16011 (2019). 10.1073/pnas.1906223116

84 Ip, J. C. et al. Host-endosymbiont genome integration in a deep-sea chemosymbiotic clam. Mol. Biol. Evol. 38, 502–518 (2021). 10.1093/molbev/msaa241

85 Hongo, Y. et al. Expression of genes involved in the uptake of inorganic carbon in the gill of a deep-sea vesicomyid clam harboring intracellular thioautotrophic bacteria. Gene 585, 228–240 (2016). 10.1016/j.gene.2016.03.033

86 Sotiriou, S. et al. Ascorbic-acid transporter Slc23a1 is essential for vitamin C transport into the brain and for perinatal survival. Nat. Med. 8, 514–517 (2002). 10.1038/0502-514

87 Aydemir, T. B. et al. Zinc transporter ZIP14 functions in hepatic zinc, iron and glucose homeostasis during the innate immune response (endotoxemia). PLoS One 7, e48679 (2012). 10.1371/journal.pone.0048679

88 Turi, Z., Lacey, M., Mistrik, M. & Moudry, P. Impaired ribosome biogenesis: mechanisms and relevance to cancer and aging. Aging (Albany NY) 11, 2512–2540 (2019). 10.18632/aging.101922

89 Hay, B. A. Understanding IAP function and regulation: a view from Drosophila. Cell. Death. Differ. 7, 1045–1056 (2000). 10.1038/sj.cdd.4400765

90 Dubrez-Daloz, L., Dupoux, A. & Cartier, J. IAPS : More than just inhibitors of apoptosis proteins. Cell Cycle 7, 1036–1046 (2008). 10.4161/cc.7.8.5783

91 Pflug, K. M. & Sitcheran, R. Targeting NF-κB-inducing kinase (NIK) in immunity, inflammation, and cancer. Int. J. Mol. Sci. 21, 8470 (2020).

92 Otten, E. G. et al. Ubiquitylation of lipopolysaccharide by RNF213 during bacterial infection. Nature 594, 111–116 (2021). 10.1038/s41586-021-03566-4

93 Price, D. R. et al. Aphid amino acid transporter regulates glutamine supply to intracellular bacterial symbionts. Proc. Natl. Acad. Sci. USA. 111, 320–325 (2014). 10.1073/pnas.1306068111

94 Ponnudurai, R. et al. Comparative proteomics of related symbiotic mussel species reveals high variability of host–symbiont interactions. ISME J. 14, 649–656 (2020). 10.1038/s41396-019-0517-6

95 Tashian, R. E. The carbonic anhydrases: widening perspectives on their evolution, expression and function. Bioessays 10, 186–192 (1989). 10.1002/bies.950100603

96 Hicks, S. C., Townes, F. W., Teng, M. & Irizarry, R. A. Missing data and technical variability in single-cell RNA-sequencing experiments. Biostatistics 19, 562–578 (2017). 10.1093/biostatistics/kxx053

97 Dobin, A. et al. STAR: ultrafast universal RNA-seq aligner. Bioinformatics 29, 15–21 (2013). 10.1093/bioinformatics/bts635

98 Lun, A. T. L. et al. EmptyDrops: distinguishing cells from empty droplets in droplet-based single-cell RNA sequencing data. Genome Biol. 20, 63 (2019). 10.1186/s13059-019-1662-y

99 McGinnis, C. S., Murrow, L. M. & Gartner, Z. J. DoubletFinder: Doublet detection in single-cell RNA sequencing data using artificial nearest neighbors. Cell Systems 8, 329–337 (2019). 10.1016/j.cels.2019.03.003

100 Stuart, T. et al. Comprehensive Integration of Single-Cell Data. Cell 177, 1888–1902.e1821 (2019). 10.1016/j.cell.2019.05.031

101 Singh, P. & Zhai, Y. Deciphering Hematopoiesis at single cell level through the lens of reduced dimensions. bioRxiv, 2022.2006.2007.495099 (2022). 10.1101/2022.06.07.495099

102 Wu, T. et al. clusterProfiler 4.0: A universal enrichment tool for interpreting omics data. The Innovation 2, 100141 (2021). 10.1016/j.xinn.2021.100141

103 Street, K. et al. Slingshot: cell lineage and pseudotime inference for single-cell transcriptomics. BMC Genom. 19, 477 (2018). 10.1186/s12864-018-4772-0

104 Moon, K. R. et al. Visualizing structure and transitions in high-dimensional biological data. Nat. Biotechnol. 37, 1482–1492 (2019). 10.1038/s41587-019-0336-3

105 Qiu, X. et al. Reversed graph embedding resolves complex single-cell trajectories. Nat Methods 14, 979–982 (2017). 10.1038/nmeth.4402

106 Cao, J. et al. The single-cell transcriptional landscape of mammalian organogenesis. Nature 566, 496–502 (2019). 10.1038/s41586-019-0969-x

107 Thisse, C. & Thisse, B. High-resolution in situ hybridization to whole-mount zebrafish embryos. Nat. Protoc. 3, 59–69 (2008). 10.1038/nprot.2007.514

