## Supplementary Figures for "Deciphering deep-sea chemosynthetic symbiosis by single-nucleus RNA-sequencing"

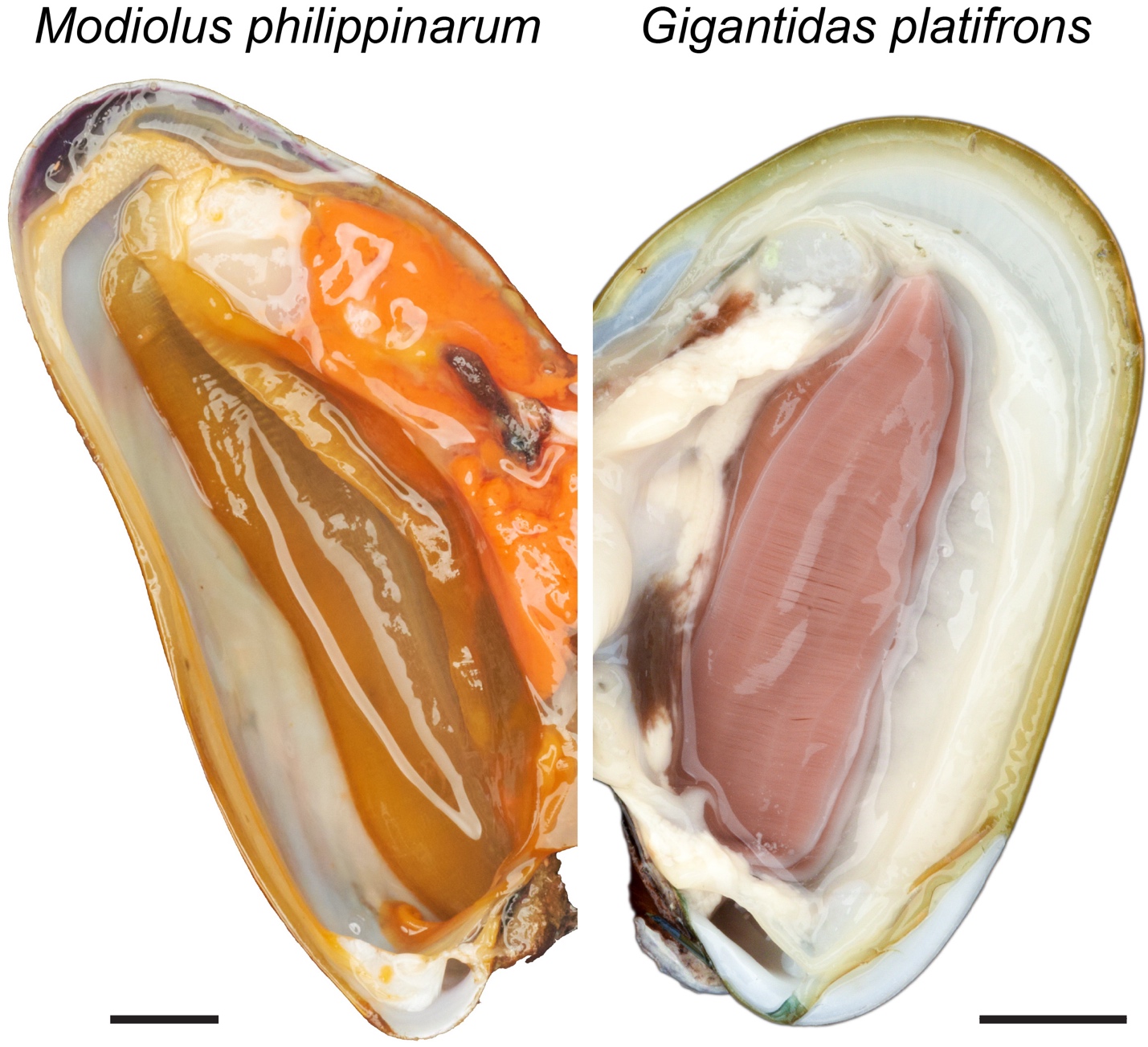

**Supplementary Figure S1:** Image showing the gill tissues of the coastal shallow water mussel *Modiolus philippinarum* and deep-sea chemosynthetic mussel *Gigantidas platifrons*. Comparing with the thin gill tissue of *M. philippinarum*, the gill filaments of *G. platifrons* are noticeably expanded. (Scale bar = 1cm)

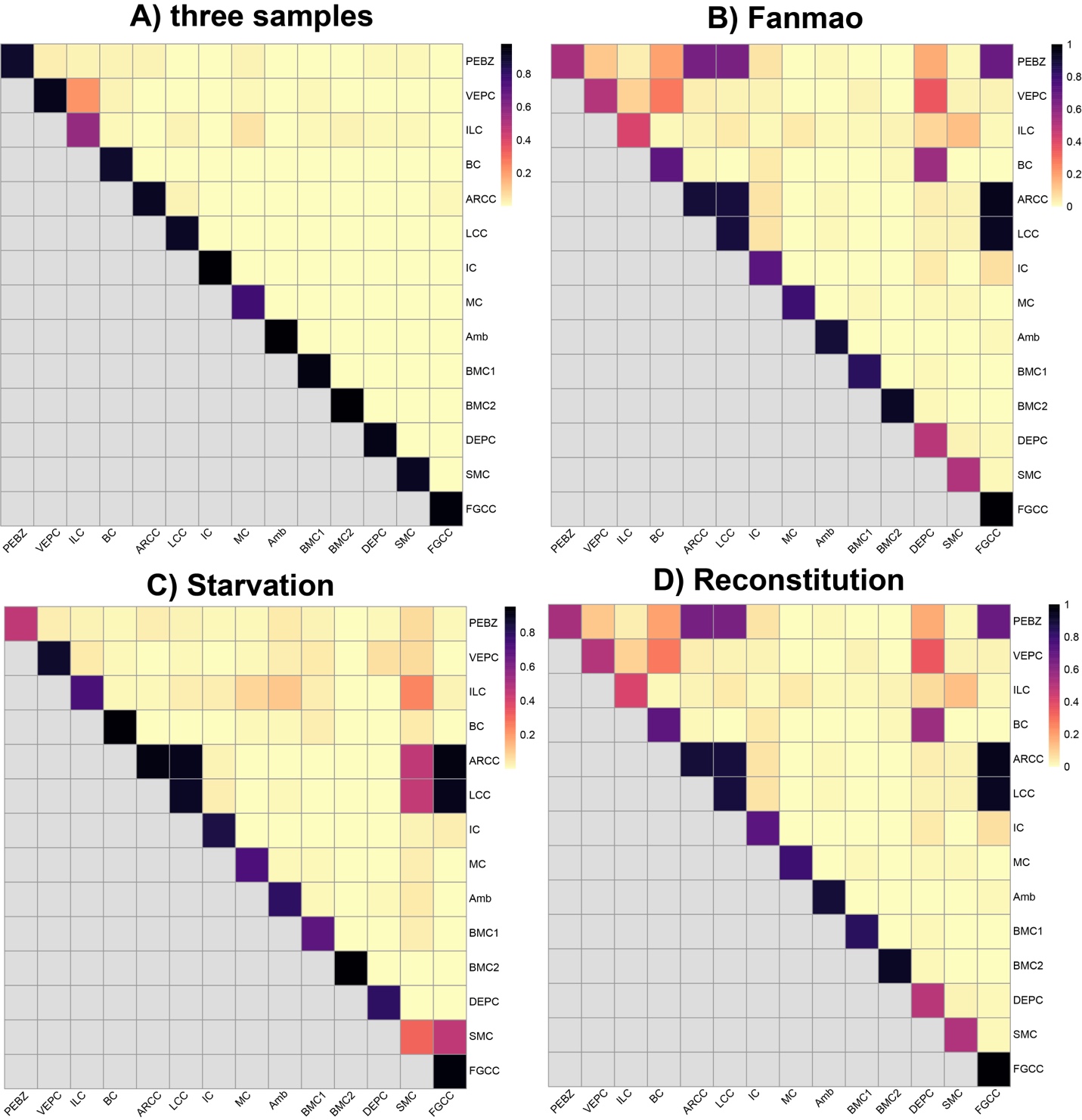

**Supplementary Figure S2:** Heatmaps of co-assignment probabilities of bootstrap sampling estimated for recognised clusters using cells in all three samples (a) and using cells in individual sample of Fanmao (b), Starvation (c) and Reconstitution (d). PEBZ: Posterior end budding zone cell; VEPC: Ventral end proliferation cell; ILC: inter lamina cell, BC: Bacteriocyte; ARCC: Apical ridge ciliary cell; LCC: lateral ciliary cell; IC: Intercalary cell; MC: Mucus cell; Amb: Ambiguous cell; BMC1: Basal membrane cell 1; BMC2: Basal membrane cell 2, DEPC: Dorsal end proliferation cell; SMC: Smooth muscle cell; FGCC: Food groove ciliary cell.

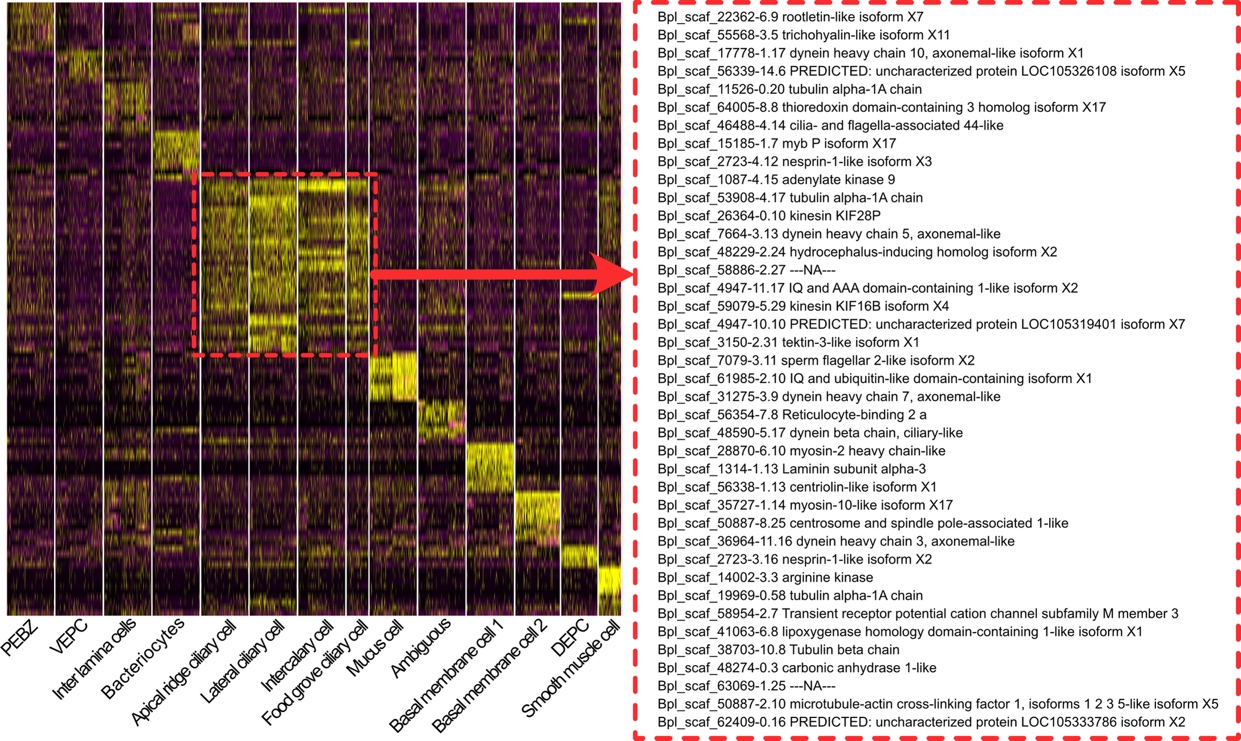

**Supplementary Figure S3:** Heatmap of the shared marker genes expressed in all the ciliary cells.

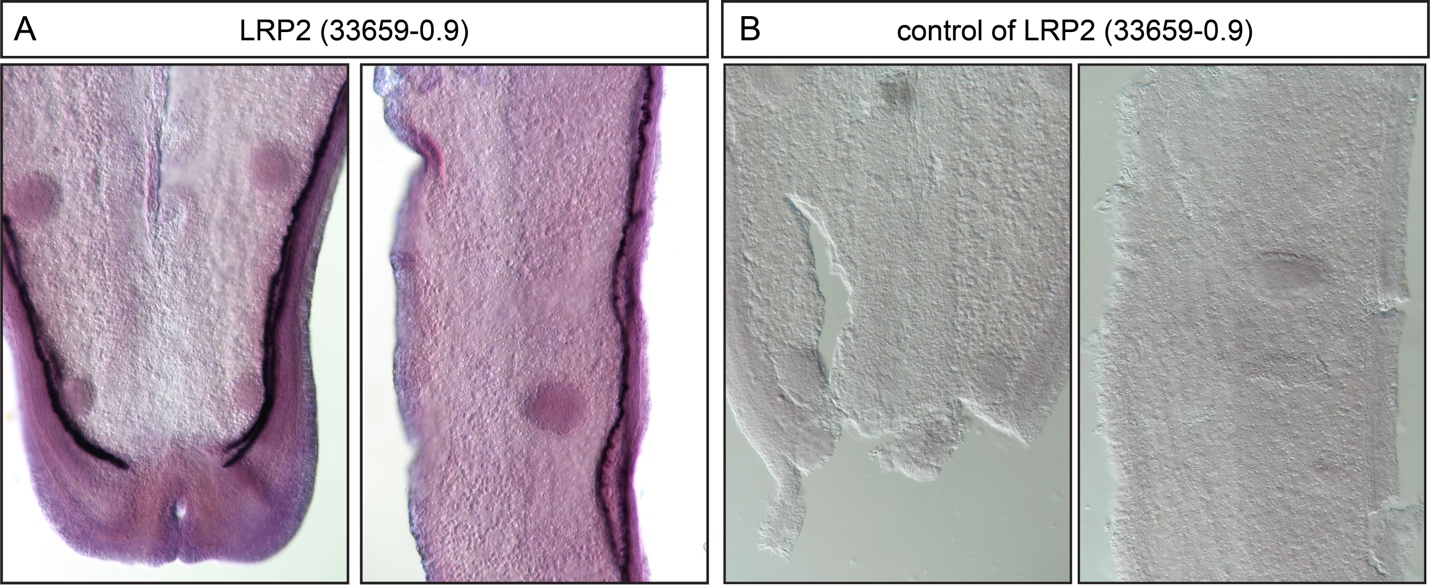

**Supplementary Figure S4:** WISH characterisation of the gene Bpl_scaf_48274-0.3, LRR2. The left and right images of each pannel are the tip and mid part of a gill slice, respectively.

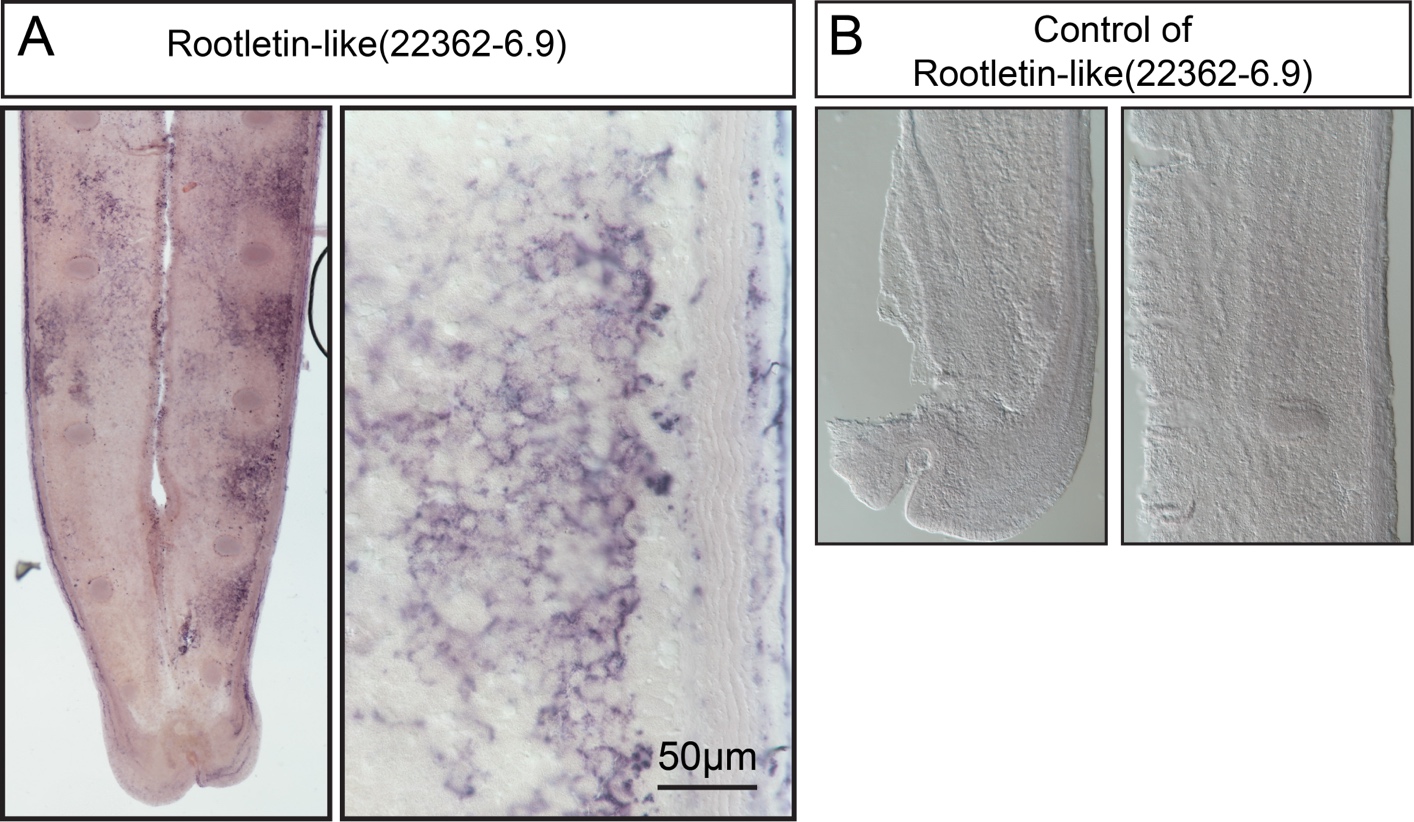

**Supplementary Figure S5:** WISH characterisation of gene Bpl_scaf_22362-6.9, Rootletin-like. The left and right images of each pannel are the tip and mid part of a gill slice, respectively.

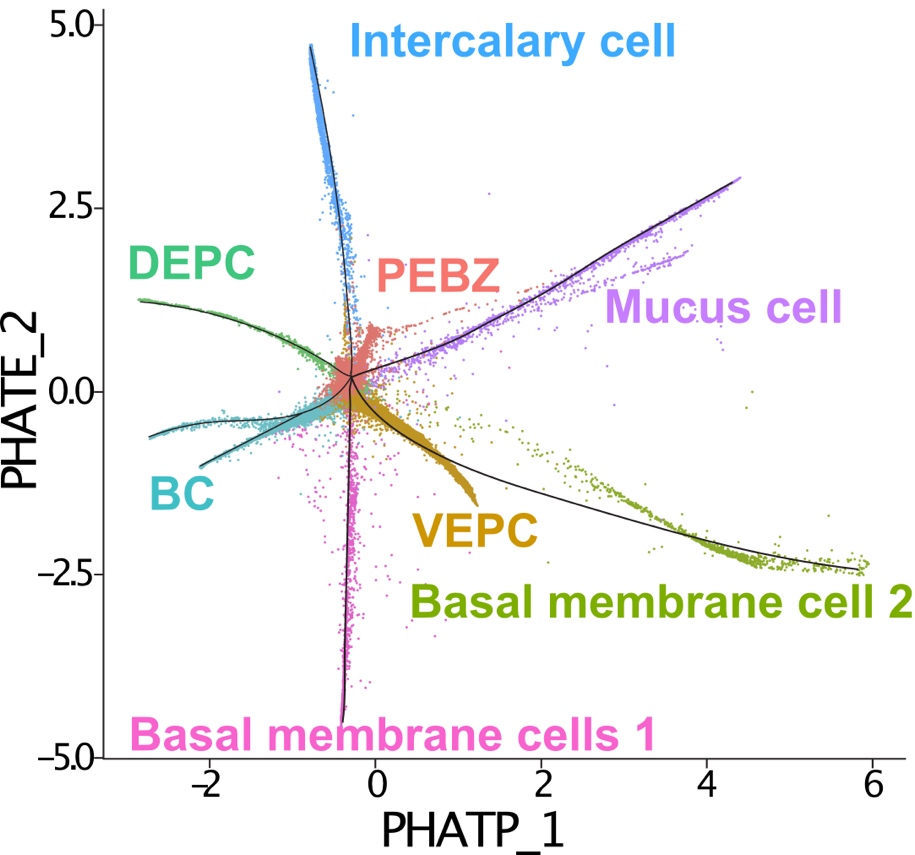

**Supplementary Figure S6:** Slingshot trajectory of selected cell types (a dot represents a cell) identified by slingshot and Phate. The selected cell types include: "Posterior end budding zone", "Intercalary cells","Dosal end proliferation cells","Vential end proliferation cells", "Mucus cells","Basal membrane cells 1", "Basal membrane cells 2", and the "bacteriocytes".

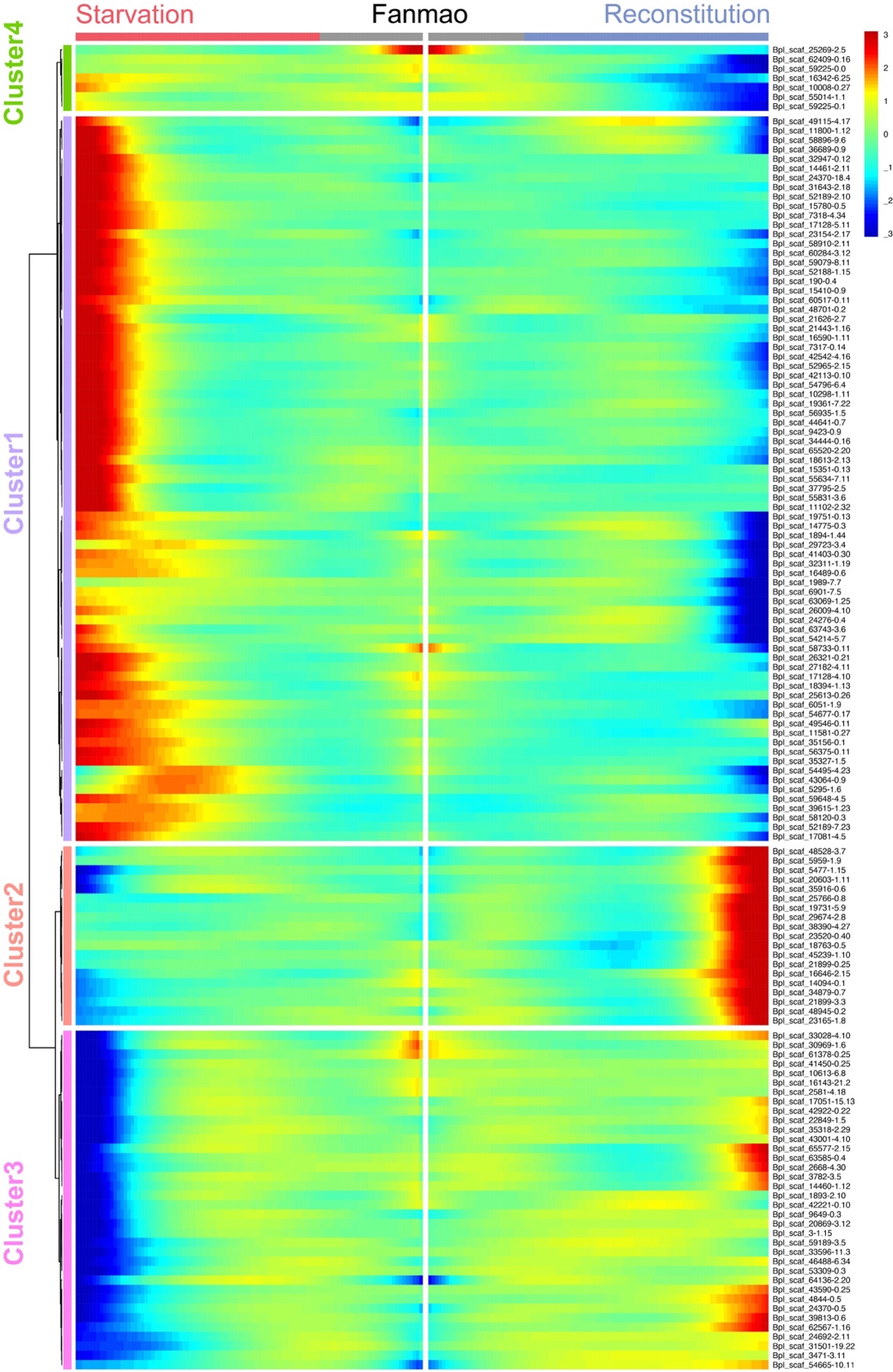

**Supplementary Figure S7:** Heat map showing the gene expression profiles of all bacteriocytes' DEGs.

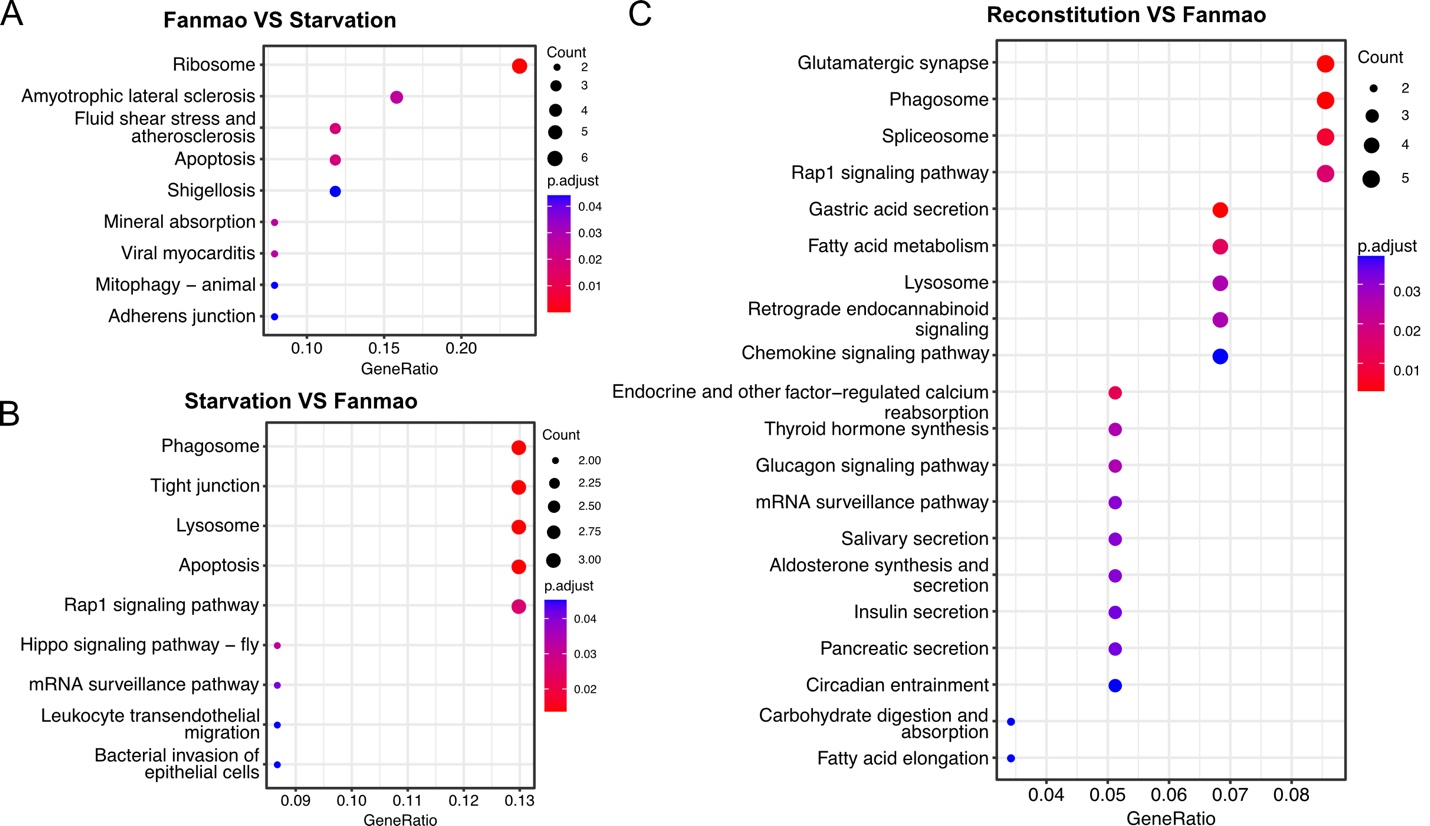

**Supplementary Figure S8:** KEGG enrichment analysis of the bacteriocytes’ DEGs.

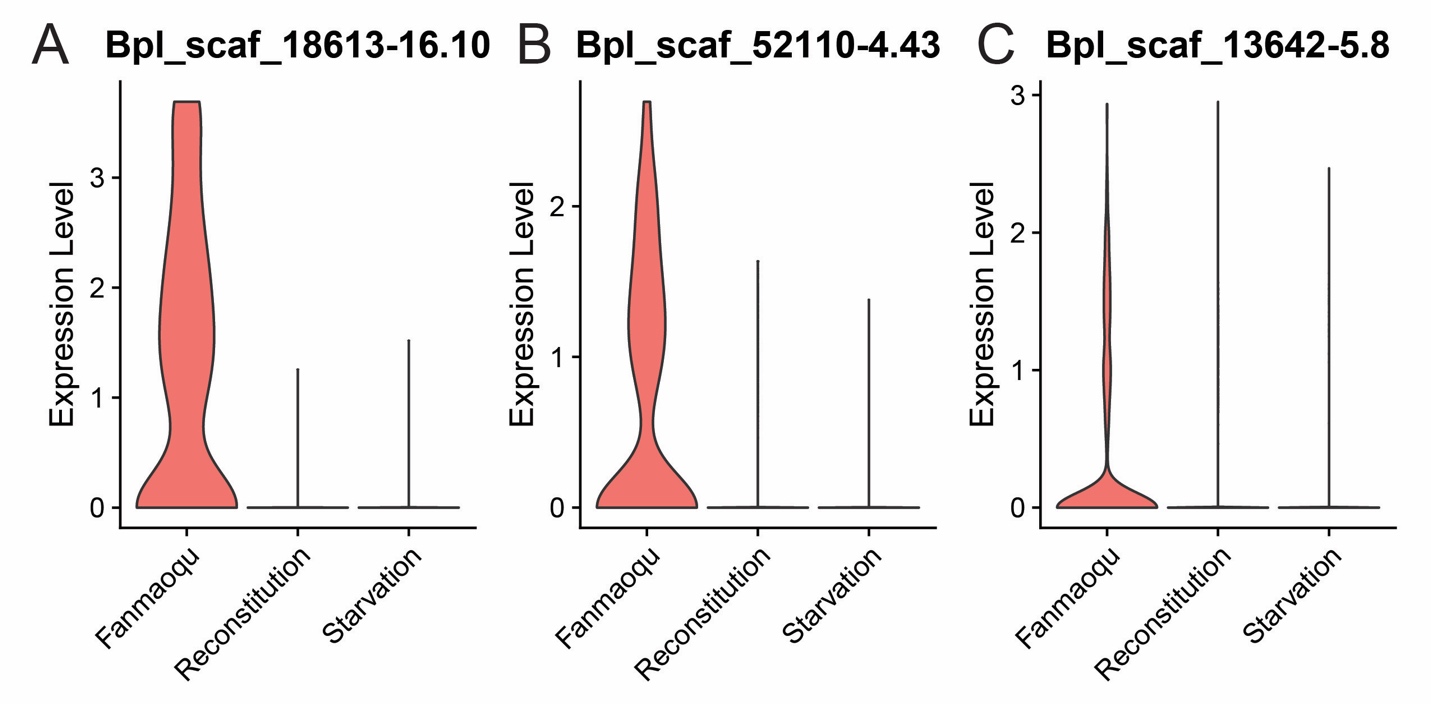

**Supplementary Figure S9:** Gene expression level analysis of selected genes. A: Bpl_Scaf_18613-16.10 (A BCB P-glycol); B: Bpl_scaf_52110-4.43 (canalicular multispecific organic anion transporter 2); and C: Bpl_scaf_13642-5.8 (Sodium- and chloride-dependent glycine transporter 1-like protein)

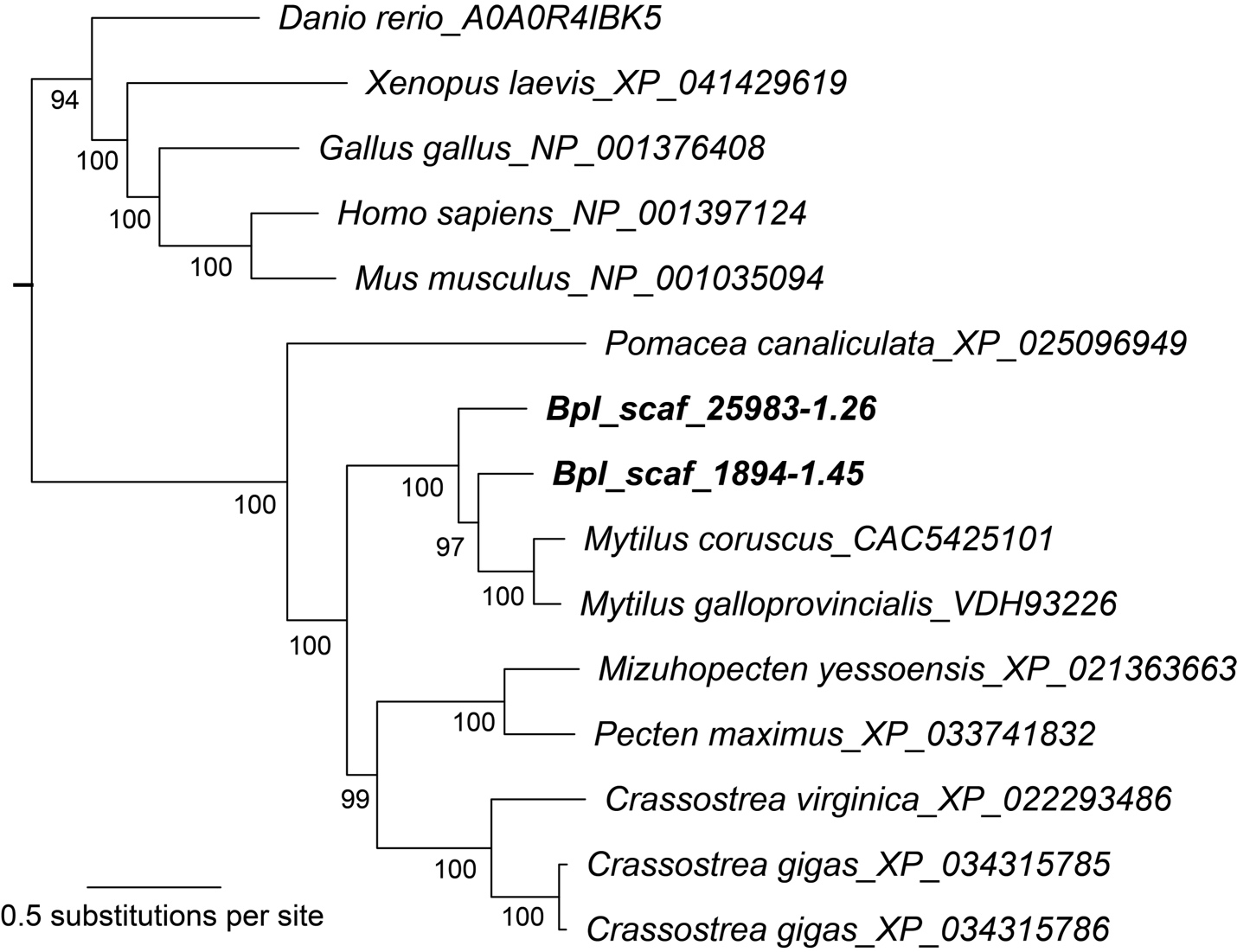

**Supplementary Figure S10:** A RAxML phylogenetic tree estimated using E3 ubiquitin ligase RNF213. Tips are species names and GenBank accession numbers. Node numbers are bootstrap support values.

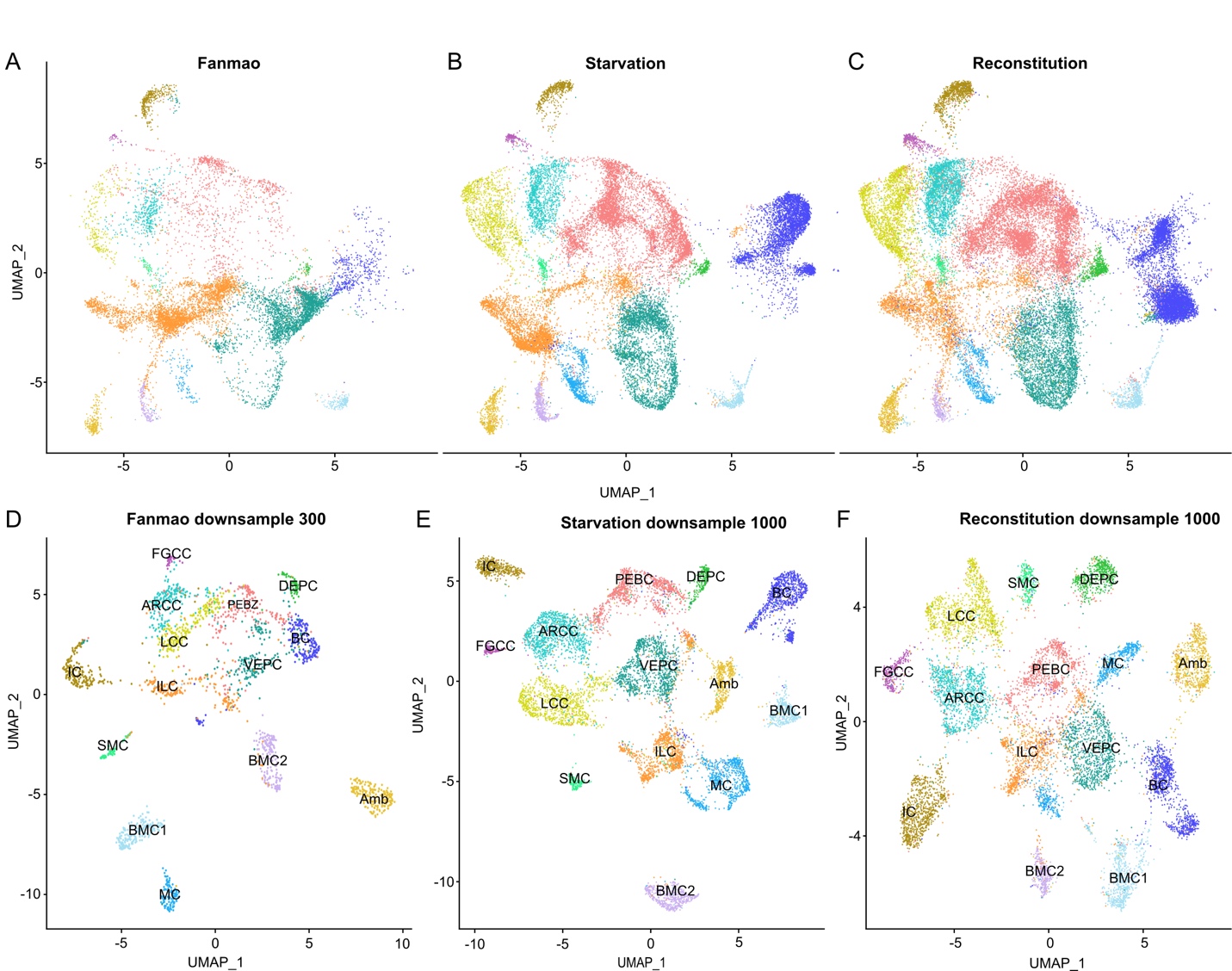

**Supplementary Figure S11:** UMAP plots showing distribution patterns of 14 clusters in Fanmao, Starvation and Reconstitution states. A-C show a representation of cells per cluster in each individual state when analysing all three samples together; D-F shows a representation of cells in each cluster when analysing UMAP for each sample individually.

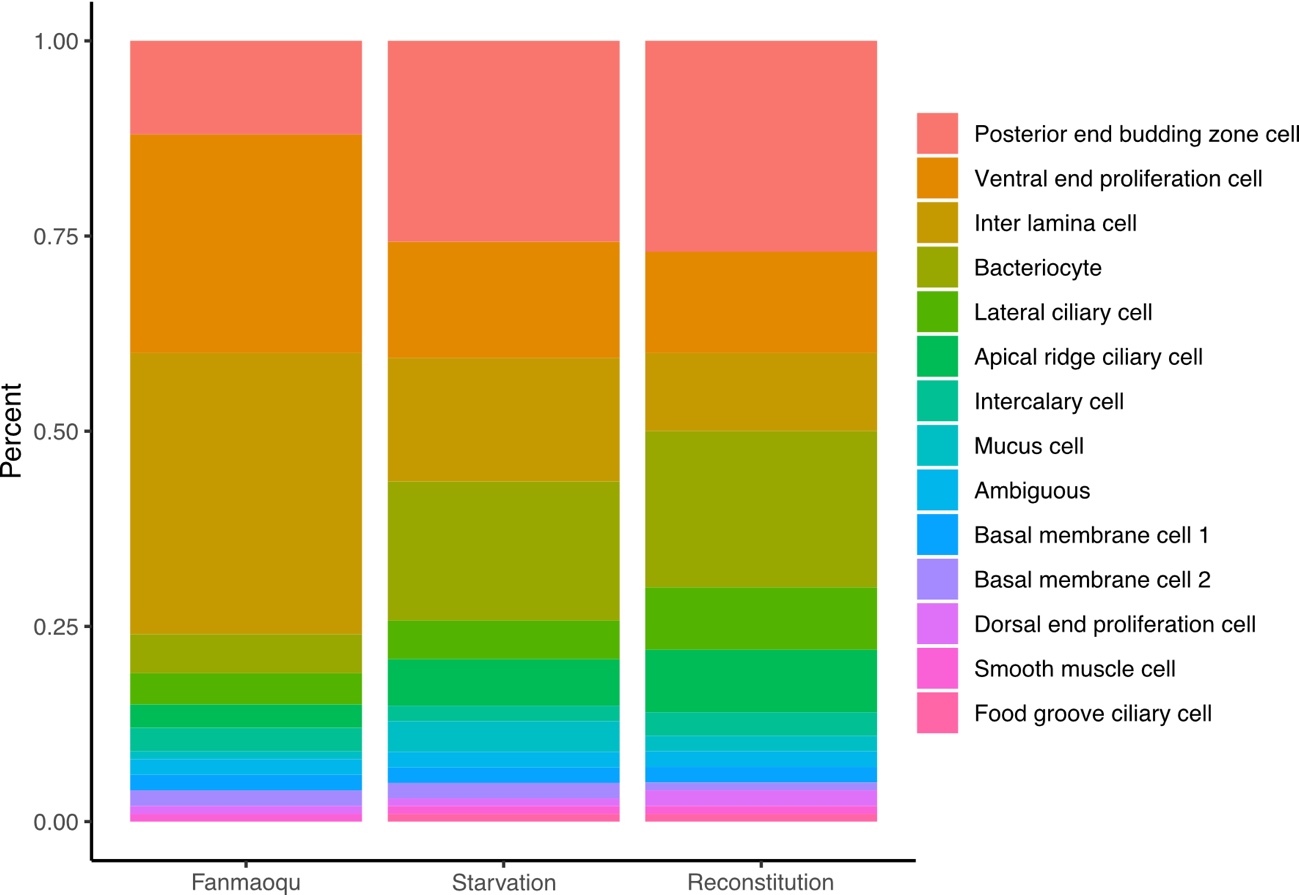

**Supplementary Figure S12:** Stacked bar plots showing the percentages of cells per cluster in each sample (Fanmao, Starvation and Reconstitution).

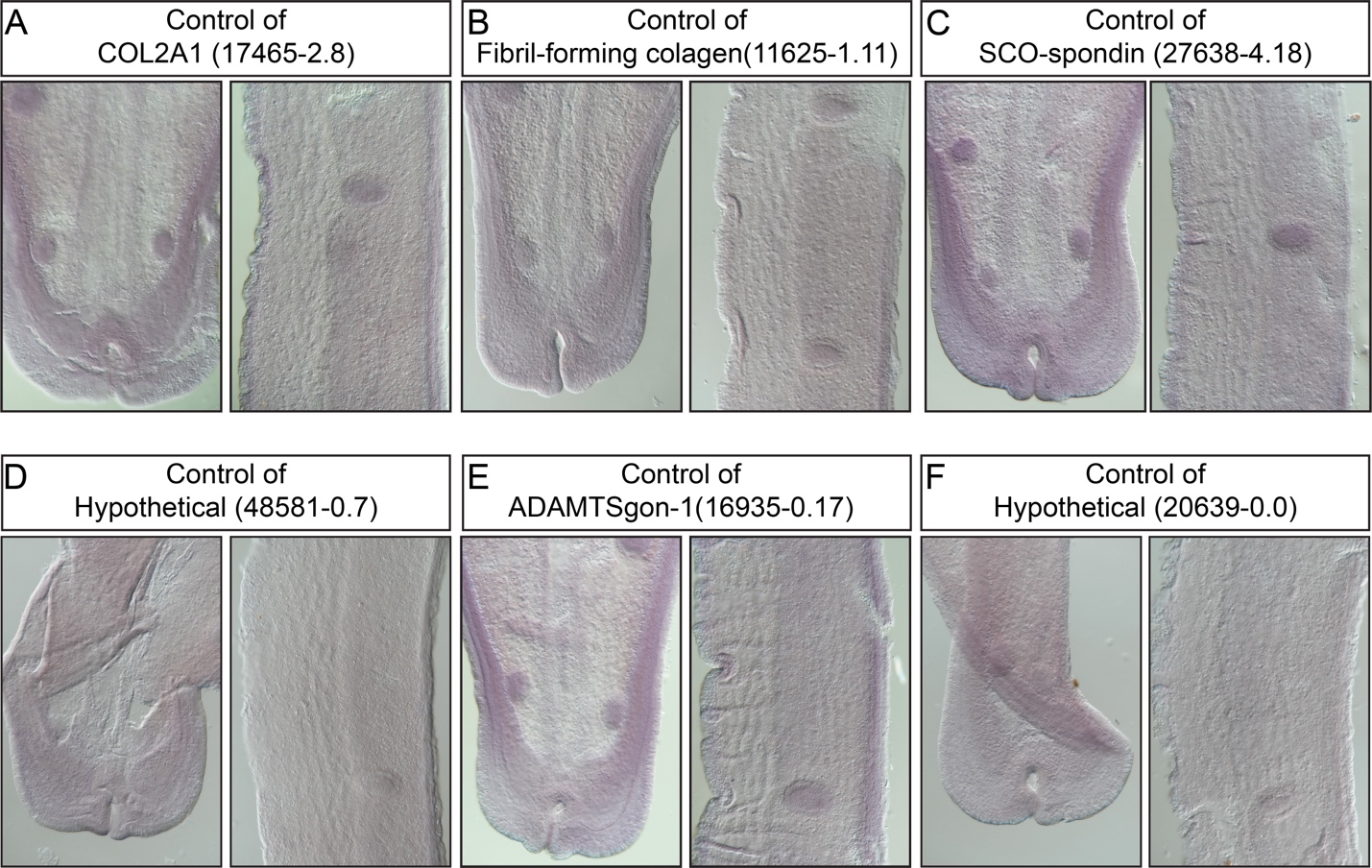

**Supplementary Figure S13:** Control hybridisation of the supportive cell markers. The left and right images of each pannel are the tip and mid part of a gill slice, respectively.

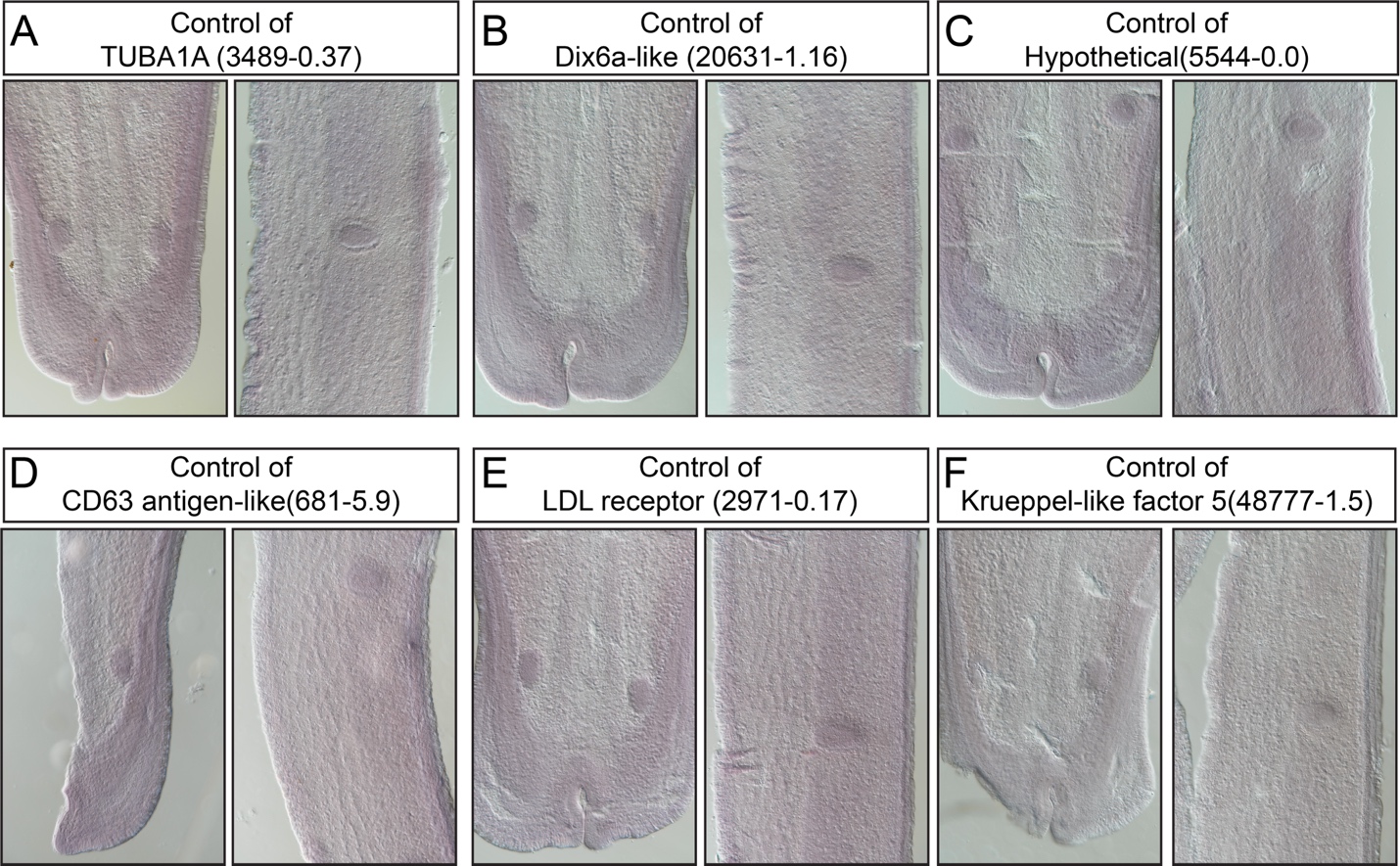

**Supplementary Figure S14:** Control hybridisation of the ciliary cell markers. The left and right images of each pannel are the tip and mid part of a gill slice, respectively.

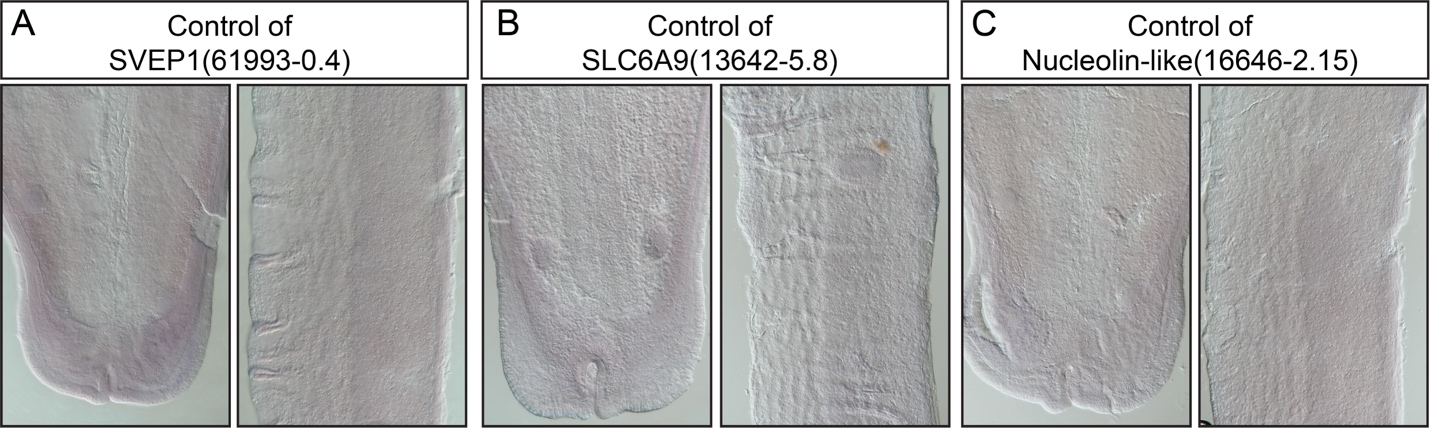

**Supplementary Figure S15:** Control hybridisation of the proliferation cell markers. The left and right images of each pannel are the tip and mid part of a gill slice, respectively.

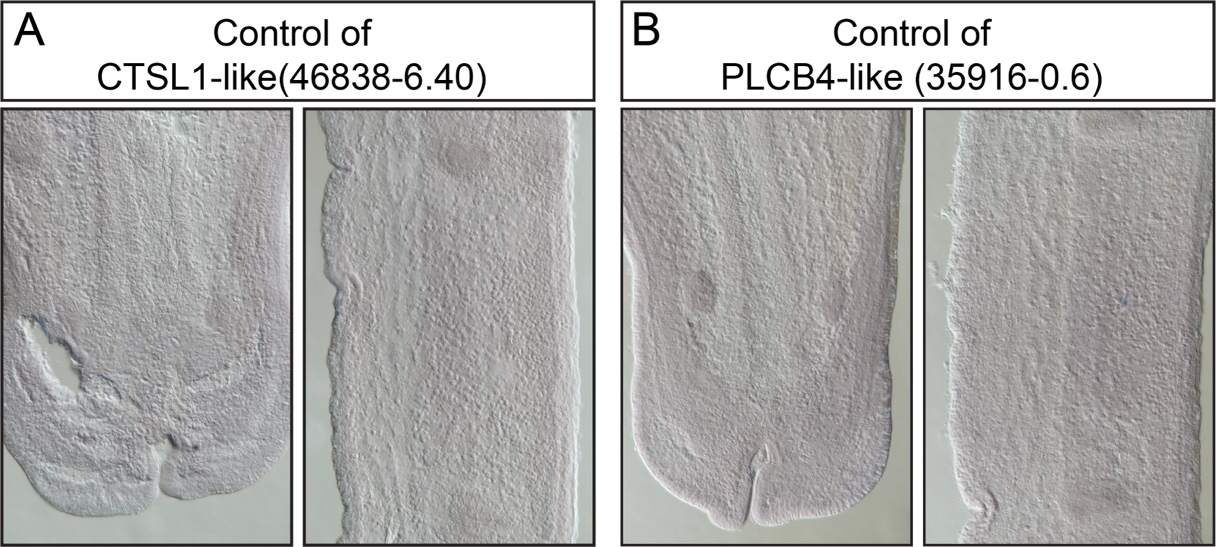

**Supplementary Figure S16**: Control hybridisation of the bacteriocyte markers. The left and right images of each pannel are the tip and mid part of a gill slice, respectively.

**Supplementary Tabel S1.** Summary statistics of single-cell data information, including numbers and percentages of cells and numbers of available genes per cluster in each sample individually and in all three pieces combined.

**Supplementary Table S2.** Identified cell markers of the supportive cells, including the inter lamina cell, the basal membrane cell 1, the basal membrane cell 2, and the mucus cell.

**Supplementary Table S3.** Identified cell markers of the ciliary cells, including the Apical ridge ciliary cell, the Food grove ciliary cell, the Lateral ciliary cell, the smooth muscle cell, and the Intercalary cell.

**Supplementary Table S4.** Identified cell markers of the proliferation cells, including the Posterior end budding zone, the Dorsal end proliferation cell, and the Ventral end proliferation cell.

**Supplementary Table S5.** Identified cell markers of the Bacteriocyte.

**Supplementary Table S6.** Cell cluster specific Differentially expressed genes among the three deep-sea *in situ* treatments, the Inter lamina cell.

**Supplementary Table S7.** Cell cluster specific Differentially expressed genes among the three deep-sea *in situ* treatments, the Basal membrane cell 1.

**Supplementary Table S8.** Cell cluster specific Differentially expressed genes among the three deep-sea *in situ* treatments, the Basal membrane cell 2.

**Supplementary Table S9.** Cell cluster specific Differentially expressed genes among the three deep-sea *in situ* treatments, the Mucus cell.

**Supplementary Table S10.** Cell cluster specific Differentially expressed genes among the three deep-sea *in situ* treatments, the Apical ridge ciliary cell.

**Supplementary Table S11.** Cell cluster specific Differentially expressed genes among the three deep-sea *in situ* treatments, the Food grove ciliary cell.

**Supplementary Table S12.** Cell cluster specific Differentially expressed genes among the three deep-sea *in situ* treatments, the Lateral ciliary cell.

**Supplementary Table S13.** Cell cluster specific Differentially expressed genes among the three deep-sea *in situ* treatments, the smooth muscle cell.

**Supplementary Table S14.** Cell cluster specific Differentially expressed genes among the three deep-sea *in situ* treatments, the Intercalary cell.

**Supplementary Table S15.** Cell cluster specific Differentially expressed genes among the three deep-sea *in situ* treatments, the Posterior end budding zone.

**Supplementary Table S16**. Cell cluster specific Differentially expressed genes among the three deep-sea *in situ* treatments, the Dorsal end proliferation cell.

**Supplementary Table S17.** Cell cluster specific Differentially expressed genes among the three deep-sea *in situ* treatments, the Ventral end proliferation cell.

**Supplementary Table S18.** Cell cluster specific Differentially expressed genes among the three deep-sea *in situ* treatments, the Bacteriocytes.

**Supplementary Table S19.** Euclid Distances between Cell Types in three different treatments.The distances were computed based on the centroid coordinates of each cell type in each condition, and were shown in Fig 6B

**Supplementary Table S20.** Quality control for each sample sequenced using the BD Rhapsody platform.
