## Supplementary_data S3 for "Deciphering deep-sea chemosynthetic symbiosis by single-nucleus RNA-sequencing"

>Bpl_scaf_17465-2.8 ATCGTCAGCAACCCTCTGATTAAGTTATATCACGTTACACACTACAAAATGTTTTCGTTGCAAAATCTGAGTCGTCTGCTTCCATCTAATTCTACCTCGTTACAGGTCAAAGGCTTATTGCTTGCATGTTTCTGATTGGTTGTTTGTTGAGCATTAGTACAATTAAAGCTAAACATGATAAAACAGTAGCGTAACGCGGCATACAGGCTGCTCAACTGCTACCATTTTCTAACAGTGGCCTAGGAATCCTGTGTTCCAATTTTCCATTGGAGCACGGCTCTTCCTAGTTGGTTTGGAGCCGGGATTCCTGGCTCGGACTTTTCTATAGATCTCAACGTTTCCTAATTCCAACCTGAATGAGGTAGATCAGAGACTATAGACAGTAAAATGAAGTTCGGATCAGTTCGGTGGAAGAACTGTCTAGCACCTTTGCTATTATTATTTGTGCTAGTTAACTCTCAGGAAGGAAAAAAATGTGTATATGATAAGCAGGAATATTTTGATGGAGACGACTGGAAACCAGATGCCCGGAAATGGTGTGTGTGTAAAGACGGGGAAGCTATGTGTACAGATGTTGCAGAAGGAGACGAATTTTCCGGTTTAACAAATAGTGGTGGCAATAATGTCAACGTAGCGCCACCGCAAGCATTGGAAACAGAAGCTGAGGGTTCTCCTGGAGGCAGAGGAAGAGAAGGAACAACGGGAGTACAGGGACCTATTGGAGAAATGGGACCAGAAGGAGAACATGGTACACCAGGATCTCCCGGACCCCCTGGGATTCCACCAATGTCAGCAGATCAAGCTTACAATCAGTATTTCCAACAGACATACGGACAGTCCTTCAAAGCTGGAGGGGCTGCCATGGGACCACGATTTCTTCAAGCCAACGTTGGACCGAGTGGCCCCAGAGGATCCCCAGGTCCCCCTGGACCACCGGGACCACAAGGAATGCAGGGTATAAGAGGAGGGCCTGGAGATTCTGGTCAGCCAGGATCACCAGGACAACGAGGAGCCCCTGGACCATCTGGGCCTGCAGGCCCTGAGGGAGACTCTGGAAGAAATGGAGAAACCGGACCCCGTGGTTTACCAGGACCAAAGGGCCCATCAGGACCTGCTGGAATGCCTGGTATGCCAGGAATGAAAGGACACAGGGGTTTCAGAGGCCCACACGGTCCAGGTGGTGAACAAGGAAGACCAGGTGACAAGGGAAGTTCTGGCCCTCCTGGAGCCCCAGGAGCTGGTGGCCCACAAGGACCACGAGGAGGTCAAGGTGAAAGAGGTGGGGACGGAAACTCAGGATCAATGGGATTACCAGGAGTTGATGGTTTAGCTGGAGCTCCAGGGGAACCGGGACCAGTTGGTAGACCAGGGCCACCCGGTATTCCTGGGCATCCAGGCTCTAAGGGTGATGCAGGTGCAGGAGGACCAAAAGGAACCCAAGGACTACAGGGAGCACGAGGAGACTCTGGAGTATCTGGGCCACCAGGTCAGGAAGGATTATCTGGGACTGATGGTCTTTCTGGACTTAACGGAGAGAAGGGAGCTTCAGGTGACCCTGGACCAGCCGGACCACCCGGATATCCTGGACCAAGAGGCCCACCTGGTATCAATGGCAGCCCAGGTAATTCAGGAGCCAAGGGAGCACCAGGTCAACCAGGAGACTATGGTTTCAAAGGTGAACGTGGACCAAAGGGTATCAGAGGATCATCTGGAAACCGTGGACAGACAGGTCCACCTGGTCTAGAGGGACAAAGAGGACAAAGAGGCTCTGCAGGAGAGGCCGGACCATTAGGACCACCAGGTGACAGGGGATCTACTGGACAACGAGGATACCCTGGAGCTGTTGGAGAGCCTGGATCTGCAGGAGAAGAGGGATCAACCGGACCAAGAGGGAGACGGGGAGAACCTGGACCAACGGGTGAACCAGGAAGACAAGGAACACCAGGACCGAGAGGAGCTAGAGGAAGTGGTGGACCAGCTGGTTTAGATGGTATGGCTGGACGACCTGGACCTGCAGGAGTAACTGGTAACGATGGAAGACCTGGAGAAATGGGAGCTCCTGGTATTGCAGGACCTGCTGGAGTTCAAGGTGCACAAGGAAATCCAGGATCAAGGGGACCACCAGGTAAAGATGGAAACCCAGGAGCCCAAGGACCACGTGGACCATCTGGAGATTCTGGACCTGTGGGAGAACGTGGAAGAACTGGACCACCAGGTGGAACTGGAGAAGCAGGAGCAAGAGGACCTGAAGGAAATGCTGGGGCACCAGGTTATATCGGTTCTCCCGGTGTACCTGGGGGTCAAGGAGAAGAAGGAAAACCTGGAGAGCCTGGACCTCCCGGGGAACCTGGAAAAACTGGACGTCAAGGATCTAGGGGTGAGCGTGGTGTACCAGGAGTCAGTGGAGAGCCTGGGCCAGCTGGACAACCTGGAGCACAGGGACCTGACGGTGGAGTTGGCAGAGATGGTGAACGGGGATCACCTGGAGCTTATGGAGAAAAGGGAGAACAAGGACCTGAGGGACCATTGGGTCCACAAGGAATGAGAGGACCACGGGGAGAAAGAGGAGCCAAGGGAGAACTAGGAGAGTCTGGACTACTCGGAGAAGATGGAAATGAGGGTCGTTCTGGAGAACCAGGTCCCATAGGAAACCCAGGACCACCTGGACCTATGGGCGTGGCTGCTGCTAAGGGTGAACGTGGAGATTCTGGACCAACAGGAGAGAATGGAGCACGTGGCGCACCAGGAGATAGAGGTCAGCAAGGTGAACAAGGAGATCAAGGTCTACTAGGAAATCCTGGTATGGAAGGTCCAACCGGAGCTAAGGGATCTAGAGGTTTCCCAGGACCAAAAGGAGCACAGGGCGAAGCAGGCATTTCTGGAAACAACGGACCACCAGGACCTAACGGAAGAGATGGTATCAATGGAAGGAAAGGAACTCGAGGTGATAGAGGATCCCAAGGGATACCAGGACAACCAGGAGGAACAGGATCTGTTGGTCCCCATGGTAATGCTGGACCATCCGGAGAAGATGGACCTCCAGGACCCCCCGGCCCTGAGGGTATCAAGGGATCTCGTGGTGAGAATGGTCACATGGGACGATCAGGAGAGTCTGGAGCACCTGGTCTGTCAGGAGAATCTGGACTTAAAGGTGCACGTGGAGAAGATGGTGAACAGGGACAAGTAGGACCATTGGGACCCCCGGGAGCACCAGGAGAAATGGGACTACCAGGAGATGAAGGAATCAGAGGAGAGAGGGGACCTGCTGGACCAACAGGACGAACAGGATTACCAGGAGACCCAGGTAGAGAAGGACATGATGGTATGCCAGGAAAAATGGGACCACAGGGACCACAAGGAAATTCTGGACCTCAAGGAGATGTTGGACACTCAGGAACCCCAGGACCTGATGGAGCCCCTGGTTTACAAGGATCCCAGGGAGAAAAGGGACCTGATGGTGATATTGGACCACCTGGAGCTTTGGGACTGATGGGCTTTGGTGGAGCACCCGGAGCACCAGGACCTGGCGGACCTGCAGGACAAAGAGGAGAAAGGGGAGAATCAGGACCAAATGGTGTAGCTGGTCAGCCTGGAGGACGAGGACCACCTGGACCTTCTGGACCACAAGGAGATACTGGAGAAAGAGGAAAGGGTGGAGTAACAGGTGACAAAGGAGACCCTGGTCTACAAGGAATGAATGGATTACCTGGACCTGAGGGACCAGTTGGAGATCAAGGCAGTGATGGACCCCAGGGACCTCCAGGACAGAGGGGACCTGACGGAAGACGTGGTGATCCTGGAACTGACGGAATGCCAGGAAGTTCTGGGCCACCTGGACCATCTGGTAGACGAGGACCTCAAGGAGAAGATGGACGCAGGGGTTCAATGGGAGAAGCTGGAAATCCAGGTCCACCTGGTCAAGCAGGAAGATCTGTTTATGGTGGAGCTATGAGGCAATGGTTTGGTGGTAACAATGGAGGAGGAAAAGGTTACCAAGGTGATGAGCCTCTGTCAGCTGATGAGGTTGATTCTGATGTGTTCAAAGCTCTGGAAGAAGTTACCCTACAGATAGAGAAGATCAGAAACCCAACTGGTGAACGTGATTCTCCTGCCCGTACCTGTGAAGATCTCAAACTCCATAATCCTGACATCAAAGATGGTGAATACTGGATCAATCCCAACCTTGGTCCTATTTATGATGCCATTCAAGTCAAATGTGACTTCAGAAATAAGAGAACCTTCACATGTGTTAAACCAGAAGTTAAAGTGCTGGAAAATTTGGCAGTGGTTCAGACAACTGACCATACATGGTTATCTGATGTCTTAGGATACAAGTTTAACTATCATCCAAGTACTTTCATCAGACCACAAATCAAATTCCTACAATATCTTCACCAGAAGGTCAACCAGAAAATAACTTACAATTGTAAAAATTCTGTCGCCATTGATAGCGATGATGAGAAATCAATCAAACTTGCTGGATTTGATGACAGTACATTATCAACCAAAGGCAAGAAGAGCATCAGATACAAAATTTCAGAAGATTCATGCAAGGTTAAAAATGGCCAGTGGGGTAAAACGGTCCTGGAGATTAACACCAAGAGGACACAATCCTTACCCATCATGGATATAGCTGTTTATGATGTAGGAGGCTCCGACCAAGACTTTAATATAGAACTAGGAGACGTCTGCTTCTTCAACTAAATTAATCATAAACTTTACATTTCATTTTTACCATTTTTTGTTTATTATAATCATTTTTTTGTTGATGTTATGATACTGTTCAAGACTTTATTTTAAATTAGAATCTATTTTTAGCACATCGGGACGGAAGGTTTGTTGATTTTTTTCAAATAACAATGATATATATATGTGCAATAATTTTTGTATTTTATGATTTGTTATTTGAATTTTCATCCAATAGGAAGGGACCATTCCTTTTACTTCATAACATCAGGGAGAATGAATTTTGTATGTAGATAGAAATATTTTCTATCAAATGAAAATTGCCATGTATGTTTTTAATTGTGTTTTTTTTTTTATCTTTTGATGAAATTTTTTGCCGTTGGACAAATTTTACATCTGTGGTCTGCCATGATACCACATTCTAACGGCAAAAAATTTCGGCCATATTTTATCTGTGATATCATTTGCCAGACACTGCCATAATTTCAAGAATGTTTACATTTTTTTTATATCATTTATGTGTAAAACACACTGTTTTATTGTATCAAAAGTCATTTTATGTGATGTCTGCTCTTTTTTTTTAACTGTAGTATTTCATAGGTCTGGAAGATTTATCTACATGATCAGCTTTCAATGAAATAAAAAAACATTAAAAGTTC

>Bpl_scaf_11625-1.11

ATGATACAAGTTATTTTCCAATATATAATCCATAGCGGTGTTTATAAGAATTCTGGTAAGGAGAAAGGACAAAGTCTGGCTGTCTGGGGATCAGGTAAGGAGAAAGGACAAAGACTGGCTGTCTGGGGATCAGGTAAGGAGAAAGGACAAAGACTGGCTGTCTGGGGATCAGGTAAGGAGAAAGGACAAAGACTGGCTGTCTGGGGATCAGGTAAGGAGAAAGGAGAAAGGACAAAGACTGGCTGTCTGGGGATCAGACTGGCTGTCTGGGGATCAGGTAAGGAGAAAGGACAAAGACTGGCTGTCTGGGGATCAGGTAAGGAGAAAGGACAAAGACTGGCTGTCTGGGGATCAGGTAAGGAGAAAGGACAAAGACTGGCTGTCTGGGGATCAGGTAAGGAGAAAGAACAAAGACTGGCTGTCTATGGATCAGGTAAGGAGAAAGGACAACGACTGGCTGTCTGGGGATCAGGTAAGGAGAAAGGACAAAGATTGGCTGTCTTGGGATCAGGTAAGGAGAAAGGACAAAGACTGGCTGTCTGGGGATCAGGTAAGGAGAAAGGACAAAGACTGGCTGTCTGGGGATCAGATGACTCAAAATGTAATTTTATGAGTAAACAATATGATATCGGAGAAAGTTTTCAACCCGATTCTCAAAATTTACCATGCCGTGTATGTAAGTGTATATCTGCCCCTAGAGATATATCGTGTGTAACCCAATCCTGTGATCCTTTGACATGTTATGAAGGCGAGAAAGAGGTCAGCCTACCAGGAGAATGCTGTCCTGTCTGTCAAATAACCATCAAACATCCAAAGATGGCAATCTCAGGAGGAAAGGTTCTTATGAACAATGATGAGAAAGCTGGAGGTGGACCTGCTTATCCTCAGGGAGTAGCATATTCTTCAGTTGGCTCACCTGGACCCCGAGGACCAACAGGACCAACAGGACCTGCCGGACCTGCTGGCTACCAAGGATCAAGAGGGGAACCAGGAGAACCAGGACAACCAGGAAACTCTGGAGAAAGAGGATTTCCTGGAGCTGCAGGCCCTTCTGGATCACCTGGAGAAGAAGGTTTACCGGGAGAACAAGGACCTACTGGACCAGTCGGATCAACTGGAAATGCTGGACAACCAGGAATGCCAGGCATGCCAGGACCTAAAGGACACAGAGGATTACTGGGAATGTCAGGAAAGACCGGAGAAGAAGGAAGACCTGGAGAAAAGGGACCAGCAGGACCAAATGGAGTACCAGGAACACCTGGACCCCTGGGACCACGAGGACAACCAGGTGAGAGAGGACGAGATGGAGCACCTGGATCTAATGGACTTAGAGGACAGGATGGAAAGACAGGAGAGAACGGACCACCAGGACAGATAGGATCCCCAGGGTCACCTGGATTCCCTGGAAGTGGTGGACCAAAGGGAGATTCTGGACAATCTGGTCAGAGAGGAGAACAAGGTTTGCAAGGTCCTCCAGGAGTCAGTGGTCTCCCTGGACAACTAGGAGAGGCAGGACGACCTGGTTCCCCTGGCAGAGATGGAGAACCAGGAGGTAAAGGTGATATGGGACAGCCTGGAGCTCCTGGAACATCAGGATTCCCTGGCCCACAAGGACCCCCAGGACAGTCAGGAGAGCCTGGAACCCCCGGGGCTGCTGGAGAACAAGGTCTATCTGGACAAGATGGGCGTGGTGGAGATCCTGGAGAGAGAGGATACCAAGGTGCCCCAGGAGAAGCCGGTCTTCCAGGTTTAGCTGGAGCAGAAGGAAAAAGGGGTGCCACAGGAATCCCAGGACCACCAGGAGCTAATGGAGTTGCAGGAGAAAGAGGATCACCAGGTGCATCAGGAGCCCCAGGGCCAGGAGGACCACCAGGAGCAAGGGGACGAGATGGTGAACGGGGACCAGAGGGAGAAAAGGGAAATTCAGGGGAACCAGGGAATGCAGGAACCCCAGGAATATCTGGACCTCCTGGACCAAGGGGACCAATGGGACAGACTGGAAATGAAGGAAAGCCAGGAGCCCAAGGACCAGCTGGTCTAGGAGGAAGTGATGGTCGACCAGGAGAGCAAGGACAACAGGGAGCACCCGGCGTACCAGGTCTAATAGGACAATCAGGAATATCTGGTGAAGAGGGAAGACCTGGACGTGATGGTGAAGGTGGACCACCAGGACAGCAGGGACCAAGAGGACCGAGGGGAGAATTAGGAGAAGGAGGACCGGTAGGACCACCTGGTAATTCTGGACCACAAGGAGAGAGAGGAGCCCCAGGGCCACAAGGAGAAGCTGGCATTGGTGGTTTAGCTGGAATAGCAGGTGCCCCTGGAGAGCCCGGTAGAGCAGGAGAATCTGGAATCCCTGGACCATCTGGTGAGCCAGGAACAGCAGGAGAGAGAGGTGTACAAGGATTTTCTGGAGAGCCAGGACCTGAGGGAAGGCGTGGACCAACTGGAGAGAGAGGTATGCCAGGACCCGCAGGAGAATCTGGTGGAGAAGGACCACCAGGATCTCAAGGAGAGCGTGGAAGCCCAGGACCTATCGGATTAGTGGGATTACAAGGAGACAGAGGACCCCTTGGAGCCCCAGGATCAAGAGGAGGCAGAGGATACCCAGGTGAAAATGGAAAAGACGGAGAACCCGGTAGACCAGGGGAGCAAGGACAACCAGGAACCCCAGGACAACCAGGACCTCAAAATTTAACACCTCCACCAAAGGGAGATTCTGGAGTGCCTGGTGAAATTGGAGCAGAAGGAGTAAAGGGTGATATCGGACCACGAGGATATCCTGGTAACCCAGGACCAACAGGGCAACAAGGACCTCCGGGACCATCCGGACCAGCAGGAGAAATTGGACCTGAAGGAAGAAATGGACCAGCAGGAGAATCTGGACCTCGTGGTTATCCAGGAGAACCAGGAGCTGTTGGTGAACAGGGACGTGACGGACTTGATGGAGATAACGGTAATTCTGGTGAAGCTGGTCAACAAGGCCCAGCTGGTGCCCCAGGTAGTCCTGGTAGACCCGGGCCACCAGGACCAGCTGGTCCTCAAGGAAACACCGGATTCTTAGGAGCTGCCGGAAAAGCAGGTGCAAGAGGAGATCGTGGTGAGACTGGTCCAAATGGACCTGCTGGCAGAAATGGAGAGCTTGGACAACCCGGACAAAATGGCCACATGGGAGAGAGGGGAAGTTCTGGACCCCCAGGACCACAAGGAGCTACAGGTGCCTCAGGACCTCAAGGAGAGAGAGGATCTCCAGGATACCCTGGAGCCCAAGGTGAACCAGGACCCGGAGGTGAACAAGGCCCACAGGGACCACCAGGCCCATCAGGCAATAATGGACAGGATGGACTAAATGGACCACCGGGACAGCCAGGACCTGTTGGACCAGCAGGATTCCAAGGACCACCAGGAGAGCCAGGAACTGCTGGTTTGAATGGACAAGCAGGACAAGTTGGTAATTCAGGGTCCAAGGGATCAAGAGGCCCAAGTGGTGCCCCAGGTTTACAAGGACCCCCAGGACCTCAGGGACCAAATGGACCTTTGGGAGCTGATGGATCACAAGGAGAACGTGGAGAGAGGGGAACAGCAGGAGAAGGTGGAGCACCTGGTATCCCTGGGCAACAAGGACCACCGGGAGTATCGGGACAGGCTGGGGATAAAGGAGAAGATGGACGCTCTGGAGCTAAGGGAGACAAAGGTTGGCCAGGTATGCCCGGAGGTCAAGGTTTACCAGGACCTCAGGGACCAGGAGGAGAGAAAGGTTTGAATGGACCACCCGGACCCCAAGGACCTGCTGGAAGTACTGGTTCCCGAGGAAACTCAGGAAGAGATGGAGAGCCAGGGCCGCCAGGATCTCCAGGAGGACCAGGAAGTAGAGGACCACAGGGAGATGACGGTTTGACTGGCATAGCAGGACCTGCCGGACCTGCCGGACCCCCTGGACCACCAGGCTATGCACCAGTATGGCCAGGACTGAACCAGTACCAGCAGAACAAGGGACCAGATTCTCAATATTATGGTGATGAACCAAGCAAGAAACCAATGGTATACGATGACTTGTCCAGAATCCAAGAGGCTCTCCACCGTACCAAGAAACCAAGTGGCAAGATATATAATCCAGGTGTCACCTGTAAAGATCTCCTCATCCAGAACCCAGACTTTGAAGATGATTGGTATTATATTGATCCTAATGGTGGTAGCTTCATTGATGCTGTCCAGGTTTACTGTAGAATGAAATCTAATGGAGAAACTTGTATCCCAGCTGATCCTAAAACTTTTGAGAAACAAAGATGGACCAAGGATTCCAGACCTCAATGGTTTGGACAAGAAATACTTGGAGGCAAAGAGTTTGACTACCAGATTGAAGCTATCCAACTGAAGATGCTGCAGATGCACAGCACAATTGCCAGACAGAGGATCACATATAAATGTCTGAACTCTAACCCTACCGGTTCTGTTCTCAAGTCCAATGAAGATATACGTATAGACACACGTGCTGATAGACAAGAACGTGGCCAGTCAACAAGTGTGGATGTATCTGGCAACTGTGCTGAAAGTAAAGAATGGGGTGAAATGGTCTTTGATATAAAATCACAGAGATCAGAAGTATTACCCATCACCGATATAAAACTCAAAGACGTTGGTTTACAGGCTCAAGAATTTGCTCTTATCATAGGAGAAGTCTGCTTTAACACATAAAAGACTAAATAAATTTATGTACATTTATATCAAATTGGCTTGAAATCGTGGCAGTTTTGTTTTATGGCAACTTGAAATCATGGCAATGTAAAAAAATATTGTTGGAAAATCATGGCAGAAATTTATTTGCAAATGTATACTGGTTTCCTGTCCTCTATATAATGGACTGGGAATCAGTTTACTTGTAATATTGAATAATTTCATAACAATGGGACCATCACATTAAATTTTGTAAAAAAAAAAAAACATGGTTCTTTTGAAAATCTTTCAATTGATAAATCATGGCAAAATAATTATAAATATTGAAAGATTTTAGAAACGACCATTGTGTACAAATATTGTAAAAATCATGGCAAATATTTATTTTTGTAAGACAGACCTCATGGCAACTTGATAATTAAAGAAAAGAGTTACATTACAGATATTGTGATGTTATTTGTATGAAGGAGTTCGTTCCCTTTTGATTCCGTCCAAATGTGTGTGTAAGAGAGCGAGAGAAAGAAATGTATGCAAATTTTATAAGAAAATCTTATTTTGAAATATTTTATTGTTTATTCAATTTTATATGCTCAAGTCTTTCTTAGTTCGTCAAAGACAATCATATTGAACTTATATCCTTGTAGTTTGATGGAAATTAGTTGGACTAAAAAATATGGCAACTTTTTTTATGGTTTTGAAAACATCATCATATTATGTTAAAAAATATTATTAAACAAATCATGAATATGCATAATATTTACTCAAGGAGAACATTTATTGTGAATCAGTCATATGGAACATTTTATTAAACATCAAGCAAATATGAATAAAATCACTCATGCAGAAAAATAGATCTATTTATATCTTACAGTGTCATATCATCTATTCATTTGCTTCATCAATAATTCAGTTTATGGAATAAAATTTCAGAAAATGTATAACTTTCTTTTATCACTTTATTTATGTTGCCTAGTTACAATATTTGATACATGGGGCCATTTTGTGAACTTACCAGAGCTCAAGTGAGTTTT

>Bpl_scaf_27638-4.18CGTCTTCAGAGGACAACTAGTCCCTCCTAAAGAACAGATTGAAGGAGACATAAGAACTTCTGTATTGTTTTAAGTGTATACGGACATAGCTACAGACATAGACTAATTGGAAGTTCTTTACAACAATTACGGTTTTTTTCTACGCCCCATTTTTGGGACCAAATATGGCCAGGTCTTTACAGACCCTTTTCATTGGGATTTTATGGATTTTAACAACCGGAACGTTACAATGGACCTTGGCTGAAAGTCACATATGCCCAACGAAGGTTACAGAAAAGGAAGAACAAACCGGAAGTTGTGGTGACTTTATGACGATGACAATGCAGCGTTATTTCTTGGATAAGGAGAAATCATTCCCTGGTCAAACAGAAGAATACATCCAAAGTGAAATAACCAAATACCTGAACATCTACAACAAAAATTTGCCTCCGAAGACATGCTATTTTTATAGGAAAATAGACCGGGATGTAGATGATTGTTGTCCAGGATACACCAATGGAACAGGTCTATGTGACACTCCTGTTTGTGGAGATGGATGCAAAAATGGCGGTTCCTGCCTTAAACCTGGTGTATGCAAGTGTCAATCTGGATATACAGGATTTGCATGTGAAGATCTAGCAGAATATGCTGCAGGAGATTTGAGGTATTGTTACAAAGGAAAGACTTGTCACAATTCTGCAAAAACTTTGGGTGGTATACCTGTTTCCCAGGAACAGTGTTGTAATGCTGCCGATGCCTATAGTTGGGGTGTTGGATCTATATCTTGTATGGAGTGTATTAAAACAAGTGCATCAGGAGTTATTATGCAAGAAAACTCACTGAACTTCAGAACTTGTCTTAATTTTGGGACAAACTATTACAGAACATTTGACGGTTTACAATTTTCATTTGGAGGACGCTGTACATACACCATGGCAATGTCAGAACATTGGAATGTACAAATGCAAATCATAAATTGCAACTTGTTTGATACCTGCAGAAAGAGAATTACACTAGACATTGGAGGAGAGAAAATAGTCTTTGAAAATGGAATGGTTTGGTGTGGCGATAAAGAGTTTCCATTGGAGGATCAGCAACCAAGAGCGACTGATAATGGCGCCTGCACCATTTTCTCCAAGGGAGACTTTATCTTTTGTGAATGTTCAAGAGGAATCAGACTAAAGGTGGATGCGTTATCAACTGTTTATATAACTGTGGTAAAAGATCAAGTGGCAGCAGAATCACTACAGGGTATCTGTGGTAACTTCAATGATGATGCTACTGACGACCTTAAGACGAGAACCGGAATGAAATCTACAAACCCAGTATATGTGGCCAACTCTTGGGCAGTCGCTAATGAAGCAAATGAGTGTCCATCTGCTGGATCTCAACCTGATTATTGTAAAAGCGCTCAAGACAAGAACATGGCTGAACAAGTATGCTCTACAATGATGTCCGGAATATTTGGAGAATGTCATAAGTTGATGAGTCCATATTTCCTGTATCACCTCTGTATCAATGAAGTTTGCAACAACAAAGACAATGCAGATATGATCAAGTGTGAATTTACCAGTAGATTTGCACAGTCTTGCGCCAGCCTCGATGTGATTGTGTTCTGGAGATCACCTAATTTTTGTCCAAAGACTTGTGAAAATGGTAAAGTGTATCAAGAATGTTCATCTAAATGTCCCCGGACATGTAAAACACTTTACACAGTCATGCCAGACTCCTGCATGCAAGATTGTGTTCCTGGCTGTGAATGTCCAATTGGGCAGTTTATTCAGGACGGAAAATGTGTCGTAGCTGAAGATTGCGAGTGTCAATTTGACAAGAAATCGTACAAGACTAACGAAACAATCAAAAATGGGTGTAATTTATGTAAATGCAACATGGGTCGTTGGCAGTGTACCGAGGACAAATGCTCTGAGATGTGCGAGCTTGTTGGTATAAATCATGTGAGAACGTTGGACAACTATGAGTACTCATTCAATCCTGGCAGCCTTTGTGAATTCAAAGTTGTGTCGCCGTATCCAGCAAATGTAGCATCAACAGATCCACGTGCAGATCTGCATATTGACTTAGAAACCGATAAATGCACTAATATGAAATCTGGTTATTATTGCTTGAGCAAGGTGATAATTACATATAGAGGTACAAAGGTGACGCTATCAGGAAGCTCAGTTACTGTTAGAACAAATGGTGGAGATACGACCGACCTTTCAGCTGAAATAGACAATAAACCTTATCATACAAAACACATCTACGTCAAAGCGCCAAGTAGACAGTACAGATTGGTCAAGGGTTTTGGATTTAAGATATTATATGATCAGAAAAGGGCATTGTATGTCTACTTAGCACCTTATTTTGCTAAAAAGGTGTATGGTCTTTGTGGATACTACAATTACAGACAAGATGATGATTTAACAACAATTAGTGGTCTGCCTGAACTTAATCCATATTCTTTTGTGAAGAAAATTTGCAATGCCGAATGTAATGTCAAAGAACCAGCAAATGAAATAATCATAGAATCAGAAAATACAATGGCCAAGGCAGAATGCTTGTATTTGAACCCAGCAGCTACAGATTCTATTTTTAAGTATTGTATTGAAGCAACTTATGATAAAATACAATATTATTATGACCGATGTGTAGCTGATACTAACCTGAGATCCAAGACCACAGAAAAATCTACAGTATGTGATATGGTATCCGCCTTTGCTAGACTCTGTTCGCTAGCTGACGTACAAGTAAAATGGTACGAGTATGGAGATTTACAAACCAATTGTGCATCTGTGTCATGTACAAATATTGGCCAAGGAGGAGAAATATACCAAGAATGTGGTAGATTATGCAAGTCAACGTGTAGAGACTTTGAAATAAATGATGCAAGTTGTGAAGATGAATGTATTCCTGGCTGTCAATGTGAACCCGGGACCTACAGAGATGACTATGGATCATGTGTTAACCTAGAACAGTGTACCTGCTACGACATGTACAATCTCCAACAGAAAGTATGGCCAGCAGGCTCAAACATTACAAGACAGTGCTCTACTTGTACATGTCAGAAAGGTGCTTGGGATTGTGATACTGATTCCTGTGAGGATATTACGTGTCCAAAGAACCAAGAATATATAGATACTGAATCAATATGTCAAGCAAAAGTAACATGTGCCACATATGACTTAAAAGCAAGTTGCAGCGATTCCATCACGAGGTTCAGAGGATGTGGATGTAAAAATGGCACAGTAATGGCACCAGATGGTACCTGTGTAGTGCCTGATAGATGTCCCTGTATGTATGGAATGGATTACTATGATGAGGGTGAAGAAATTACTGTTAAGTGCAACAAAATGAAATGTCAAAGTCGCAAATTCATCAAAGTAGGTCAAGTTGACTGCCCAGGAGTATGTTGGGTATACGGTGATCCTCATTATGTCACCTTCGATGGAAAACATTACATGTTCCAAGGAGCCTGTCGTTATGTCTTGGCCAAAGCAATGAATGGCGATTTTAGCGTTGTAGTAGGAAACATGCCATGTGGATCCACTGGAGTGACTTGCACAAAGAATGCAGAAATTACCATAAAGGGTATCAAAATGCACCTTATTAGAGGCAGTGCTGTCAAAATAGGAAATACTACACTGAATGAACAATATATAAGTGAAGGACTTGAGGTAGCCACATACAGTTACTGGACATCAATCGTTGCTAAAACATTAGGAATCGAGATCATGTGGGATGGAGGCACCAGAATGAAGATAAGCTTAGATAAAAAGTGGATGAATGGAGTAGAAGGTCTTTGTGGAAATTTTGATGGAGAGTCTGAAAGTTTAGATTATAAAAAACCAGACGGAAGTCAAGGTGTCTCAGCCAACGACTTTGCCATGAGTTGGAGTGCAGACTCAACATGTGACGATAAATCAGCCAGCGGTTCAAACACAACTATACCTACAGGACCATGTACCGGAGACTTAGCATATAGAAAGGAATGGGCTCAATCGTCCTGTAGTATAATATATTCTGATGTGTTTAAAGACTGTCGCGGAGCAATAAGCTCGGCAGATGTTACCAAATTCTATGATGATTGTTTGTATGATTCTTGCAGCTGTGACAGAGGAGGAGACTGTGAATGTTTGTGTACTGCAGTTGCTGCCTTTGGGGAACAGTGTAATCAGGCTGGCGCCCCTGCCAAATGGAGAAATCCAAGATTTTGTCCAATTATGTGTCCACTTGGTTTCGAATACAAAGCTTGTGCTAAGCCATGTCCACAGACCTGTAAAAACATTGGTGATGATCCAGACCCATGGTGCAAAAGTACTTACTGTATTGAAGGTTGTTTCTGTCCAGATGGAATGGTCCAAGATGGAAATAAATGTGTTCCGGGTAATGAATGCCCATGCATGCACAACCACAAACCATATCCACCAGGAACATACATCACTAGCGACTGCATGAACTGCACTTGTATCAATGGAAAATTCGAATGTACAGGAACCAGCTGTGTATCAAAATGTAATATCACCGATGAGTTTACGTGTACAAACGGTGACTGTATTGATGAAGTATACCGTTGTGACAAGCATCCTGACTGTAGGGATGGTTCAGATGAAGTTAATTGCACATATACCTGTCAGACCAACGAAATGTCTTGTGATGATGGTAGGAAATGTGTTGCCAATGGCTATCGATGTGACGGCATGAGTGACTGCTTAGATTCAACAGATGAAATGAATTGTGTAATCAAATGTGATAAAACACAGTTTACCTGTGCTAGTGGAAAATGTATTCACATGAAATATGTCTGCGATAATTATCCTGATTGTGGTATTGATGACAACTCTGATGAAGAAAACTGTAATGCTACTGTGTGTGCATCCATGCTAGAATTTAAATGTGCAAATGGAAAATGTGAGCCAATAAATCATAGATGTGACGGGCATGATGATTGTGGTGATGGATCGGATGAGGAAGACTGCACAACTACACAACCATCAACAACTGAAATAACTACTCACGTATCAACAACAACAATTTTCTCCACGCCTACTTCAACAGGATCAACAACAACAATAGTTTCAACGCCTTCTACAACAACAACAGAATATTGTGTACATGTAGAATTAATGAGTGACAATATCAACATTCCACTAAGTAAATTAAAAACACCTAATAATCCAGGATTATCGGACGAAGATAAGGAAAAACTTCGACCAAATAATCAGGAATTTCTCGATGTATTTGGTGGCTCTTTCATAGTACAACTAGAAAATATTGATGCTGAAATAACCGATATTACCATCGAGGGATCCAACATTGCTGAAATTACAGTAAAATATGACGATGTTCTAGACAATATAGAGTCTTTAACACCACTGGGAACGTTTACTGCTGGTGACAAAATTCCATTGACACCGAAATTGTATGAATTTATCAGAATATCTATAAAAGGAGGTTCTCCTGATAAACCTATCACAATTAATGGCATACAAGTTGGTGGTTGCGCAAAAATAAGTACGACAACAACAACAGAAAGTACAACTACATCCCTGCCTACTCCTACTACAACAACGGTATCTACTCCTTGTGTGGATCAAAACATAATGGAAGACTCCATCAAGGTCCCTCTTAGTTCTATTATAACAAGTCCAGTAATCTCGTCTGAAAATCTGCAAGCCATTCTTAAACCAAGTAACCAGAGTCCAGTAACCATAGAGGGTTCTGAAATATCAATAACTGTTAATGTTGTGGCAGAATATACAGCCATTGAGGTCAACGGAGAAAACATCAATATAACATCAGCAACGTATATACCTGTTGGTAGTAATGTTATTAAACCCTTGCGTGATGAGCCATTTTCAACAAACAAACTTGAGGAATTACCTGCATCGGTTGAGGCAACATTAGTGGTAATAACAATAGCAAAGACCAATCCAGAGTTGCCAGCAACAATAACCAACATTGGTGTAAAAGCCTGTATTGAAGGAACTACAACTACATCCCTGCCAACTACTACTAGTCCTACTACAACAACGGTATCTACTCCTTGTGCGGATGACAACATAATGGTCGACTCCATCAGGGTCCCTCTTAGTTCTATTACAACTGAGCCAAGCATCAGCACATTTGAATTACAAACTATTCTTAACCCAAACAACAATATTTCCGTGCAGGTTCCAGTGAATACTGGTCTCACAATAACTGTTAATGTTGAGGCAGAATATACAGCCATTGAGGTCACTGGAGACAACATTGAGATTACTGGAGCATATTATATACCTGTTGACTCTAATGTTTTAACACCATTGGATGGTCAGCCATTTTCACCAAACCAGCTTGTGGAATTACCTGAATCGGTTGAGGCAAAATCAGTAGTAATTGTAGTAAATGTTTCCACAGGTGTGGAAGCATCAATAACCAATTTAGCTATTAAAGCCTGTATCGAAGGAGCAACAACAACTCTACCTTCAACAACTAAACTAACAACAACTGGAGAAACAACTACAGCTACTCCATCAACAACAGGACAACCAACGTCAACTCCTCCACCAACCACAAGTACAAGCCCACCATCAACACCACCCCCTTCAACCACTATCCCGCCAACTACCACAGGTATCACTCCAGAATCAACCACATCAGGAGCTATCACCACACCATGTGCAAAAGAACCTCTGGATGCAAATAATGGTAGGGATGATTTCGTTTTATCGGCATCATCAGGACCAGATATATACAACGCCTTACTGGATATGAGTTCATCAGTTACTAAATCTTGGAAACCCAATTCAGGTGATTCAACCCCATCAATCGTATTTACAATCACCAGTTCGGATATTCCAAGATTAATGTTTGTGTCATTGGTAGTGACTAACACTGAAACTGTGGGATTACAAGTAGCTGACACCGTAAAGTCAAAGCAAGTAAGCGGAAATGGCACAGAAAGGGTTGACTTCACTTTCCCAGAGGGCTTACTCTTAAACGAAAGAAATTTTACAATACTATTGATGCCACGTAATGGTATGGTACCTAGTGTCAGTGGAGTGCAGGCTGAAGTTTGTTATGAACCTGAAGGAACCACTACCACAGTTCCAGCAACCACAACTATAATTGTCACAACTACACCAGTCACTGTAACTACTCCTAGAACAACGACAGCATTAGCATGTGATCGACCAGAGTGGAGATGTAATGGTACAAAGGAGTGTGAAGATGGCAGAGACGAAGAAGGCTGTATTTCAGCAACTACTACGCCTCCAGCAACTACGCCAAAGCCTTGTGAACCTCCTGAGTTTAAATGTGACGATGGAACCTGCAATGTGGAATCTCAAAAATGCGATGGGATTTGTGATTGCCTCCCTTCTTGTGAAGATGAAAAAGATTGTCCTGAAAAAACCACAACTCCACCAATAGAAACAACGACAGTTGTAACAAGTGCGAGTCCTGGAATTAGTTCAACAACTCCTGAGATTTGTGAATATAGATGTGGTGCAACATTAATTTGTATTGAACGAGAACAACTGTGTGATGGACACAATGATTGTGGTGATTATGAGGATGAGAATGCGTGTTCTACGGCTGTCACAACTGTTGTATCATCAGCAAAGGCAACGACGTCTGAATCAATGGCAAGTCATAAGACAATGACACCGCCAGTAACAACGTCCATAGGTATTACCCCCGTGAAGCCTAACGGCTCCACCACACCTGAAGTTACAATTTTGACAATGTTTTACTATCGACGAACTGGAATTTAGGGTCCAACAAATGTGATTCTAAATGGTATGGAGACAAGAAAATCTATTTGTATTGTCTGATTAAAATATGAATTTGATTGAATTTTCCACTTATTCTATATGAAATACCAAAGAGAATCTATAATGAATTTTGACATACATATAATAATATAAAGCAC

>Bpl_scaf_48581-0.7

TGAAATTCCAACACATAATAAATAACATCGTGTCTCTTACATAGTTCTGTAATATGAGATGGTGCATGTATCTGTACAATTGAAAGCGTTTTATCGACTTTTTAGTATTCTGTCAATAATACTTGTAACAGCTAGTTAGTAGTCTCCAGACACTGTCGAAACATGCAGCTGCACTTCGATTTGCTGGTGATTTGTTCCGTAATTATTTTGAGTCTTCTTGATTCTGGCGATGCGCACAGGAGGTGTTCAAGAAGATACACGTGCTATAAACCACAACGATGGAAAACAACCCATTATAGAGCATGTGGTTGGTTGTGGAGACGCAGATGTACACGACATGTGACATACTTCAACAAAATGATCAAGTCAACCTGTCATAGAACCTGCAAGGTAAACGGTGGATGGAGTTCTTGGTCCTCATGGAGAAAGTATGGATCTTGTTCAGCATCGTGTAGGGGATTATCCAGCTATCCACAACAAAAATATTACAAAAGACGATCTTGTAATAATCCTACTCCACGATATGGAGGAAGACGTTGTCGAGGATCTTCGTGGCGGTACAAATACGTCAAATGTAACACACATTACTGTAGAAATCCAAGAGGTGGATGGAGTTCTTGGTCACCATGGCGCAAGTCTGGATCTTGTTCAGCATCGTGTAGGAAATATGGATCATCTAGCTATCCACAACAAAAATATTACAAAAGACGATTTTGTAAAAATCCTCATCTACGACATGGAGGAAGACGACATAGAGGAAGACGACACGGAGGAAGACGTTGTTCAGGATCTTCTGTTCAGTACAAATACGTCAAATGTAACACACATTACTGTTGAAATAAAAAGACTCACAGATGATTTCTGAAGATTAAAATTCATGAAGACTCACAGATGATTTCTGAAGATTAAAATTCATGCATTATCAC

>Bpl_scaf_16935-0.17

CAATGGCAGCTTTCTTCTTAAATACAAACGGATCGCAGTGGCTTATCAGACCCCACCACAACCATGGGACAGATTACCATACTATGTATATTTTTCGTTTCATTTGTTGCTCTTGTTGAAGGGGTTTGCAACGGTTACTACTACAAAACATGTCACAAGTATACATATCAGAAAGAGAGATATAGAATAAAATGTGGATTGTGGCGGTTGAGGCGGTGCGCTAGATACAGACAAAAGAAAATATCCGTGTATTACCGATGTGCAGAGAGATGTCCAAAAGTGCATGGAGGTTATGGTAGTTGGGGTAGGTGGAGTTCGTGGGGCAGTTGCAATAAATATTGTGAAGGTGGCAGTCAAACTAGATACAGACTCAGGTCATGTAATAGTCCGACACCACAAAATGGAGGACGCAATTGTTATGGATCTAGCAGTCAGTCGGGAAGTAGATCATGCAATACACAGAAATGTAAAAGAAACGGTGGTTGGGGATCTTGGGGCACATGGTCAAGCTATAGACAATGTACTAAGACATGTGGAAGAGGAGTGCAGTCAAGAACTAGAAGCAGGTCATGTAACAGACCAGTTCCCAGATACGGTGGTAGTAATTGTCCTGGTTCATCATCATCGTCATCTCAAAGGTTATGTAATACCAGGGCTTGTCCAGTGCATGGAGGTTATGGTAGTTGGGGTAGGTGGAGTTCGTGGGGCAGTTGCAATAAATATTGTCGAGGTGGCAGTCAAACTAGATACAGACACAGGTCATGTAATAGTCCGACACCACAAAATGGAGGACGCAATTGTTATGGATTTAGCCGTCAGTCGGGAAGTAGATCATGCAATACACAGAAATGTAAAAAAACATAATGTGGATAGATATAAAGCGTCGGAAATAAAGAGACTACTTGATTATTCCTTAGAAATAAATCCATTTTAAAGCATG

>Bpl_scaf_20639-0.0

GGTAAAGACGAACCGAACATCGTTTTTATGCCGAAATCGTAACGAAGTTATCGTCATCCTAACATATCTATCTATATTTTTGTTCTAAAAAACACGATGTGTATTTATGTTCCTGTATTGTCCTAATTATATGACGCCACGATATAGCTGAAATATTGCTAAAGTTGGCTTTAAACAATCAATCAATCAATCAATCAATCAATCAATCAATCAATCAATCAATCAGTCAACCAATTATAGAGGATTAAGAATTTCAATTGAACTGAGTATTTATAACCTTTTATATTAACAGATTATGACCGTATTTGTATATTTGTTCATTTTGACATGTTGGTGCTATCAACCCGGTTCCACTGCAGCAATAAATGAAGTACCTATTATCATATCAGCAAGTTTTGATGACGTCAAGATGGAGCAATACATTGACAGCAAGATCTCCAAAGGATTAAAGGGCATAAAACGAGGATATGTCGGTTGTTTTTATGATGATGGACGGAGACTTCTTAAGTACAAAATACGAACCTTCGACGTCAACTCCATACACAGGTGTCGTGAACATTGCAAGGGATATAAGTATCTAGGATTACAGGTA

>Bpl_scaf_3489-0.37

ATACCTATTTTTCTCACACGACACCTCAAGTCTCAGTGAGCTTAGGCTTTTTGTTTTTATTTTAAAAACTCAACAACCAAACTTTCAACATGAGGGAATGTATCTCCATCCACGTCGGACAGGCCGGAGTCCAGATCGGTAATGCCTGTTGGGAATTGTACTGTTTGGAGCACGGTATCCAACCTGACGGTCAGATGCCATCAGACAAGACCATTGGAGGCGGTGATGACTCTTTCAACACCTTCTTCAGTGAGACCGGAGCTGGCAAGCACGTACCAAGAGCTGTCTTTGTTGATCTGGAACCAACAGTAGTCGATGAGGTGAGAACCGGTACATACAGACAATTGTTCCACCCAGAACAACTCATCACCGGAAAGGAAGATGCCGCTAACAACTACGCCAGAGGTCACTACACCATTGGTAAGGAAATCGTTGACTTGGTATTGGACAGAATCCGTAAATTGGCTGACCAATGTACCGGTCTCCAAGGTTTCCTCATCTTCCACAGCTTCGGTGGTGGTACCGGATCTGGATTCACCTCACTCCTCATGGAACGTCTCAGCGTTGACTACGGAAAGAAATCAAAACTGGAATTCGCCATCTACCCAGCCCCACAGGTCTCAACCGCCGTTGTTGAGCCATACAACTCCATCTTGACCACCCACACCACCCTTGAGCACTCCGACTGTGCTTTCATGGTAGACAATGAAGCCATCTACGACATCTGCAGACGTAACTTGGACATTGAGAGACCAACCTACACCAACTTGAACAGATTGATCGGTCAAATCGTCAGCTCAATCACTGCCTCACTCAGATTCGATGGTGCCCTCAACGTCGACTTGACCGAGTTCCAGACCAACTTGGTACCATACCCACGTATTCACTTCCCTCTGGCTACATATGCCCCAGTCATCTCAGCAGAAAAAGCCTACCACGAACAATTGTCTGTTGCTGAGATCACCAACGCCTGCTTTGAACCAGCCAACCAGATGGTCAAATGTGACCCACGCCACGGAAAATACATGGCCTGCTGCATGTTGTACAGAGGAGATGTTGTCCCCAAGGATGTCAACGCAGCCATTGCCACCATCAAGACAAAGAGAACCATCCAATTCGTCGACTGGTGTCCAACTGGATTCAAGGTCGGAATCAACTACCAACCACCAACTGTTGTACCAGGAGGTGATTTGGCTAAAGTACAACGTGCCGTCTGCATGTTGAGCAACACCACCGCCATTGCTGAAGCCTGGGCTCGTCTTGACCACAAATTCGACTTGATGTACGCCAAACGTGCTTTCGTCCACTGGTACGTCGGAGAAGGTATGGAAGAAGGAGAATTCTCAGAGGCTCGTGAAGATTTGGCCGCTCTTGAGAAGGATTACGAAGAAGTTGGTGTTGACTCCGTTGAAGGTGAAGGTGAAGAAGAAGGAGAAGAATATTAAACAGTGAAATAAAAACAATGAATTCTAACAAAATACCATCTTGAATTAAAATAAACTTGTTATAAGATCAAAAATTAAAAGTTTATTGACCGATAATTATGTTACTTTTGTTTTTATTGATTGATCGATTGATGTTTAACTCAAGAGCGGCC

>Bpl_scaf_20631-1.16

TCTTGGGTGACGGGTATTCTAACGTCTAGTGAGCACGCTAACAACATCAGTAACGTAAGCCAATCAAGACGAATTCAGTTAAAAGTGGGCGGGGCTATGTGTATTAATGAGATTCAAGCTCTTCCTGTTCGTCAATTTTACAAACATACTCATTGGACTAAATGTAATTTAGATAATGTCAATGTACCACTGAACGGAGCAAGCACTACTGACAAATATTAACATTTATGTTAAAATAACATTGCAATAAGGTCTTTTATTTGATATATGAATCAATACCAATATTGGACCGGCATATTGGACTATTAAACAATATCATGCTGTGTATGGGCGCTTTACAAATGGCTCACATAAAATCTACACTGATTCACTAAAATTGGACACTATGGAGACACTTCTGTGAACAAGTTTTTTTGTTTTGTTTTGTTGTGTATACCATTCAAGAAGAAATGGCTCATGAAATGACTCAGTCTCCTTACCCAGGAAGAGCAATGTATCCACCTTACCATGATTCTGGACAACAAGGGGATAGCATGTTCTCGTATCAACAACATATGCGATCTTTAGGATATCCATTCCACATGAACTCTATGTCTCCTAGCGGATATCAACATCCAGTGGGACAGACATTTTCTATGCCACATTATGAACAAAGTCCTTCACCGCCAAGAGATGATAAGACACAAAAGGATGGTCTACGTCTAAATGGTAAAGGAAAAAAGCTGAGAAAACCAAGATCCATATACTCCAGCCTTCAGCTTCAGCAGCTAAATCGTAGATTCCAGAGAACTCAGTACCTGGCTCTTCCAGAAAGAGCTGAATTGGCAGCATCGCTTGGAATTACTCAAACTCAGGTGAAAATTTGGTTTCAAAACAAAAGATCAAAGTATAAAAAAGTACTAAAGCAAACCGGAAATGGAACTAATCAGGTCATGGATCAAGTGAACGTTTCTGAAACACCACCGATGCAATCTCCAAATCAATCAAAACCGGAAGACCTACATCCAATTGAAAAAAATAGTCCTCATGAACAGACAATTTCTGAACAAAATGGAAATGTGTACGATACTCATCCTTCTATGATGACGGCATCATACTCAAACGTGAATACACCGTCAACTTGGTCGGAAATGAACTCTGACTCCCGCCAGGTGGACATGACTTCCGGTTCCCGCCAATCATCCGCATATATACCAATGACTTCCATGGAGCCGGGGATGACGCACATGTCTCATTATCCTTCCTGGTACGCTCAACAACCAATGCGTCAACATCAATTATGAAATATAAAATAATTATTTATTTTATGATGATCATTATGTGATAAGCAAACAATGTAACATCCACGCATGACCGGAAGTGAAATTTATATATATACTAATTACAAATCATTTTCTTTGTACATTGTATATATGCCATTGACTACTATTTCATGTCAACTGTATGTATGTGGCAAAATCTGTCTTGTTCATGGGAAGACAATTTTGGTAAATGTATGGTTATCGCAATATTGTCATGTCATGAA

>Bpl_scaf_5544-0.0

AAAAAGGGGAACTCCTAGCTCCGGCACTACCCAACATTTTACTATCATCGGTTAATCAACTCATACAAGGAAAACAAAAATGCATTCTCTTATGCTGATTTGTATTGTTGCTGGCGCATGCTGTGTGCAAGGGTTTTTATTGAAACGCAAATTAGATCGAGGAAATGAATGTTTTGCACATATGGCAGGGGAGATAACCGAATTTTATTTCGATAATGAAGAAGCTGGTGAAATGGTTAAAGGCGCCATGAGGGAATGCTTAGAGCAGAAAGACAATAAAGAGAAATGCTTCCTTGATAAAAGAAAGGAATTTGTCGAGGTGTATGGGGAAAACGCGGCAAAAGGTATAGACGCCATACTGGGAGGTCTTGTCCATGTGGGGACAGCTGTTTTGCAACATTGTGGTGGTTTGGAAAAATACGCTTTGAAGATATGCGCAGGACATGTACTTCAGGGATTTAAAGATCATTATGCATGTGATGAAAAATGTGACGTTTTTATTGATCTGTTTGGAGAGAAGGGTGCAGAATGTGAGGAACTGGATATGATAGAGGGATCCAAAGAGGGTAAGGAGGGAAATAAGGAAAGCGGGAAGGGATCCAAAGAAAGCCAGGAGGGATCACAGGAAAGCGATGAGGGATCACAAGAAAGTGAGGCAGAAAGGGAGACCAAAAGAATATTAGGTGAACTACAACGTCTTCGATCTGTTAGAATTGGCAAATATAAAAAGTAAAGATTACCAGTGAACTAAAAACGCTAACAAATATAATATGACGATAACTGACAAAGACATGATTCATTCACTTGTTAAATGTTCAATCAATAGACCTTTTACGCAATCCTGATTTAATTCTAAAATTATACGTTTTTAATACCATTTTTATACGAGTTATTTCCCTTATGTCAGGCTAAAAGTTAGAACAATGTAAGGACAACGAATTTGAGTATAAATTTCTAATAATTTGTAAATAATTGAACCTTGCAAAAAAATCAGAGGATAATTTCATAGGAAGTAGAAACTTCACAAGTTGTTACAACATTAAAATGTCGAACAGATTAGAGGAACAAATATGT

>Bpl_scaf_681-5.9

TTGAATTTAGATTGTAAGTTTATTCTTGATTGTTGGAGGGATCATTGCATACAAAGCATCAGAGATAGAAGAAATCAAAGGTATAGAAAAAACAATAAAAGATTCCCTGAAGGCATTAGCTAAAAATACTGAAAGCTCAGATGATGGAATCCAGGATTTCTCCATATCACAGTTATTGGATAGTATAGGCATGGCCTTTATAATTTCTGGATGTGTACTACTATTTCTCTCATTCTGTGGCTGTTGTGGAGCATGTTACAAGTTCAGGACGTTGATTTTCATCTACGGGTTGATTATTGCAGTCCTCCTACTAGCTGAAGTCATTGTTGTTATTTTGATGTATGCTGTTCCAAGTACAATCCAAGAAAACATTAAGGATGTCCTGTTGGAATCCTTGAAAAAGTTTCAAGGTATGGGAAAATCTGATATAAATACACTGGGATGGATGTTTGTCATGAACGAGATGAAGTGTTGTGGTGTGAATGGATACACGGATTTCTCTACTTATGCTACAACTTGGAATAGAACAATTGGAAGTTCTAGTTCAGAAATAGACGCACCTCTGGTCTGCTGTAAAACGGCACCCACAAATGTATCAGCTATCAACTGTGCTAGAAACGGAACACTTGATATAAATGAGGAGGTATGT

>Bpl_scaf_2971-0.17

TCCTATAAAGGCGGTATAACGATGGTAATAGCTACACATATTTACTTATCAGGGAAATTCAACTTGAAGATATGAAGACCATTGGAGTGTTACTGCTTGTTGTTTTACTGGAGACAGTAACTTGTCGACCAGATCAAGCCCTCATTGAATCCAAAAAGAAAGCAGTAAAGGATTCCAGTCTGGCAGCTCGAGATATGTACTTTTTATGTGACAACGGCAACTGGATTTTTGGTTCATATAGATGTGATGGCGATAACGACTGTGATGATAACTCAGATGAAATAGGATGTGCTGGGTGCGGTTATATTGAATTCAAGTGTCACGATGGCACATGTATACCAGGAAGTTATGCCTGCGACTCTTGGGCCGACTGTAGTCAGGGAGAAGACGACTGTGGATGTGGAGTGTGTGAGACTGACGAATTTAAATGTGACAACGGTAACTGTATTCCTGGTTCATTTAGATGTGATGGCGATAACGACTGTGATGATAACTCAGATGAAATAGGATGTCCTGGGTGTGGTGATAATGAATTCAAGTGTCACGATGGCGCGTGTATACCAGCAAGTTATGCCTGCGACTCTTGGGCCGACTGTAGTCAGGGAGAAGACGACTGTGGATGTGGAGAGTGTGGTCTGAGTGAATTCCAGTGTGGTAATGATAACTGTATACTAGAGGGTTACATATGTGACATGGACAACGATTGTCTAGACAACACTGATGAGCAGGGATGCTCCAATATGGACAAGAAGGCTTTTCTATCAGCACTAGGGTTGAACAAAGCAACAAAAGATAAGAAAACTGCCGGGCTCAAAAGGGGAAAACACATAAGAAAGAACCTGAAGCATAAAAAATCAGAATTGACAACTAAAACAGCTGGG

>Bpl_scaf_33659-0.9

ATGATAACTCCCATGAAATATGATGTGGTAAGTCACGTGACATCAGATGATAACTCCGATGAAATAGGATGTGCTGGGTGCGGTTATAATGAATTCAAGTGTCACGATGGCGCGTGTATACCAGCAAGTTATGCCTGCGATTCTTGGGCCGACTGTAGTCAGGGAGAAGACGACTGTGGATGTGGAGTGTGTGATCTGAGTGAATTCCAGTGTGGTAATGATAACTGTATACCAGAGAGTTACATATGTGACGATGACAACGATTGTCTAGACAGCACTGATGAGCAAGGATGCTCCAATATGGGGTGTGAGACTGACGAATTTACATGTGACAACGGTAACTGTATTCCTGGTTCATATAGATGTGATGGCGATAACGACTGTGATGATAACTCCGATGAAATAGGATGTGCTGGGTGCGGTTATAATGAATTCAAGTGTCACGATGGCGCGTGTATACCAGCAAGTTATGCCTGCGACTCTTGGGCCGACTGTAGTCAGGGAGAAGACGACTGTGGATGTGGTGAGTGTGATCTGAGTGAATTCCAGTGTGGTAATGATAACTGTATACCAGAGAGTTACATATGTGACATGGACAACGATTGTCTAGACAACACCGATGAGCAGGGATGCTCCAATATGGACAAGAGGGCTTTTCTATCAGCACTAGGATTGAACAAAGCAACAAAAGATAAGAAAACTGCCGGGCTCAAAAGGGGAAAACACATAAGAAAGAACCTGAAGCATAAAAAATCAGAATTGACAACTAAAACAGCTGGGAAGAAATCTGATGTCAAGAAAGTAGCTGACAAGAAGAGTCATGTTGATCAAACCATTCAAAAAGTGGAAGAAGCAATAAAATCAAAGATTGAAAAATAAATATCCAGGAATATTATCCCAACAGCCACGAGGGACAAAGTTATCCATGATGCTGAAAAGAATGTCATTTCTATTCTCATTACTTAATATATATTCGCACTTGGACTTTTAACATTCTGTTGTGTTTTAATGATATTTTCAACATTCTGTTGTGTTTCAATGTTATTCTTTAACATTCTGCTGTTTTTCGATGTTATTTTAAAGTATTTTGATTTTATTTAAGGAATATTTGAAATTGTTGAACAGGTTATACGATATAACCTGGTTTGGGTCAGTTTACGCCTAATAAAACGCGAAGCGGTTTATGAAAAGAGAAAACTGACCCAAACCAGGTTATGCAAGTATAACCTGTTCAACAATTACAAATATTCCTTATAATTTAATTTTCACAACAAAATTTAATGATTCCTTGATTGTAATTACGATTATTTCATGTGATTATTTGAATTTGTCGTTGGCGGATGAACACGTTGGTGACGTCACAATACCAACCCAGGTTATCTAGTATAACCAAAATTATCTCCAGGCGGATTAGCCAATCAGATTGAAATATTCAAATCAAATTAAATTATATGTCTCTAAAAGTAAGAGTATTGAGTATACAAATAATCAAACATGTGATGTATATTTCTTAAAACTTAGTAACAATAATATTTTCTGTCTATGTTGCTGTTTTTCTCTAACAATTTCATTGTAATTCATCCTAATTCTATATTTCCAAATAAAAACATTTCTCTGC

>Bpl_scaf_48777-1.5

AGCCATTGCAATTCCACAGTTATTCCAAATTAAACAGTGACATTGTGCAGATATTAATTTATTAATTTAGATGATTTTTGTTTTAAATTATTGTACGAAAAGACACAAGAGTATTTTCAGTTCAAATTGGTCTTCACCGTCTTGGATTTTTGCCATTACGATAAACAAAGTCCAAGGTCAGTTTTCCATATAAGGAGTTACTCCGCCTCTAAAGGTTTCCCATTGGCTAAGGAAATCTAGATTATGTAGATATACTTGAATCACACACGATTGATCTCATTCATTCATAAAACTGCGAAGGAATACGTAACATTGAAAAATATTCTTAAGGATTTGATACCGTCTTAAATTTAACTTACTGTGATTTCATTTCAAATTCAGGCATATTGGAAAATATTAATAAGTTTGGTGTTACTCCGATGGCTACAGTGGATATAGATTTGACAGGTGAACAAGAATTCTTTGATGATCAAATTGTACCTGATCAAGACGATTGTTTCTTACCGCCAAGTGATATAGTTCCACCACCTCATTTCAATACTAAAGAGCTTGACAACTTCTTTGCTACTTTAGACGGCTACCCTCCTGAAAAGAAACAACGCCGTGAGAGTTCATCTCTTGTAGATGAATTCTTCCAAGATGACAAGAACGCGCCAGAACTCCTTAGCGCAAAACAGCGCATGAGCATGTTGTATGGAAACAGAATATCTGACACGTCCTTTTTGAGAACGAGTATTCTTCAAGCTAAATCTCATGCAGCTCAGCAAGATAGACGAGAAAGTTCATCTTTTGTTGATGATTTCTTTGATCAACCCTCGAAAAAAGATAAACATAACAACAACGCAAGCGTAGGAGCTCTTGGAAATAAACCCGAGGGAAGTTATTTCGACATAACTGCTACAATGATGGGAGGAAATGGGGTCGTGCCTTTTCCAAACATGTCTGAACATTGTTTGCAAGAACAATCTAAAACTGTGGATTCGTTATGGGAGGACATTCGAGATAGTATTGACATTGATAATTATTCCGAAACTGATACTAGTAGTCAAAGTGGGAGCGAGTGCGACTTACAACGTGTGAAAATTGAGCAGCCAGACCCCTCATACAAATCGTCGTGCCAATTTCAAAATACAAGGACATCAAGTTTTGAGAACGTGACTATAAAAACGGAGCCAATAAAATCAAGTTGTGCAATGGATTCACCCATGGGTAGTCCAATAAACTCTCGACCCAGCAGTTTACAAGTATCATCATGTCAACAGAACATTGTTCTTGCACAGCCCGGTTCCACAATGGTTCAGTACGGCTCTAATGCTCAATCTCGTATGAGAAAAGTGTTGACTTCATCGTCAAATTCGTACAGTAGTACATCAACGCCACCCCACTGCGTTCCAGTATTTCTACCACCGACTCCTCCCAACTCTCAGCCTGGAAGTCCTAGTCAAGATCAATCTTTCAGAAGAACACCGCCGCCGCCATATCCTGGATTCATCAGACCGCAATCTCACACTCCAATGACATCACTTCCGGTGGCATTTACTCCTATTCCACTACCAACATCGACATTATCAGCGATTGAAAAAAACCGTAAAACCCAGCAGACTCACCCAGGATGTTCAACTATTAAATACAATAGAAAGAATAACCCAGAGTTAGAGAAAAGAAGAATCCATTTCTGTGAATTCCCAGGATGCAGAAAAGCTTACACAAAAAGTTCACACCTAAAGGCTCACCAAAGAATTCATACAGGAGAAAAGCCATATACATGTCACTTCCCAAGTTGTCAATGGAGGTTTGCTCGTTCAGATGAACTGACTCGTCACATACGTAAACATACTGGTGCCAAGCCATTCAAATGTAAAGTCTGTGACAGATGTTTCGCCAGATCAGACCATCTTGCTCTTCACATGAAGAGACATGAACCCAAAAAGTGAACCCCCTGGTTGACTTAGTCGACGGCTTTTTGGAGAAACTAGAGCGAACTAGATTGCGACTCGCATCTTTACAATGCTCATTGTTGTGATTTGACGACAGACTTGTCAGAGAAGTCGATTTTTGTGCAAGACTACGTTTAGTAGATAGATTTGAAAGGGAGGACTACTTTTTTTAGTCTAGACGTTTTACCGTTAAACGTCATTCCATGGCCAGTATTGAGTAGGAAAACATGACGAAAGATTTTGCAGTTGGGAGGTCCTGCTTGGACATTTAGTCGAGACTCGTTTCAACTTTTAGAGTCATTTCAGGAGATGAAGTTGACATTGGTTATTTTTTTATTTTTTGTTGTTGATCTCATGTGATATTGTTATTGTTATTTTTCTTGAGCCACTCGAGTGTTACATTTATTTTATTTGTGTAAACATCGATGTTACCGATTGTGACTGCCTGCACTGGTGCCAAGTTAAAAATATGTAGAGATCACTAGAATCATATCACTTTATATATTGTTTGAAATCCAAAATGAAAAGAAAACATTGACTTGATCCAACATTTATTTGTTAAAATTGCTTGTCGATTCCATTTCATATCGTGCTTTCATATCATACGGGTCCTCCGTTTGGACATGTGATGGTTTGTTTTATTTCCGGTTCCGGTTTGTATGAAATTGACGAGGGCATGCGCAATGATCATGAATAGTCAGTCAGAGTGGGAAATGAATGGAGCGAAAAAGAGGTTGTGTTGGAGAAAGTAAGGTAAACAGAATATGTGGAGATCAGAGCGAGAGATGCTGCTTGTTGTGACTTTATCTTAAATGTAATGCTCAATCTATATCTATAGATTTTATTTTCTATTTTTTGTACATGTTTATTATAATGTGTTGACGTAAAGATAAAAAATTAAAAATTAAAATATATTGTGAA

>Bpl_scaf_22362-6.9

CTGCACTGTGTACTGAAACATCAAAGCTGCTAACATAAAAGTAAGGAAAAAAAAACCTCACAGAAATTAACATGGACGTTGATATGAACATGGAGACGACTTCGTCATCTTATTGCTACAAGAGGTCTATTACGCTTGAGCAGAACTTGAAGGGGGAAGGAGCGGAGTACAGCGAAACACCGTCCAAAAACCTCGTCCAACAGAACATCGAGTTGAGACGGAAACTTGAAGAGGAACATCAAAGTTATAAACGTAAGCTACAGGCATATCAAGACGGACAGCAAAGACAAGCCCAACTTGTACAGAAACTTCAAGCCAAGCTTTTACAATACAAGAAAAAATGCACTGATTATGAAACAAAGATTCACTCACAGTCAATTACACAGTTACAGTCACAACAAGATACTTACCAGAAAGGTTTAGATAGTGAAAGTCGTCTCCGCCAGGAAGCAGAAAGCAACATGGACATGGAAGCAGCTTTGATCAAATTGGAGGAGGAGCAACAAAGGAGTGCTAGCTTGGCCCATGTCAATGCCATGTTACGTGAGCAGTTGGATCAAGCAACAGCAGCCAACCAGTCCCTTACAAATGATATCCACAAGCTGACCAATGATTGGCAGAGAGCTAGAGAAGAATTAGAGGCTAAGGAAGCAGATTGGAGAGAGGAGGAACAGTCTTTCAATGAGTACTTCAGCAATGAACATGGTCGCCTCCTCTCATTGTGGCGGGAAGTTGTTGCTTTCCGTCGCAGTTTCGGTGAGTTGAAGACAGCTACAGAGCGGGACATGTCACATCTACGATCAGATGTCACTAAAACATCCAGGAGCATGCACTCTGCCTGTCTAAACCTGAGTGCTAACCAGAGAAGCTCAGACACACAACTTTCGGTTCTGCTCGATCGTGAGAAACAGGAACGCATGTCTTTGGAGAACCAGCTGAGAGACAAAACCAGGGAGGTAGCAGAACTCCAATCTCGCTATGATACTCATAGTGCAGAACTTAATTCTAAGGTGAACGAATTGACCATGATGAATGAAAAACTCAAGATTCAACTGGAGGAACGTGACAAATCCATCGTTAACCTACAGCGTAACATCAACAATTTTGAAACTCGTCTTGGTGAACAAAGATCATTTGACATCCCAGAAAATGAAGCCTCTCGTCAATACCGTGAAGAGACAGACACCATTCATGAGGCTCTTAGAAACATTGCAGAGGCAGTCATTAATGATGCTGATGAGCTGGATGCCACTGATGGAGCTAGGTCAATGTCACCTTCAAGGAACAGGTCAATGTCACCATCTGCAAGGGCAAGGTCACCCATTCTTAGAAACAGATCAAAATCTCCAATGGCAAGGTCAAGATCACCAGCCTTTGCTGATGCCACTTTCTCTGCTGTTCAAGCTGCATTGAACAAGCGTCAGCTCCAAGTGTCAGAACTCAGAGCCAAACTTATCGCAAGTAAGGATCACAATGGAGCTCTGAGAAAGAACTTAGATGATGTGGAGAACGAGAGGCGAAGACTTGAAATGCAAATCATTAACCTTAAAGAAGACCTTGACCTTTCGAGAAGAGACAAAGATGACACATCAAGAGAGAGAGATAGACTAAAGAACTCTCTCCACTTGACTGGTAATGAAAAATCCCAGTTAGAGAAGGTTAGGTCTGAAATGAATGAACAGATCGATGGACTTCAGTCTGAAAATGAGAAGTTACAGGCAGCTAACACTGAGTTACAGAGAATGAGAGACAACTTGGAGGACGAGAAAGAAGACGTCACTAAAGACAAGGAACGACAACTCAAGGAGAATGATAGATGCCACAGAGTAATTGACCAATTAGAACACAGAGTTAGTAGTATTAAGGAGGAGCTTGTGGCAACAAAAGAGGCTCTCAACAGGGCAACTCTTGACAAGGAGGTTCTAGAACAACAGAAGTCTGAAGTCAGTGATGCTTTGACAAAATCTGAGATTCAAAAGTCTGACCTTGAACTTGAGTTAAACAGAGGCAAGACAGAGGAAGCAGGACTGAGAGATGCATTACATAAGATGCAGCAGTTAAATGAAGGCCTCGGCCAAGACAAGATTGAACTGAACAAGATTATTATTATGTTGGAGAATGAGAAAGCCTCTCTACAGGGAGAGAAATCAGTTCTAGAACAAGAGAGGAGTGGTATCAGAGAAGAACTTGTCCGTGTTGAACAGGAGAAGATGGATTTGGACACAGAGAAAAGGGGTTTGAACCAGACCTTAGAATTGAGTGAGATGACAAGACAACAATTAGAAGAGGAGATTACAGCTCTCCACAGGGAGAAGGGAGAAGTTACTGAACAACTGAACAATATTGCCAGACAGAAACAGGCCTTGGCAGAAGAATTGGTAGCTGTCAGGAAAGAGATTGAGAGAGTAAACAATAATCTAAAACGTATTGCCATCGAGAAAGAAAGATTAACTCAAGAAAAGGGTGAACTGATTGTACAAGTTACCGATACCGAGAGAGAAAATCGTCACCAGAGTGAAGTTATCTCCTCTTTGAAAGCAGATAAGGATTCTTTAGAAAGTGCCCTCTATGAAGTACAGGAACAATTCCGTCAATTGGAAGTCCGTAAAGAACAATTAGAGGGAGAAAATCAAGAATTGATCATCAGAAAGGAAAATCTACAATCTGAAATCAACCGCCTTTGTCAAGAGAAGGATGCTGACAATGAGAAGTTTGACTTCCAGAGAGAAGATCTAAACCGTCGTTTGGCTCAGTTAGAACGTGACATGCAGATGGCCTTGACCCAAGAGAAACAAGCCCATGAAGATGATGTTGATCGTCTCACTAGAGAAAGGGATACACAGAGAGGTGAGTTTGAAGCTAATCGTGAAGAGATGATTCTCCAGTACACAATGGAGAAAGAAGAATCCAACAATAAGTATGACAGAATGAGAGAAGAATTGATGGAGGAACTATCCACTGTACAGAGAGATAGAGATAATTCTATATTGATGGCTGAAAATGATAAACAACAGACAATGTCACTTTTGGAACAAGAAAAGAGCACCTTGGCAGAGAAGAATAACAACCTCACTATGGACTTAGCTAATTCTAACGTAGAGTATGAAAGACTTAAGAGAGAATATTATGCCAGACAGGAACAGGATAGGACTACCATGAATGGTCTTAATGGTGAACTGAAGAACTTACGTAGTCAATATGATGAGACTTGCATGAACCATGAAAAAGAATGTAAAGACCTTACAAACCAGATCAGAGAACTGGAGAGACAGAAGGAGAGTGCTCTTAGAGAAGTCGATGAGCTTAAAACACAGCTTATACTTGTGGAGGAGAGCCGTGATAATATCCGTAGAGAACTGATCGAAGCTAAACGTAGAATCAGAGAGGGAGAGGAAACCAGAGATCTAATGAGAAAAGACATTGTAGAATTGAAACGTAATATCAATGATGAAGTTAGAGAGAAAGATACAGTCAGTAAGACTGCTGAAGATCTTAGAAATACCGTTAAGAGGAACGAGTCAGACAAGATTGAACTGAATAGATCATTGCAGGATCATAGACAGAAATGTGCAGTATTGGAAGAACAAAAGGCAAATGTCCAAAAAGAGGCTGGTGATCTTAGAGCTAGTCTTCGTGAGGTTGAAAAAGCACGTCTGGAAGCACGTCGTGAACTCCAGGAATTACGTCGTCAAATTAAACAGCTGGATAGTGAAAGAAGCAAGCTTGGAAAGGAAGTGACAGACCTCCAGGGCCGAGTGGCACGGGATGAAGAAAAGGAAGAGGAATCCAGAAGGACCTCATTTGACCTCAAACAAAAGGTTGTTGAGACTGAAGCTAGCCGTGAAGCTCTGCGAAAGGAACTGGCAAACTATCAACGTAAGATGGGCGAACTTATGGATGAAAGTCGCATGAAAGAAAAGGACTACCAGATGGCTCTTGAGGACAGTCGCAGAATTGAAAGGAAATTAGATGACCAGAAACGTAATTTAGAGATTCAATTAGAAAACACAAGTGCCGAGAATGAAGAGCTCAAATTGAGGCTAAGCGGAGCTGAAGGAAGAGTTAATGCCCTTGAAGCAACTCTAGCCAGACTAGAGGGCAGTAAACGTGACATTGAGTTTAAACTGAGCAGCATTGTGTCAAGTTTGAGGAGAACAATTGGATTCAGACAGGAAATGCCTAGAGCCCGTAGTCCAGTCAGATCTCGTACACCAAGTCCAAGGCGGTCCAGACCAAACTCTCCAGCTAAAGGTTTTGAAAATACCTATGCCACCACTACAGAAGGAAGAGGCAGTCCTATCCCTAGAACAGGATCTCCAGAGAGAGCAGGTAGTCCTATTAGGGTGTCATCACGAGGCGTGTCACCTGCCAGGTTTGAGATGGCTGCCATTGATGTAGATCCAGAGGCAGTTAGAATGGCTCTCCGTGACTTTGTTCAACAATTGGCCAATGCTGAGAGAGAAAGGGATGATGCTTTAGCTAACACTAAGAGTATGGGAATACAGTATAAAGAATTAGAAGAAGAGAAAGGCAGAGTAGAGAGACGTTTAGAACAGCTTCAGAAATCTCTCGGAGATGTAGAAGAAGATAAACGTGGAATTGATGGACGTCTTGCTAGTGCTCAGACCGCCTTAATGCTCCAAGAGGAGACAATCCGTCACAATGAACGGGAACGTAAAATGATGCAAGACAAGATAAATGCCTTAGAAAGAAGTCTCAATTCTGCTGAGACAGAAAAGAGACAACAACTTGATAAGATAAGCAAAATGAAGGCTAACGAAGGCAGACTTGATGATGACAAACGTAACTTGAGACAGGGACTAGAAGAAGCAGAAAATAGGTGTACAAAATTAGAACTGGCTCGTAGATCTCTGGAGGGTGATTTACAGAGATTCAAATTGTTAATGAATGACAAAGAAACAGAAAACCAGGTCCTGCAGGATAGAGTAGAGACTTTAAACAAGGTGGTTAAGGACTTAGACAGTAAAGCCCAGTCCCTACAGTTGACAATAGACAGATTGTCACTAACTCTAGCTAAGACAGAGGAAGATGGTATACAACAGAAAGACAAGGTTCAGTCATTAAATATGTCACTATCTGATAACAATGCTGCCTTAAACGAGGTCCAAGAACGTATCACACAATTACAAAGAGCTCTGACCAGTAGTGAACACGATCGTAGAGTATTACAGGAGAGACTGGATTCTACAAGACAAGCTTTGAATGAAGCCAAGAAACAGAACTATAACCTACTGGAGCGTGTACAGACCTTACAGAATGAGAATTCTGAAAATGAGGTCAGAAGGGCAGAGATTGAAGGTCAACTTCGTCAAAGTCATAGTATGTTGGTAAAGAGACAAGAAACTGAACAAGAAATGAACCAGACTATACAGAAGCTGAACCAAGATAAACAGAACATGCAGGATCATATCACCGGTCTGTCACGAAATCTCTCAGCCGTAGAAACTCAGAAAACTGAGATGGAAAGAACTTACATTAGACTAGAGAAAGACAAATCTGCTCTCAGGAAAACATTAGATAAGGTTGAACGTGAAAAACTGAAGACAGAAGAAATTGCTAACACTTCATTGATGGAAAAGGGATCTTTGGATAGATCATTGGTCCGTTTGGAGGAAGATAACGTTGATATGCAGAAACAGATTCAGCAGCTACAGGCACAACTAGCAGAGGCAGAACAACAGCATGCTCAGAGGTTGATAGATGTAACCACAAGACATAGAGCTGAAACAGAGATGGAGACAGAGAGACTCCGAACAGCTCAGATTCAAGCAGAAAGAATGCTAGAAACCAGAGAAAGAGCAAACAGAACTAAAATTAAGGAAATGAGAAAACGACAGCAATACATTTCACGCAGTGCTCGTACAGGTGATGAAATCAAAGACATTCGTAGTATGCTGGATTCTTCTCTCTCTAACGTCACTCGGGACCAGTCACTCGATCCACTTCTCTTAGAAACAGAAACCAGGAAGTTGGACGATTCTTTAGAATTCCGTGGAAGTTACAGAACACAACCAAGACGGAAAACCAGTCCAGGACGTTCGCCATTGAAATATTCTGACAGGTTAACATCCACCCCTGCAATGCGACGAACCCAGAGTCCGATAGCACTGCGTAAAAAACTCTTGAAATAA

>Bpl_scaf_55568-3.5

AAATTAAACCTTAATCGTGGGCTTTATTGTGACAACCATATCACCAAATTAGTTTGGCCGGTTAGTTCTTGGCATGGTTTTGGAGACGTTCTGAAGGATAAATCCTACACACCTTATGATGCTGCTATAATCAAAGGCAATAATGAAATCGCTGAATATATCAGATCTAAGGGAGGAGTAACGGGTGGTGATATAGACAATATACTACAAGCCAAGGGTAAAAAGAGAGACGCTAAATCTGCTAAGTCACGGAAAAGTGTACAAACATTAGAAAGTACTCCAGAGGAAGCAGAACCAGAGGATAATAAGAAAGTAGACGAAAAAACAAAAGTTGAACCTGTAGAAGACGACAAAAAAAAGAAAGACAAAGGGAAAAAGAAAGAAAAGGAGGAAAAAGAGCGAGAGAAAGAAAAATTAGTAATCGTTCCGAAAGATAAGCAGAAAAAGAAAGATCAGGAAGAAAAAGGAAAAGAGGATCAGAAAGAAAGTGAAAAGGAAAAGGCAGACGAAGACAAAGAAGGCAAAGGTGAAAAGAAAGATCTCACTGACAAAGAACTCAAAGAACTTCAAGACAGAATTAAAGCTGGAGAAGTTGCTGTAGATGAAAGGGATATAAAACGGAAAGCAATAAAAGATTTAGGAGAAAAACAGAAATTAGATGATGTGGTAGAGGAGGATCAAGAAGAAAAAGAAAAGGAAAAAGACCTTACAGAAGAGGAACAAAAAGATTTGGAAGCAAAAATAAAAGCAGGTCAGAGTGACTTAGACGAAAGGGATAAGAAACAAAAAGAAGGGAAAGAAACAGAAGATCAAAAGAAAAAGGATAAGAACAAGACTGACAAGAAGGAAGGAGAAAAGGACACAAAGGGAAAGAAAAAAGATTTATCTGACAAAGAACAAAAAGCTGGTCAGAAGAAAGTTAAATCAGGTGATAAAACTGAACGAGATAAAAAACAAAAGAAGACAGAGGAGGAAAAGAAAGAAAAAGGTGGGAAGCCTGATTTAACAGACACTGAAGTTGCTGCTATTACTGCTGCAGTAGTTATAGCAACAGACAAAGACAAAAAAGATCAAGATCAAAAAACAAAAGACAGAAAGGATCAGGATCTGAAAACAAAAGATAAAAAAGATCAAGATCAGAAAACAAAAGATCAAAAAGATCAAGATCAGAAAACAAAAGATAAAGATCAATTTGAAGGACTTGCTATTCCTCCATCATCTCCAGATTTACGTGATCCTAAACAAAAAACAAAAGATGGAATAAATCAGGATCAGACATCAAAAGACAAAAAAGACCAAGATCAGAAAATAAAGGATACAAAAGATCAGAAAACAAAAGACCAAGATCTGAAAGCAAAGGACAAAACAGACCAAGATCAGAAAATAAAGGATAAAAAAGACCAAGATCAGAAAACAAAAGATAAAGATCAATTTGAAGGACTTGATATTCCCCCATCATCTCCAGATTTACGAGATCCTAAACAAAAAACAAAAGACAAAAAAGATCAGGATCAGAAATCAAAAGACGAAAAAGACCAAGATCAGAAAGTAAAGGATAGAAAAGACCAAGATCAGAAAACAAAAGACAAAAAAGATCAAGATCAATTCGAAGGACTTGCTATTCCTCCATCATCTCCAGATTTACGCGATCCTAAACAAAAAACAAAAGACAAAAAAGATCAGGATCAGAAATCAAAAGACGAAAAAGACCAAGATCAGAAATTAAAGGATAAAAAAGATCAAGATCAGAAAATAAAGGATAAAAAAGATCAAGATCAGAAAACAAAAGATAAAAAAGACCAGGATCAGTTGTCAAAAGATAAGAAAGATCAGGATCAGACAACAAAACGTAAAGAAGACCAATTCGAAGGAACTGATATTCCTCCATCTTCTCCAGATTTAAGTGATTCCGAGCAAAAAAGAAAAGAAAAGGATAAGGACGATCAAGCTTCAGGTGAAGATACAAGTGATAGAGATCAAGACCAAGAATCGTCAGAAGAAAGAGCAGATGAATTAGATAGTAAGGAAACTAGAAAGAAAAGAGGTTCAGGAAAACAGGATATTAAGGGAAAAAAGGAAGTTCATGTTCAGATTCCTGTAGGTGCTTTTGAGAGATCTCCAGTAGCTACTGCTTCAAATAGATCAAGACGAGATTCACCTGTCAAATACCGAGGTAGTAGATCTAGATCTAAATCTTCTGACGCAGAGGAATTTGAGTTTCCAGAAGGTGCACATGAAAACACAGAAGCTGCAATAGCTGCAGCTGAAGCACGTAGAAAAGCTCGAGAACGTTCAAGATCTAGAAGTGCAGATCGTGAATCAGTCGAGAGACAAACAGAATTAAGAAAATACTCGAAGGTTAATACTGACGGTAGAATTCAGCCTGGTCTTTTGGCAGAAAAAAACGGTAGAAATAGAAGATTTGATGAAAGGGATACTCCTTCACCATTAGATTCCGAAGACGAATTTATCGACGATCCTTATTTCGGACGCAGAATAAAAACTTACGGTGGTGGGTTACGATTAGGAAATAAAAGAATTGACAATAGGGAAAAAATATTACGTGAAAAAGCAGAACTAAAACGAAAGAGACAACTTGAAAAAATGTCACGTTCATCCCAAAGGTTGCCATCTCGTGGAGGTGTTTCTAGAATGTTACCGTTGGGTTATTCTAGATTGGAGACTCCACAAAGACCTATTACTAGAAAATCTAATTCCATACAGGAAAGTGTAAGGAGATATCAAACAGAGAGAACATTCGTTCGTCAACTTCATCAGTTAAAACGTGCTCAGATTTATACTGGACCAATGCATGATATTGTGTTATTTAGTAAGTTAATGGACGTTTATAAAAACAGTTCTGCTAATGACAGTGAACTGGATATTGACTCCCAGGCACTAGATGATTGGGATGGATACTTGCAAGACCAGCTAAGATTTGTCTCCCATTACTACGAAGACGACAGGACAGGACTTAGAGACATAAAAGGAGATGCAAAAGAGTATCATCATCAGTTAGAGAAATCTGCTACAGATGCCAAGAAAACAGTAGAAAGTAGGATAGATCATATCATCAAACAAAATGAAAAAGATGAGACAACAGAACGAGAAGAAAGAGAGAAATTAGATAAGAAGATATCTAATATTGCAGAGAAAACTGGACACATGTTGGATAGTGTTCGTCAACAAGCCAATGAAAGTGTGGCAGGTGCTAGAGATCTCAACAGCAGTTTGGAGGAGGAAATGCGAAGAGAACGCCAAGATAGACTGAAACATTCCAAATATAGTGGTCGAAATGAAAAGGCATGGCTTCTGGAACAGGAAGCAATAGATAATAGGCAGTCCAGAATGGATGAGCAAGACAAAGAAAGAAAAGAGAAACACAAAGAATGGATGCGTAAGAAAGATGAAGAAATGAGACGAAAACTAATGAAAGTCTATGGTCCTACTCCAGTATTTGACCCATTCGTTTCTAAGAAAACATCACTGAAGGCATCTACACCGAAGCCAAACTTTAACCTTAACTCAAGGTCATCTTCAAGACCAAAGTCTATGATGTCTTCAATATCAGGGTCAAGGTCACGACCTCAAGATCAGGAGTTAGAGGTTCATATGACAGAATTTGGAATGAGGAGGGTAAAAAGAGATGAATTTTCCTTTGCCAATTGGACCCCGAAGTATGATGGTATAAATCTACGACCAAGAGCTGGATTAGTCTTTGTTGAACCAACAATATATGATGAAGTGCTTAAAGAAGAACAAATGATCAGGACTGGTAAAACGAGAAACTCAACGAGAAATTTGACTCTAGAAACTGACAATAATATGTTAGAAACTGTAGATTTGTGACACAGGGAGACATAGTATATTATCTTATCCTCATTTCATTTATAAGTAGGTCGTTGTGATACAAAAAATTATTAAACAATATTTGTTTAATATTGTTGTTCCTCATTTTTTAAAAAATAATTTAATGTGATA

>Bpl_scaf_61993-0.4

ATGTTTATCTTTCCTGTAGCTGATGCATGTTTTAGAGAGTCCTTCCCTACAAGTGTTGCTAATGGTGGCATTCTATGTACCTCTGGTAACATAAGTTGCACTGTAACCTGTGATAGTGGATACACCTTCAATGATGGAAATAAAACAAAGATATATGATTGTCATTCAACAAATGTATGGACACCAGCTCTACCTGTTGATACATGTATTAAATATGATGAGCCTTTCTACATAATTAATCTGCAGGTTGCATATAAATCTGAAGGAGTAATTGCATTCTTGTGTGTAAATGGTCATAAACAGATGTTGATCAATAACACACAATCATTTACACAGGAGATATTCAGTACATGTAACTCTCTATCAGTTACTGTTGGAGCAGTAGAAATTACAAAGATGGAGTCTACAACAAGCAGTTTTTCATATATAACAGTTTTTGAAATCAAATTATTGGATCGAACGACAGATGTTGCTCTTAGCCAGATCTGCAGTTTTTTACTCCGTGCTCAAGCAACAACAAAAGCAACTTTTCTCCAGTATGATTCCTCAGTATCTTGTAATTCTGGTTCAACTCAAGTTAACAGAGATCATACCTCTGACATAAACAAGCTGATAAGTCATGGGAATCTGTGTACTGCTGGTGTTAAGAAAGATACAGATGATGGTGAAAAATGTGTGCCTTGTCCACCTGGAACATTTTCATCTGCTGCTGACGATACCTGTACTAAATGTCCAGATGGAGAGTTCCAAGATGGTTTTGGACAAACTACATGTAAATCATGTCCTGCAAACAAGCCTGTACCATCAGCTGATAAGAAAACGTGTGTGGGTGTGTATATTTTATACATTTATGGTATATTAGAAAGTCATGTTTGACAAAACTAGAAAATTTAACAATATACAGTCATACTTACCCTAACAGTCACCTTAAATTAACTGACCTATGGTGGGAAGCTTGAAGTGGGAGGGGTTCCTACATAAATTCAAAGATTAATACCTTAAATTA

>Bpl_scaf_13642-5.8

AAAATGTTGGCGCATGCCCATGGATATCCACTCTCCCAGACTCATTGTATAATGCACCGGTAATGAGATTGTTCATTCATTGATTTCTGAAAAAGCAGGCCGTGTGTATTTTCAATACATGTGCAATGGAAAAATACTTAGCAGAAAAAGAGAAAGCATTGGTTGATGATGACTCGTCATCTCCCTCAAACAGTCAGTCTAGTCTTGATAGTGGAGACGAGAATGAAGAAAGAGGCAATTGGACAGGAAAACTTGACTTTTTCTTGTCTTGTGTGGGATATGCTGTTGGTTTAGGAAATATTTGGAGATTTCCTTATCTATGTTACAAGAGTGGAGGAGGTGCTTTTTTGATACCTTATGTTATTTTCCTTGTATTATGTGGTCTACCATTATTCTTCATGGAGGTATCCTATGGTCAGTTTGCCAGCCTTAGTCCTATCACAGTTTGGAGAATGTCTCCTCTCTTTAAAGGAGTTGGTTATGGTATGGTAATAATCTCAGGAATCGTATGTGTGTACTACAACATCATTATAACGTGGACATTATACTTCCTGTATATGTCGTTCCGGGCAATACTTCCATGGAGTACGTGTTCCAATTCATGGAACTCGGACAACTGTTATTTGCATACAGAAAACAGTACAATGGGAGATAACTCTACATCATTTAAGTGGAATGTGACGGCTTTAAGCAGCATGGCATACAATTTAACAAATGGTGGTAATATGACAGAGAATGAGTTAACAAATGAAACAGCAAAACAAATTACTCCTAGTGAAGAGTTCTGGCAGAGGCATGTTCTACATATGACAGATGGTATAGAGAACATGGGTGGAATCAGATGGGAGCTGTTGATATGTTTAGTGGTAGCATGGGTTCTGGTCTTCTTGTGTTTATGTAAGGGTATCAAATCCTCAGGCAGGGTTGTGTATTTTACTGCAACATTTCCATATTTAGTACTACTGATTTTGTTGATAAGAGGTGTAACTTTACCAGGAGCTTCTGAGGGACTCAAGTTCTATCTTATTCCACAGTGGGAAAAACTAGCTACTTTTCAAGTATGGGGAGATGCTGCAGTACAGATATTCTATTCTGTTGGTATGGCTTGGGGAGGATTAATTACTATGGCTAGCTACAATAAATTTCACAACAATGTATACAGAGATGCTATGATCGTCCCTTTGATAAACTGTGGAACCAGTATATTTGCTGGATTAGTTATATTTTCTATCCTTGGATTCATGGCTCATGAGACGGGAGATAAAATAGAAAATGTTGTTACACAAGGTCCAGGATTAGTTTTTGTAGCTTACCCAGAAGCTGTTGCTAAGTTACCGATCTCACCTTTGTGGGCGGTACTTTTCTTTTTAATGTTGTTAACGATTGGATTGGACAGCCAGTTTGGGATGTTTGAAACAATGACTAGTGCATTCATAGATGAATTTCCAGAATATCTTAGAAATAAGAAAGTTCTATTCACAGCAGGAATGTGCTTTGTGGAATTTTTGTTGGGATTACCATGCATTTTTGAGGGAGGAATATACTGGCTACAGATCATGGATTGGTATTGTTCTACGTTCTCTCTGATGTTGTTGTCCTTAACAGAATGTATTGTCGTGGGATGGATTTATGGTGCTGACAGATTCTACATGGATATTGAACTGATGATAGGATACAAACCAAATGTTTGGTGGAAGATTTGTTGGAAATACATCACTCCAGCTGTTATAGCTTTTGTATGGCTGTTCAGTGTTACCCAGTTAAAGGCTGTAACCTATGGCGATTATGAATACCCTACATGGGCTATTGTATTAGGATGGATTCTAGGACTGGTGTCATTAGCCCCACTACCTATCTGTATGATAACAGCTATCTATAATTGTAATGAAGGAACTCTACTAGAGAGAGTGAAGAAGTTACTACGTCCTGATGAGAAATGGGGACCAGCCGTTGATAGATATCGTCAAGAATATGAAGCTTCAAAAGATTCAATGTCCTCGAATAACGCTATATGTATATCATACAGAACTGATATAGTTAGTCTGGCTTCTATGAAAGATACACCTGAGGCAGAAAATCTTGTTATATAATACCATATCTATAGGGCAAAGTAACTTATTACAATTAGTATGAAGGACCAGAGACAGTTGTTCATTGTAGGAATATGTTTATGATGATGCTTGTACTGGTAAATTATTACAATTAGTATGAAAGGTCAGAGACAGTTGTTCATTATAAGAATATGTTTATGATGACGCTGGTACTGGTACTTAAGTTTTCAAGGTATGCAATGTATATTGAAGGCTAGGTATAGAGGGTACGATAATGATGACCAAAAGATATCTATATAGGTTGCCTATACTGGTGTTGAAAAATGTGAGATGTTAATTGTCCTGAAAAAATTTCAAGACACATCAGCTTACAAACAGATTAAATGATATTTTTTTGAGAGTTGAAAAAAAATATTACTTTACTTCATATCAAGGTATTATGCATCATAAAACACCTATACAAATCTTTTATTCTACACCTTTGCTAAATTGCAAACCTGGAGATAAATTATTTTAATTTGATTCCTGACAAAACATTAATTTTGTGACACATGTTTTTTTCTTATCCATTCAACACATTTATTTGAAATTTAAGAAAAAAATATATGCCTTTTTGCTTTTATGAATTATTAATTTAAATTATGATATGTTCTTTCAACATATAAGTAGAAATTATATAGCTACGTTAATGTGTGTCATCCAAATATAAATGTTTAACCTTTTATTAATAGTTGAGAAAATTTATAGCAATTATACAAAATTTTAATGTCCTGTCTTGTTTTCCTACAAACAGAGTTCTGAATCTTGAATAGGAATGTTATTTTAAAAATTATTTCTCATATCACACCCCCAAGATTTAAATATAGGCATACAAGCTTTATGAATAGGAATGTCATTGGAAAAAAAATCACATATCATATCCTATATCAGCCCCCAAGATTTAGATATAGTCATAAGAGCATTATTTCTTGGGCTGATAATGGTTATCAGCCCCAAGATTTAAATAAGAGCATGGGCTGATATTGGTTTAGAGTTAATGTTGCTGCAAGCTGTTATACACAATTGTTTGGTCAGCTATTAGAGAATAGTGTTGTTGGACTTTAGGAGAAATATATTGTTTAAATACTCGGTACACATATTCTGTGATAGAATTATTATAATGTGTTCATTTATATCATTATGCTGTGTAATGTAATTATAATCAATTGTTTTAATACAAAAGCATTCATATCAGTTTAACGATGTTTTAATTGATATCATTTAAGATCACTTATATTACATTTTTAATTATTTCACATGCCAAATTAAGTTTTATTTTGATATAAGTAGAATGTAACAAATTTATTATTATTTTTTTCATGATGATAACTGTTATAACATTTTTAGTTATTTTTTGAAAACATTGTATTTTGTTATCATTTAAGCAAATGTTATAGGCAGCCTACCGCATAATGAATGCGTTTCTTTTGTGACCCGTTTCACGAAGAGTCACGTGACCTGCTGCCTTCACATTCATCAAATCAAAACATATTTTTTTAGTCATACCCAGTAAGTTTTATCCCTGTGTCAATTCTTTTGAAACTGATTTAGCAACGATCCTAGGATGCTTGTCTTTCAAAATTGCATTTTACAACCCTACCACCAATCAAGATTGACCACCATTGCTCAAACTAGAAATTTTAATTTTGCTTGTATTTATCATGGAGATGAAAGAATTCATATATAACTGGATATTTGCTAGTAAGAAAGCAAGTTTTTAGCAGAATGTATTTTCAACAGATATCCATATTACAACAGCTTGCTTGAATAGTCCATCTTGAGCAAAAAAAAATCCCATTTTGTGTTTTATTTTTGACAGTGCCATCTATATAATAAATGGATAATATTGTAATGAAATTAAAAACTTCAATATTTTTAAATTTTGAATAAACTGAAATTGTCTTCGTGTGATAGAGTTGTTTATTTATATCCTACATTTTGTGTAAATGTTTGTGCATTTGACAAATGGTTTTGTCTATTTTGCAGATAAAATCTATTTTCTCTTAAATTCTTATTAAGCTCCTTTAAAAAAAAACAAATTATGTATTAATAGATTTTGTTGTTATTTTATTTTTAAGCTCAACCTATTATTGTAATAAAAAAAAATCTCAGATGAAATCGGTAACTATTTCTATTTATTTACATTTAAGGTATTTATCTGTACGGCAAGCTAGTAATAATGTTATATTGTGTGCTTCCCATCCATTTAAGTTGAAGCCTAGTCTAATTTTAGAACAATCCACATTTTTATATGATATGTTTTTTTTATTACGTTTTACTCATGTAATTAAACAGAATCTCGTTGTAAGTTTTGTTGAATGTAGATTTTAGTTCAGGAGATACAAAAGAACATAATTTTTATTAAAATGCAACTTTTAATAATGATTAGCTGTGATTGTGTTTAACATGGAAAGAAAATATGTTTGATTTCATAAAATATTAGGGAGACGTTGAGGTCGTGATCATATGGTAGTTGGACTTGCAACTACTACCTATATATGCAATCAGTGCCTATCACCACTAAAGTTATGAGTTCGAACCCCGTTCACGATGAGGTGTACTGGATACAACATTCTGTGATCAAGTTTGTCAGTGACTTGCGGCAGGTCAGTGGTTTTCTCCGGGCACTCTTTAAGTTTCCTCCACCAATAAAACTGACCGACTGCCACGATTTAACTGAAATGTTGTTGACAGTTGCGTTAAACACCATAAGCCTAAATCAAATAATACAGTTCTATTATTATAGCTTTAAGTAAACAAGATATTTATTTTACATACATGACTGTATTACAGTAACAATAATAAAATCATGAGAAAGAAAATCTTATTTGTTGAAAGTAAAAAAATAAAAAATATTTCATAGCCAAATGTTTCAAAATGGAAAAAAAATAGCTTCTGTGTTTTAGCAGGAATTTTATTGCAAATATTTTTTACTGATGAAATCCTTGATATTTTCAAATGTATACTTTCTGTTTTCTGACATGATTATATAGATCAAATACTGATAAGGGTGTTGTTAAATACAAATCAATCATAGTTATGTCTTTTTAATCTGTTCTTGTATGCTATTGTTACAGCATTGGTATGGAGATTTATTATGTTCGTAATTACTCAAATGTATTGTGGACCATTCTGATGAATTAATAGTAGACCAATTATACATTTTTATCCTGTAATTTTACTGGGACTATATTTCTGTTGATTTTACAGGGACAATTTTTCTGTTAATTTTACTGGGACTATTATGCTGTTAATTCTACTGGGACTATTTTCCTGTAATTCTACTGGGACTATTTTCCTGTAATTTTACTGTGACTATTTTTCTGTTAATTTTACTGAGACTATCGTTCTGTTAATTTTACTGGAACTATTTTCATGTAATTTTACGCAGACTATTTTTCTGTTTCTTGTCTTAACTTGTAGAGGAACAAATTTCTTTACTTGTAAACATTTTTAGTTTGCAATCTTTTATGTGTTTCAGTGAAATAAAAACATTGTTTTATTGTATATGGTTGTTTGATTTTTCCACCCCAGAAAACCAGAGAAATGCAGGATTTTATTTCTCTCTTTTAAGAAGCCATAGAACATTTTATTGTAGAAACAATTTCAATTCCCCTTTTAGAATCATAGACCTGCAACATTTGATTTCCTTCGAAGAGAAATTGAAAATCCCACTGTAACTGGGGCTTTCGTGGTCGTGTGCAGTCAGTTGTGAATTCGAATCCCGCTCATGGAGAGGTGTACTCGATACCACATTATGTAATACTTCTGAAAGGGTACTCGGGTTTCCTCCACCTATAAAACTGACTGTCAAAATATATCGGAAATATTGTTGAAAGTGGTGTTAAACGCAATGACCCTAACCCTTTGTAATAGTAGAGAATTACACCATTTGACATCCCTATATAAGAACCCATTGCAATTCTCCTTTGAAGAACAAGGGACTACAGCACTTGACTAAGCTTTGTAAGAATGCTTTGTGATTCTGTTTGGAAAAAGCAGAGAAATGCAACGGTTTACAAAAATCTTAAAAAAGACACTGTTATTATCCTTGTAAGAACAGAGAACTGCAACATATGACTTCCCTTGCTTATATAAGCCATTTAGAATATTTCAATATGAGTTCATAGAGCTTTTAGTTTACGTAGATCAGAGATGTATGTGAGGTATTGTCGTCAATATGTGTCCGTCATAATTTGTGTCTGGTATATTTTTTTATATTACAATCTTATTGGTAATCCCTAACCGCATTGGTGGTGTAATTGTTAGCGTGCTCGCCTTGAGTGCAACAGATAATGGGTTTGAGCACCTATCAGGTTAAACCAAGGACTATAAAATTTGTATTCGTTGTTTCTCCTTTCAGCACGCAGTATTAAGGAGAAAGAGCTCGGAATCGGGATAATGTGTCCGAGTAGGGCGACATGTCTATCCGGACTGTTGTTTCAGTGAGCTACACTAGTAAAATGCAACTTAAGTTGGTTTG

>Bpl_scaf_16646-2.15

ATGGGAAAGAAGAATGCAGCAAAAAATGCTGGCAGTTCCCCACAGAAACGCAAGGCAGTTGAAGTGGAAGAAGACAGTGATGATGAGGATGATTCTGATGACAGTGATGAAGAACCCGTAACCAAAGTTCCACAGAAAAATGCAAAGAAGGCCAAGGTTGCTCCACCTCCTGATGAAGAAAGTGATGATGAATCTGATGAAGAAGATGAACCAATGAAGACAGTTCAAGCACCAGCAGAAGAAAGCGATGATGATGATGATGACTCGGACGAGGAAGAAGAGCAAACAAAAGTTACACCAAAAGCTAAAGTAGTGAAAGCTACTGCTCCAGCAGAGGACAGTGATGATGACGATGAAGACGAGGAAGATGATGAGGAGGATGAAGAGGACAGTGATGAGGACTCCGATGATGAAGAAAAGCCAGCAGAAAATGGAAAGAGGAAGAAAGATAAACAGAAGAACAAAGAATCTAAGAAACAAAAAACTGGAGAGGATGTATGTGTATTTGTTGGTAATCTTCCAAAAGAAATGGATGAAGAAAAACTGAAAAAAATGTTTGTAAAGAAGGGAGTAGCTGTAAAGGAAGTTCGTAAACCAGCAAAGAAAAGATTTGGATATGTTGACCTTGAAAGTGAAGATGACCTTGAAAAAGCATTGTCACTTAATGGAAAGAAACTTGGAGATGCTGAATTGAAAGTAGAAAGGGCCAAGTCAAAGTTTGATGGTAGTTCTCCAGATGCACCAAAAGATAAGAAAACTGATGCCAAACAAAGTTCTGGAGGAGGAAAAGATGATGCAACATTGTTTGTCAAGAACTTGTCAGAGGACACTGATGAAGAAAGTTTAAAGAAATTCTTCCCTGATAGTGTAGAAATAAGAATGCCAAGAAAACCAGATGACACTCACAAAGGATTTGCTTACGTTGTTTTCAATGAAGCATCAGAAGTTGATTCGGCTCTTGAAAACAAACAAGGATCAGATTTGGATGGAAATGCTTTATACCTTGACAAAGCTGGATCCAAGAAAACATTCCAGAGCCCGGGGGGTGGTAGAGGATCGTCACAAGGAGAGGCAGGAAAAACTAAAGTTCTTTTTGTAAAGAACCTTTCATTTGACACTGATGAAAATGGCTTGAAGAATGCATTTGATGGAGCTACTTCTGCTAGAATAGCCAAATTTCCAGATACACAGAAACCTAAAGGGTTTGGATTTGTGGAATTTAACACTGCAGATGAAGCACAGGAAGCTTATAAAGCCATGAAAGGACAAGAAATTGATGGCAGGCAAATCTTTGTGGACTTTGCTGGAGAGAGAGGAAGTGGAGGTGATCGAAGGGGTGGTTTTAGAGGAGGACGAGGTGGTGGTGGTGGTAGAGGGGGATTCAGAGGAGGTCGTGGTGGATTTGGTGATAGAGGAGGATTTAGAGGTGGTCGTGGTGGATTTGGAGATAGAGGAGGCCGAGGACGTGGAGGTTTCAGAGGCAACAGAGGTGGAGTACAGTCCTATCAAGGAAAGAAGAAGACATTTGATGACTAAATAACTACATGAACATTTATTTTTTATATAAAGATGACGTTTGTATGATCTGAAGAAGAGAGAATGCCATTTAAATAAGATTGTGATATCCAAGTTTTAAAAGAACGTTGTAAAGCTAGTCATACATGTTTAAATTGGCAGTTTGGTAGTAGAAAAGTTACTAATGTACAAATAAGTGTTCTTTAAAGTTAGACTGCAGTAAATATGTCTATTGTAAAAACCAAATACTCTCTGCACCTTGTTTGATAGATTGATCATGTAGCAATATTTAGAATTGTCGTGTAGAAAACAAGGTTTAGGAATTGTTTTTGTACATTTTATTCAAATTTTATATTGCACATAATTAAACCCCTTTTATATGTTGAAAATCAAACTTAAATATTGATTACATTGGATTGCATTAATGAAATGGATGTGGAAAAAAAAAAAAAAAGACAATTTTAGAAATGGTTCAAATAGAAATGAATTTAATTAATATATAGATGAAATGCTGCTAAAGTTTGGCATTAAGCACCAACCTATCAATCAGTATTGTCAATTGATGACAAAAAAGGATCAAACACAAGTTTTTTCAACTTTGGGATTGGCTTTTCCATAATTCTTAATTAAAGGAAAACGCCATGGTCTTGACAAGTTAGCGTGAAAATTTAAATGACATTAAATAAAAGATATTTGTGTTGATGAATGTTAAAAATAGGCAAAAGGCATAATCCATTCTTGGATTATACCTGCAATTAAAGCTCTTTCGAAACAGCCATGACGTCATGCAATCAGTATCGTATATTATAGCAGCATTGCTCAATTTAAAAATTGTGTAAATCGTAAAATAACTTTTGATATTTACAAAATCATGACTTCATGTCAAGCATTTTATTGCACTAGCGAAATGGGAAAGTGTGAGAAGAATTTTTTTTTTTTTGGTAATTACTGATCAGGACTACCCTAGTTAACTAGTTAACACATGGACCAGTGACCACGGGAAAGTACGACAGCATTGGCGACAAAATAAATACATTTATTTTGTCTTTCAAAATAAATGCAGTTTGGAAGATGAACACAACAATGTGAAATAAAAATGCCAATAGTCTGAATTATCTCCATTCAATTTTGTAGATAATTCATTGGTCAACATTTCAGTATGCTATTATAAGAAAAAGTTCTTTGATTTGTATGAAAAATATATAAAACTGTTTAAAGCTTTTCTTGTTGTCTCGTTTAAATTGTATGTATGATGTGTTCTGTTTATGAAGTCATTATAAATCAATTGTATTTTATCATTGTTGTCGTTTGTTAAGTTTTCATCACATCATTTTAATCATACTAGCAAACAATTGAAAAATAAATATTGAAATCCCTTA

>Bpl_scaf_46838-6.40

GTTTCAGATAACAACGGTGTTTTATAACAGGAAGAATTTTGTAAAGGTAGTTTTTTTCTGAATGAATAAGTCACATGCGGCAAGCTAGGTCATGTGATTTCATTTTCAACAGGTACATTAATATATGAAGTTATAAATTACCTGGTATTTGTTAGTCTTTACAGAAGAGAGAATAACCAAGCCTAACCATGCTTCGATTTTGTTTATTGTTGATTTTGGTTGTGACAGTATACTCAGCTCCTCAGAATCAGGACCAGAATTCCTTTAGTCGTTTTTTCGATATCAAGCCATCTTCAGTGAAGTTGACAGTAACAGTTCCTAAGAGGCAGGCCATCACTTCATTTGCGCCATATGATAAATCATGGGAAAGATTCAAACTTGTTCACAGTAAATCATATCATTCCATACAAGAAGAAATATATCGTAGAAATGTTTTTAAGAAGAATGCCTTAGTGATAGAGGAACACAATAAAGAGTACAATCTTGGTAAAAAGTCATACACTCTTGGAATCAACCAGTTTGCTGACTTGGGACAATGTGGATCATCTTGGGCTTTTAGTGCAACTGGTGTCTTGGAAGGACAACTTTTCCAGGCAACAGGGAAACTTATTTCTCTGAGTGAACAACAATTGGTGGACTGCTCTAGTGCCTATGGCAACGAAGGGTGTAATGGAGGATTAGTGGATAATGCAATTGAATATATTGAAGCCGTGGGAGGAATAGAATCAGAATCAGATTATCCATACACAGCCGAGCAAGGGGACTGCAAGTTTGATGTGACTAAGGTGGTAACAGGATGCATAGTTGGACAGAAAGCTGGGGAGACCAAGGTTACATCAAGATGTCTAGAAATAAACAAAATCAATGTGGTATAGCAACACGGGCCAGTTGTCCTCTTGTTTAAAAAGTAATTATTGGACCTTTCCGAACAACTCCAGGTCACAGAAGTAATATAATGGAACACTGTTACTAAAAGATTTTTGAAAACTAAGCAGTTTTGGTGGAAATACAACCATTTTTTTTTAATCTATTTTTTGAAAGGAGAGGTTGATCTAAAAGAACAATGTTCTATATTTTTGTTAAATTAATTGTTTTTACATTTGATGTTTGAAATCTCAATATTAAAACTTGTTTTCTTTAATTTAATT

>Bpl_scaf_35916-0.6

TAAACTCTTGGTATGTCTGCTGTTTTGTTTAACTTCAGATCATTAGTCGAATGATTTTTCCCCCAATTTGAGACCTGGATTCAAATGGGACAAGAGGGACAACAACGGGAGAACAAGGAAGCTGTTTATACTTGGAGCGAAGTAAAATTACACACCAAGAAGGATGATAAATGGCTCGTCGTGGATGGACAAGTCTACAACATTACAAACTGGGTCGGTAAACATCCTGGTGGAAACAGAGTAATCAGCCATTATGCCGGTCAAGATGCTACGGATGCTATTAGAGCTTTTCACAACGACTTAGATAAACTAAAGAAATATTTAAAGCCACTCCATGTCGGTGCTGTACAAGATATACAGAACAGAAGTATAGATGAAGACTTTAGACAATTAAGATCAACAGCAGAAAAAATGGGCCTGTTCAAGCCAAGCTTTCTTTTTTATGCTGTATCTCTAACTCACATTATTGGCTTAGAAGTGTTAGCTTATCTAATACTGTATTATTTTGGAACCGGATGGTTGCCGTTTATAATATCTATACTTTTAGTTGCCGCTATGCAGGTCCAGGTTTGTTATTTATCTCATGACTTTGGTCACCTATCAGTCTTTCACAGCAGGACATGGGACCATTTCTTTCATTACTCAACATTAGGATTCCTAAAGGGGATTTCTCCAGCTTGGTGGAATCAAATACACTATCAACACCATGCAAAACCCAATGTGCTGGATAAAGATCCTGATGTACGATTGGACAAGTTGTTTGTCGTAGGCGATGTGATGCCAGTAGAGGTGGCCAAATCCAAAAAGCATGTTTCTGTCCCCTATAATCATCAACACAAGTATTTCTTTATAGTTGGGCCACCTTTATTGTTTCCAGTAGTTTTCTTCATAACAAGTATGAGCTATATTTTCAAGAAAAAGTTATGGCTGGATGGAGTTGTGACGTTCCTCTACTTTTTCAAGTTTTTCTTCTTTATGGTACCAATGGTTGGATGGGGATGGGCCATCTTCTACTACGAAGTTTTCAAGTTAATAGAGTCCACTTGGTTTACATGGGTTGCACAATCTAACCATATACCTATGAATATAGATCATGATGCTGAGAAGCCTTGGTTGAAACTTCAATTAGCAGCCACTTGTGATGTTGAACAATCATTCTTCAACGACTGGTTTACAGGGCATCTAAACTTCCAGATAGAGCACCACTTGTTCCCAACGATGCCTCGTCATAACTTATACAAGATCCAACCTTTGACAAAATCTCTATGCAAAAAACACGATATTCCGTTCATTATAAAGCCCCTGTTGACATCTTTTATAGATATAGTCAAGAGTCTCAAGCACTCTGGAGATTTGTGGAATCTTGCATACCACGCCCACCATTTGTCATGATGACTTGTCTATTAATTAGACTTATATCATGGTCTCCACACACTAGTGCACACGTGGATGACCAAATAAATGTACTCTATTTTTTCTAATGTATTTTAACACAATGCTTAATTTAACATTAGTATTGAGACTTTTTGTTGTTGTTTCTTAATGAACCAGTCGTTTAAACAGGAAGCCATTTTGTTTGTTTGGTCGTTTTATTGATCTTTATCCGCCATCCACAAACATCTAACAACATCCTATATCGGTTCACATCTTGTAATCTTTCCTTCACATTCGGAAATTCTCTACCTCCTTCATAAAACCGGTATGCTATTCAATGGCTACAATTATGTAAAGTTTGACACTGATTGGTTGCTTGAAAGGTAAATCTCGATATGGAATAATGGAATAGTGGATCACAAGCTTATAAATAAAGAAAGATATACCAAACATTTAATTTGAATTTGAAAGTAGGATGAACTATCAGACACATACTAAAAACGGAAGTAAAAATGAATGAACCATCAGACACATACTAAAAACGGAAGTAAAAATGAATGAGCCATCAGACACATACTAAAAACGGAAGTAAAAATGAATGAACCATCAGACACATACTAAAAACGGAAGTAAAAATGAATGAGCCATCAGACACATACTTAAAACGGAAGTAAAAATGAATGAACCATCAGACACATACTAAAAACGGAAGTAAAAATGAATGAACCACCAGACACATACTAAAAACGGAAGTTAAGAAGGAGCCTTAATTGCTCTTGTTTTTCTTCAACAGTTCAGTCACCACGATGTTTTCAATGATATATGCATAGTGCATACTTAAATGTTGGTATATATGTTGGTATAAATGTTGGTATATATGTTG
